# Sensory Neurons Innervate Peripheral Lymph Nodes and Locally Regulate Gene Expression in Postsynaptic Endothelium, Stromal Cells, and Innate Leukocytes

**DOI:** 10.1101/833509

**Authors:** Siyi Huang, Carly G. K. Ziegler, John Austin, Najat Mannoun, Marko Vukovic, Jose Ordovas-Montanes, Alex K. Shalek, Ulrich H. von Andrian

## Abstract

Immune responses within barrier tissues are regulated, in part, by nociceptors, specialized peripheral sensory neurons that detect noxious stimuli. Previous work has shown that nociceptor ablation not only alters local responses to immune challenge at peripheral sites, but also within draining lymph nodes (LNs). The mechanisms and significance of nociceptor-dependent modulation of LN function are unknown. Indeed, although sympathetic innervation of LNs is well documented, it has been unclear whether the LN parenchyma itself is innervated by sensory neurons. Here, using a combination of high-resolution imaging, retrograde viral tracing, single-cell transcriptomics (scRNA-seq), and optogenetics, we identified and functionally tested a sensory neuro-immune circuit that is preferentially located in the outermost cortex of skin-draining LNs. Transcriptomic profiling revealed that there are at least four discrete subsets of sensory neurons that innervate LNs with a predominance of peptidergic nociceptors, and an innervation pattern that is distinct from that in the surrounding skin. To uncover potential LN-resident communication partners for LN-innervating sensory neurons, we employed scRNA-seq to generate a draft atlas of all murine LN cells and, based on receptor-ligand expression patterns, nominated candidate target populations among stromal and immune cells. Using selective optogenetic stimulation of LN-innervating sensory axons, we directly experimentally tested our inferred connections. Acute neuronal activation triggered rapid transcriptional changes preferentially within our top-ranked putative interacting partners, principally endothelium and other nodal stroma cells, as well as several innate leukocyte populations. Thus, LNs are monitored by a unique population of sensory neurons that possesses immunomodulatory potential.

## INTRODUCTION

The nervous and immune systems have been viewed traditionally as functionally and anatomically distinct entities even though they share a critical common task – to detect and respond to internal and external threats that may be physical, chemical or biological in nature. The nervous system has a direct role in the avoidance of potentially injurious or infectious situations, while the immune system provides key resistance and repair mechanisms. In light of these functional synergies, it should be no surprise that the two systems interact and modulate each other. Indeed, neuro-immune interactions have been intensively investigated in a number of settings, especially in the central nervous system (CNS) (Klein et al., 2017; Norris and Kipnis, 2019; Prinz and Priller, 2017). Moreover, several recent studies have uncovered evidence for bidirectional communication between the peripheral nervous system (PNS) and the immune system in the spleen, bone marrow, intestine, airways, lungs and skin (Baral et al., 2019; Foster et al., 2017; Godinho-Silva et al., 2019; McMahon et al., 2015; Ordovas-Montanes et al., 2015; Veiga-Fernandes and Mucida, 2016). However, the mechanisms and consequences of these multi-facetted interactions are still poorly understood.

The PNS represents the somatic and autonomic neural pathways that directly interface with the CNS and all peripheral tissues. The sensory component of the PNS – principally the autonomic sensory neurons in vagal ganglia and somatosensory neurons in dorsal root ganglia (DRGs) – provide the CNS with information from the periphery. Motor commands are relayed from the CNS to skeletal muscles and visceral organs, and executed by spinal motor neurons and sympathetic/parasympathetic neurons, respectively. The autonomic motor neuron (i.e., sympathetic/parasympathetic neurons), have long been recognized for their impact on immunity (Chavan et al., 2017; Godinho-Silva et al., 2019), whereas the role of the somatosensory PNS as a critical immunomodulatory component has been appreciated only recently.

Being pseudounipolar, somatosensory neurons in DRGs each send out a bifurcating axon to directly innervate both peripheral tissues and central targets in the spinal cord or the brainstem. Traditional structural and functional characterization of sensory neurons, and more recently single-cell RNA-seq (scRNA-seq)-based molecular profiling, have revealed a high degree of heterogeneity among somatosensory neurons (Kupari et al., 2019; Usoskin et al., 2015; Wood et al., 2018). In particular, one major subset among sensory neurons, nociceptors, has been increasingly appreciated for its diversity, as well as its potential for sensing and modulating immune and inflammatory processes (Baral et al., 2019; Kupari et al., 2019; McMahon et al., 2015; Ordovas-Montanes et al., 2015; Usoskin et al., 2015; Wood et al., 2018). In many cases, nociceptor modulation of immunity involves bioactive neuropeptides, such as calcitonin gene-related peptide (CGRP) and substance P, which are released from activated peripheral terminals of nociceptors and act on various immune and stromal cells that express the corresponding neuropeptide receptors (Assas et al., 2014; Baral et al., 2019; Suvas, 2017).

Peptidergic neurons of putative sensory origin innervate most, if not all, secondary lymphoid organs and barrier tissues, but the density and pattern of innervation, as well as the targeted cell types, are notably different between tissues (Belvisi, 2002; Brierley et al., 2004; Felten et al., 1985; Fink and Weihe, 1988; Oaklander and Siegel, 2005).

This widespread distribution of peptidergic innervation therefore raises the intriguing possibility that sensory neurons targeting different peripheral sites collectively contribute to immune responses by engaging in local tissue-specific sensory neuro-immune circuits. Indeed, recent work has uncovered a key role for nociceptors in shaping immune responses in major barrier tissues, as evidenced by animal models of asthma, colitis, psoriasis, and infectious disease (Baral et al., 2019; Foster et al., 2017; McMahon et al., 2015; Ordovas-Montanes et al., 2015).

To date, the immunomodulatory impact of nociceptors has been documented primarily in the context of innate immune responses, whereby both pro- and anti-inflammatory activities were observed. For instance, nociceptors in the skin can sense invading bacteria and attenuate inflammatory responses against bacterial infections (Chiu et al., 2013). Conversely, upon topical exposure to imiquimod, a TLR7 agonist, dermal dendritic cells (DCs) require signals from cutaneous nociceptors to produce interleukin (IL)-23, a pro-inflammatory cytokine that drives psoriasiform dermatitis (Riol-Blanco et al., 2014). Interestingly, both studies reported that nociceptors also impact regional lymph nodes (LNs): systemic nociceptor ablation prior to a bacterial infection caused a profound increase in both lymphoid and myeloid LN cellularity (Chiu et al., 2013), whereas, upon imiquimod exposure, the absence of nociceptors reduced LN cellularity (Riol-Blanco et al., 2014). In both settings, nociceptors had been globally ablated, so it is unclear whether the observed effects were due to a loss of sensory innervation in the LNs themselves or in the surrounding non-lymphoid tissues.

LNs act as sentinel organs that collect, filter and monitor the constant flow of interstitial fluid (lymph) that is drained from peripheral tissue by afferent lymph vessels. LNs are the principal initiation sites for adaptive immune responses and required for the maintenance of peripheral tolerance to self-antigens as illustrated by mouse models and humans with defective LN organogenesis, as well as surgical models of LN resection (Buettner and Bode, 2012; Karrer et al., 1997; Lakkis et al., 2000; Mooster et al., 2015; Zhou et al., 2003). The generation of local adaptive immune responses against immunogens critically depends on bidirectional information flow between the peripheral site of initial immune challenge and the draining LNs, where antigen acquisition/presentation and subsequent lymphocyte differentiation and maturation are orchestrated. This process relies on the directed transport of antigen-presenting cells, cytokines, and antigens via afferent lymph vessels from the periphery, and selective recruitment of vast numbers of naïve and memory lymphocytes from the blood via high endothelial venules (HEVs). Within LNs, specialized endothelial and stromal cells delineate the lymph conduits and organize the extravascular space into discrete anatomic domains. In addition, a variety of myeloid leukocytes cooperate to support efficient antigen encounters by B and T cells to elicit an appropriate immunogenic or tolerogenic response.

Previous studies of LNs in several mammalian species have shown that LNs are innervated by both noradrenergic and peptidergic neurons (Felten et al., 1985; Fink and Weihe, 1988). While there is general consensus on the sympathetic origin of the noradrenergic fibers (Bellinger et al., 1992; Felten et al., 1985), the extent and type of sensory innervation of LNs has been difficult to establish because the neuropeptides and ion channels that are traditionally used to identify and manipulate sensory neurons (e.g. CGRP, substance P or the capsaicin receptor, TRPV1 (transient receptor potential channel-vanilloid subfamily member 1) are also expressed on other cell types (Caterina, 2003; Malin et al., 2011; Shepherd et al., 2005b). Moreover, the lack of definitive markers for nonpeptidergic sensory neurons, a subset that may also target LNs, prevents direct assessment of their presence. These caveats notwithstanding, in animal models of arthritis and contact hypersensitivity, local application of capsaicin, a neurotoxin that targets TRPV1, attenuates inflammatory responses, suggesting a pro-inflammatory role for capsaicin-sensitive sensory innervation of LNs (Felten et al., 1992; Lorton et al., 2000; Shepherd et al., 2005a). More recently, a *diphtheria toxin* fragment A (DTA)-based genetic mouse model in which animals are globally deficient in nociceptors revealed a role for sensory neurons in regulating antigen retention and flow through peripheral LNs in immunized mice (Hanes et al., 2016). This observation, together with studies in sheep that showed a stimulatory effect of substance P on lymph flow and lymphocyte recirculation through peripheral LNs (Moore et al., 1989) suggests that modulation of lymphatic trafficking may be one of the mechanisms by which putative sensory innervation could locally regulate immune responses in LNs.

Nonetheless, sensory neuron-immune interactions in LNs have yet to be systematically studied. In particular, we have little information on whether and how the sensory nervous system might regulate specific stroma and/or immune cell subsets within LNs. Here, to directly address this gap in our understanding, we have systemically mapped sensory neurons, immune cells, and stromal cells within skin-draining LNs by integrating state-of-the-art imaging, retrograde labeling, scRNA-seq technologies and optogenetics. We found that sensory neurons innervate LNs with a preferential distribution toward the LN periphery, the region most prone to undergo rapid inflammation-induced mechanical, chemical and cellular changes. Using scRNA-Seq of retrogradely labeled DRG neurons, we identified four molecularly-distinct LN-innervating sensory neuronal subtypes with a strong enrichment for peptidergic nociceptors. To nominate putative cellular communication partners of these sensory neurons, we implemented Seq-Well-based scRNA-Seq to generate a draft single-cell “atlas” of murine steady-state LNs (Aicher et al., 2019; Gierahn et al., 2017). By *in silico* matching known ligands and receptors on both LN-innervating sensory neurons and each subset of LN resident stromal and immune cells, we determined that stromal cells exhibit the highest potential for interaction with local sensory fibers. Finally, using optogenetic triggering, we stimulated LN-innervating sensory neurons and then profiled LN-resident cell types using single-cell transcriptomics to determine which LN resident cells are most responsive to sensory input, validating our predictions. This strategy allowed us to identify the LN-resident target cell types that were immediately impacted by neuronal activity (i.e. LN cell types exhibiting the largest transcriptional changes after optogenetic neuronal stimulation), which included lymphatic endothelial cells, non-HEV blood endothelial cells, non-endothelial stroma, and certain populations of innate leukocytes. Together, our results define the anatomic and molecular identity of a previously enigmatic population of sensory neurons that innervate LNs, and uncover a sensory neuron-stroma axis within steady-state LNs. The experimental and computational frameworks, and foundational datasets, established within this study should be broadly applicable to future analysis of neural circuits in a wide variety of tissues.

## RESULTS

### Lymph nodes are innervated by both sensory and sympathetic neurons

To establish whether, where, and to what extent LNs are innervated by sensory fibers, we pursued several complementary strategies. First, we took a genetic approach to label peripheral neurons of sensory lineage, including nociceptors, with tdTomato using ‘knock-in’ mice that expressed Cre recombinase under control of the *Nav1.8* locus, which encodes a nociceptor-enriched voltage gated sodium channel (Nassar et al., 2004). We then processed popliteal LNs (popLNs) from these animals to visualize the complete three-dimensional morphology of nerve fibers within the intact organs using a protocol adapted from a previously described tissue imaging method, immunolabeling-enabled three-dimensional imaging of solvent-cleared organs (iDISCO; see **Methods**) (Renier et al., 2014). By co-staining for tdTomato and the pan-neuronal marker, β3-tubulin, tdTomato^+^ nerve fibers, presumably originating from primary sensory neurons, were readily detectable as a major component of the neuronal architecture within and around popLNs (Figure 1A, **Movie 1**). Consistent with previous studies that had described sympathetic innervation of LNs (Bellinger et al., 1992; Felten et al., 1985), popLNs also contained a sizeable population of tdTomato^−^ neuronal fibers that expressed tyrosine hydroxylase (TH), a prototypical marker for sympathetic neurons (Figure 1B, **Movie 2**). In comparison, cholinergic parasympathetic fibers genetically labeled with GFP in *ChAT^BAC^-eGFP* animals, where eGFP expression was driven by endogenous choline acetyltransferase (ChAT) transcriptional regulatory elements within a transgene, were never observed in LNs, even though scattered GFP^+^ cells were found in the same LNs, likely representing previously described ChAT^+^ immune cells (Reardon et al., 2013; Rosas-Ballina et al., 2011; Tallini et al., 2006) (**Figure S1A**). As previously noted (Bellinger et al., 1992), the primary path of nerve entry into LNs closely tracked the major blood vessels that entered the LN in the hilus region (Figure 1B, **Movie 2**). Incoming nerve fibers preferentially traveled along a subset of blood vessels that were identified as small arteries and arterioles based on selective genetic labeling in *Bmx-CreER^T2^* x *Rosa26^eYFP/+^*mice in which arterial endothelial cells (ECs) express YFP (Ehling et al., 2013) (**Figure S1B**). Within the LN, the arborization pattern of tdTomato^−^ TH^+^ (putative sympathetic) neurons was notably distinct from that of tdTomato^+^ TH^−^ (putative sensory) neurons. While TH^+^ neurons primarily innervated the vasculature and often wrapped tightly around blood vessels, tdTomato^+^ fibers displayed a distinct terminal morphology and not only aligned with vasculature, but also branched extensively into the interstitial space.

**Figure 1:**
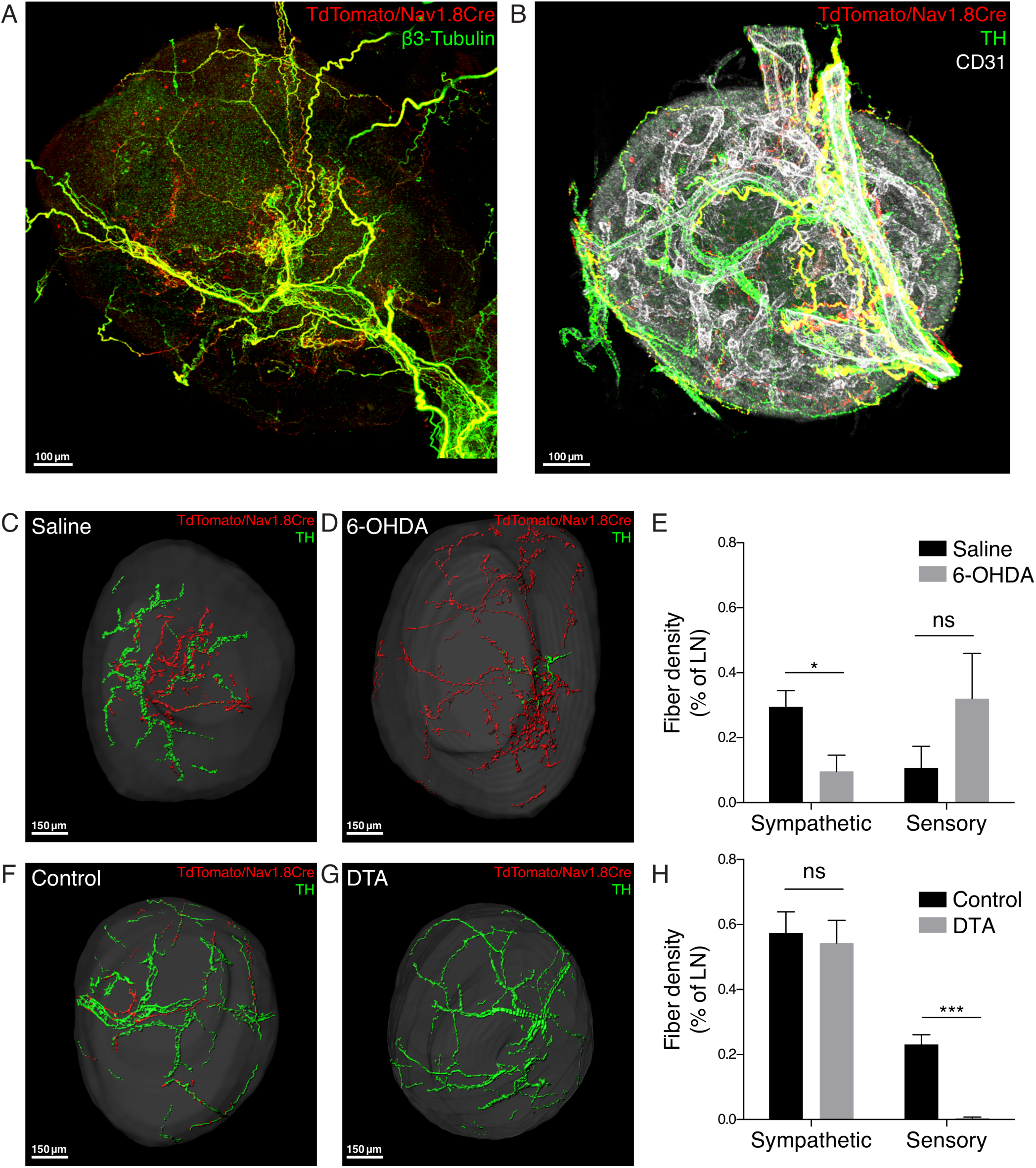
Dual innervation of peripheral lymph nodes (LNs) by sensory and sympathetic neurons. **A, B.** Representative 3D reconstructions of murine LNs stained for sensory neurons (red) and (**A**) the pan-neuronal marker β3-tubulin (green) or (**B**) tyrosine hydroxylase (TH; green) and CD31 (white) to mark sympathetic fibers and vasculature, respectively. Whole-mount preparations of popliteal LNs from a *Nav1.8^Cre/+^ x Rosa26^LSL-tdTomato/+^* mouse were stained for tdTomato (red) to identify sensory neurons and with MAbs against other neuronal markers and imaged by confocal microscopy. **C, D.** Representative rendered surfaces for tdTomato^+^ sensory fibers (red) and TH^+^ sympathetic fibers (green) within popliteal LNs (grey) of saline-treated (**C**) and 6-OHDA-treated (**D**) *Nav1.8^Cre/+^ x Rosa26^LSL-tdTomato/+^* mice. **E.** Quantification of the effect of 6-OHDA treatment on sensory and sympathetic fiber density. n = 5 LNs per group of 3 control and 3 OHDA-treated mice from 3 litters. For sympathetic fibers, p=0.0232 (*); for sensory fibers, p=0.2050 (ns) by unpaired t-test. **F, G.** Representative rendered surfaces for tdTomato^+^ sensory fibers (red) and TH^+^ sympathetic fibers (green) within rendered popliteal LNs (grey) of age matched *Nav1.8^Cre/+^ x Rosa26^LSL-tdTomato/+^* (**F**) and Nav1.8-DTA (**G**) mice. **H**. Quantification of the effect of DTA-induced developmental ablation of Nav1.8^+^ neurons on sensory and sympathetic fiber density. n = 6 LNs per group of 3 mutant and 3 littermate control mice from 3 litters. For sympathetic

Sensory and sympathetic neurons are traditionally defined based on the location of their cell bodies, with the former residing in vagal ganglia or DRGs, and the latter in sympathetic ganglia, which each represent anatomically segregated structures that can be reliably identified. Even though the above experiments clearly established that LNs are innervated by both TH^+^ and Nav1.8 lineage neurons, it should be cautioned that some Nav1.8 lineage neurons are also found within sympathetic ganglia (Gautron et al., 2011), and TH^+^ low threshold mechanoreceptors are present in DRGs (Li et al., 2011). Thus, although TH and Nav1.8 are useful to detect the bulk of sympathetic and sensory fibers, respectively, neither marker is truly specific. Therefore, to rigorously establish the anatomic origin of intranodal fibers, we retrogradely labeled the cell bodies of LN-innervating neurons in DRGs and sympathetic ganglia by injecting a fluorescent neuronal tracer, WGA-AF488, into inguinal LNs (iLNs) of *Nav1.8^Cre/+^* x *Rosa26^tdTomato/+^* mice (Robertson, 1990) (**Figure S1C,** see **Methods**). Expression of tdTomato and TH within WGA^+^ populations revealed that >90% of WGA^+^ neurons in DRGs and sympathetic ganglia were tdTomato^+^TH^-^ and tdTomato^-^TH^+^, respectively, confirming that Nav1.8 and TH adequately and specifically identify sensory and sympathetic innervation of LNs (**Figures S1D-S1H**). No WGA^+^ neurons were detected in vagal ganglia, indicating that iLNs receive sensory innervation exclusively from DRGs (data not shown).

Having confirmed that LNs are indeed innervated by both sensory and sympathetic fibers, we next asked whether either type of innervation was dependent on the presence of the other. Thus, we assessed the differential sensitivity of LN innervating neurons to either 6-hydroxydopamine (6-OHDA)-mediated chemical sympathectomy or diphtheria toxin A (DTA)-mediated genetic ablation of Nav1.8 lineage neurons. 6-OHDA treatment efficiently depleted LNs of sympathetic fibers, but had no significant effect on Nav1.8^+^ sensory fibers (Figures 1C-1E). Conversely, ablation of Nav1.8 lineage neurons in *Nav1.8^Cre/+^ x Rosa26^DTA/tdTomato^* (Nav1.8-DTA) mice (Abrahamsen et al., 2008) resulted in a profound loss of sensory fibers without affecting TH^+^ fibers (Figures 1F-1H). Thus, LNs are innervated by both sympathetic and sensory fibers, two categorically distinct types of innervation that are mutually independent.

### Sensory neurons preferentially innervate the periphery of LNs

Next, we set out to map the spatial distribution of sensory fibers within LNs. LNs are anatomically divided into the lymphocyte-rich cortex, consisting of superficial B follicles and the deep T cell area, and the medulla, which connects to the hilus and contains abundant macrophages and lymphatics. The entire organ is covered by a collagen-rich capsule, which is penetrated by afferent lymph vessels that discharge lymph into the subcapsular sinus (SCS), a fluid filled space that is densely lined by specialized macrophages and lymphatic endothelial cells (LECs). All LECs in LNs, including the outermost layer in the SCS, are readily detectable in *Prox1-EGFP* mice in which GFP is selectively expressed in LECs (Choi et al., 2011). Thus, in *Nav1.8^Cre/+^* x *Rosa26^tdTomato/+^*; *Prox1-EGFP* animals, we were able to identify intranodal sensory (i.e. tdTomato^+^) fibers, as well as determine their spatial relationship to the GFP^+^ lymphatic network and their distance to the outermost boundaries of the LN (**Figure S2A,** see **Methods**). Any tdTomato^+^ sensory fiber within or below the GFP^+^ outer LEC lining was considered intranodal. Notably, the intranodal sensory fibers penetrated into the LN parenchyma to a relatively shallow depth with an average maximum penetration distance of 112 µm ± 29 µm (mean ± SEM) below the capsule (Figure 2A, **Movie 3**). In fact, the majority (∼60%) of intranodal sensory fibers were located <10 μm below the surface of popLNs (Figure 2B). Most of these penetrating tdTomato^+^ fibers were located in the medulla, marked by the LEC restricted marker, LYVE-1. By contrast, tdTomato^+^ fibers were rarely seen in the deep LN cortex (Figure 2C, **Movie 4**).

**Figure 2:**
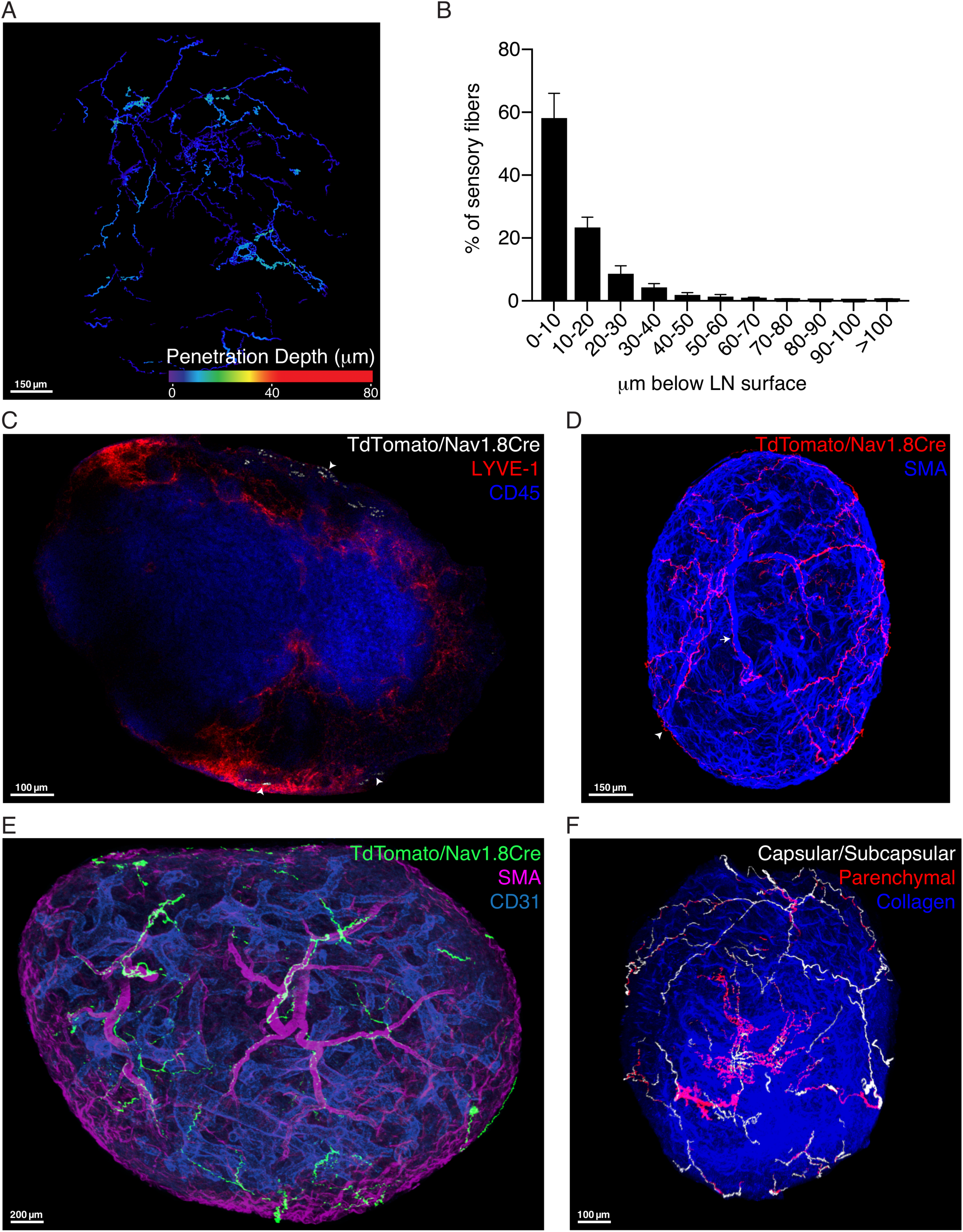
Spatial distribution of sensory innervation of peripheral LNs. **A**. 3D reconstruction of a representative confocal image of tdTomato^+^ sensory fibers within popliteal LNs of *Nav1.8^Cre/+^; Rosa26^LSL-tdTomato/+^; Prox1-EGFP* animals color-coded by penetration depth, i.e., the shortest distance of a point within sensory fibers to the LN surface outlined based on the outermost layer of GFP^+^ LECs. **B.** Quantification of the penetration depth of tdTomato^+^ sensory fibers in popliteal LNs of *Nav1.8^Cre/+^; Rosa26^LSL-tdTomato/+^; Prox1-EGFP* animals as percentage of total intranodal sensory fibers found in 10 μm bins from 0 to 100 μm away from the LN surface, (a total of 5 LNs from 3 mice). **C.** A representative confocal section of whole-mount popliteal LNs from *Nav1.8^Cre/+^; Rosa26^LSL-tdTomato/+^; Prox1-EGFP* animals, stained for tdTomato (white), LYVE-1 (red) and CD45 (blue) illustrating the spatial relationship between sensory fibers (arrowheads) and the cortex and the medulla of LNs **D.** 3D reconstruction of a representative confocal image of whole-mount popliteal LNs from *Nav1.8^Cre/+^; Rosa26^LSL-tdTomato/LSL-tdTomato^* animals, stained for tdTomato (red) and smooth muscle cell actin (SMA) (blue) highlighting the two main plexuses of sensory nerves within LNs, i.e., perivascular (arrow) and capsular/subcapsular (arrowhead) plexuses. **E.** 3D reconstruction of a representative confocal image of whole-mount popliteal LNs from *Nav1.8^Cre^; Rosa26^LSL-tdTomato/LSL-tdTomato^* animals, stained for tdTomato (green), SMA (magenta), CD31 (cyan) demonstrating preferential association between arterioles and sensory fibers inside LNs. **F.** 3D reconstruction of a representative confocal image of whole-mount popliteal LNs from *Nav1.8^Cre/+^; Rosa26^LSL-tdTomato/+^; Prox1-EGFP* animals, stained for tdTomato, GFP, and collagen type 1 (blue) showing the capsular/subcapsular plexus of sensory nerves (white) in relation to the parenchymal sensory fibers

Within the more densely-innervated outer cortical region of LNs, two plexuses of sensory fibers were apparent, one perivascular and the other capsular/subcapsular (Figures 2D). The perivascular fibers coursed through the medulla in tight association with arterioles and mostly terminated before these vessels gave rise to the capillary network. Sensory fibers were generally absent from postcapillary high endothelial venules (HEVs), the paracortical microvascular segments that express peripheral node addressin (PNAd) where blood-borne lymphocytes adhere and emigrate into LNs (von Andrian and Mempel, 2003) (Figure 2E, **S2B, Movies 5, 6**). Occasionally, individual axons were seen to branch away from the vasculature and meander in the interstitial space (Figure 2E, **Movie 5**). Fibers that formed the capsular/subcapsular network typically branched from larger perivascular axon bundles and ramified extensively within the collagen-rich capsule and, in some cases, extended into and below the subcapsular space making contact with CD169^+^ SCS macrophages (Figures 2F, **S2B-S2D, Movie 7**).

Together, these imaging studies indicate that the deep LN cortex where the majority of naive lymphocytes reside is essentially devoid of sensory innervation, whereas cells in the LN periphery, particularly those within the perivascular and subcapsular space, are in close proximity to sensory fibers, suggesting potential regionally confined functional interactions. It is tempting to speculate that the conspicuous concentration of sensory fibers in the outermost LN cortex may allow for rapid and sensitive monitoring of the organ’s physiologic state because the characteristic rapid swelling of reactive LNs would arguably result in mechanical stretching of superficially located fibers. Moreover, inflammatory processes in peripheral tissues may alter the composition of afferent lymph that could provide biochemical cues detectable by sensory fibers within the lymph conduits in the SCS and medulla of draining LNs.

### LN-innervating sensory neurons are heterogeneous and dominated by peptidergic nociceptors

Recent work has shown that DRGs harbor at least 11 subsets of molecularly distinct sensory neurons (Usoskin et al., 2015). In light of this heterogeneity, we set out to define the composition of LN-innervating sensory neurons using an unbiased scRNA-seq-based approach. To this end, we implemented a Cre-lox based viral labeling strategy that enables reliable identification and isolation of the cell bodies of LN-innervating neurons in DRGs (Figure 3A, see **Methods**). Briefly, we injected a Cre-expressing recombinant adeno-associated virus, AAV2/1-Cre (a serotype with broad tropism towards DRG neurons (Kuehn et al., 2019; Mason et al., 2010)) into the right iLN of *Rosa26^LSL-tdTomato/LSL-tdTomato^* mice carrying a Cre-dependent tdTomato reporter. Upon entry into sensory fibers, this non-replicating virus travels retrogradely to the cell body in DRGs where virally encoded Cre recombinase deletes a genomic floxed ‘stop’ sequence resulting in expression of tdTomato. Since the relative positions of DRGs along the anteroposterior axis largely match those of their respective target fields, a specific injection should lead to labeling of DRG neurons at the axial level consistent with that of the targeted LN. Indeed, following unilateral iLN injection, robust tdTomato expression was consistently observed in a handful of neurons in the ipsilateral last thoracic (T13) and first lumbar (L1) DRGs, which supply the inguinal region (Takahashi and Nakajima, 1996) (Figures 3B-3D). TdTomato labeling at the site of injection was primarily concentrated within the injected iLN, indicating spatial confinement of the injected material (**Figure S3A**). However, some tdTomato^+^ stromal cells were also detectable in the immediate vicinity of injected LNs, which raised the possibility that some sensory fibers outside the LN were inadvertently labeled. To control for this possibility, we assessed the extent of retrograde labeling of DRG neurons following deliberate perinodal injection of the same amount of virus. In comparison with intranodal injections, which consistently resulted in robust labeling of DRG neurons (16.2 ± 2.4 cells per mouse (mean ± SEM), n = 4), very few (1.5 ± 0.6, n = 4) DRG neurons were labeled after deliberate perinodal injection of AAV2/1-Cre (**Figures S3B, S3C**).

**Figure 3:**
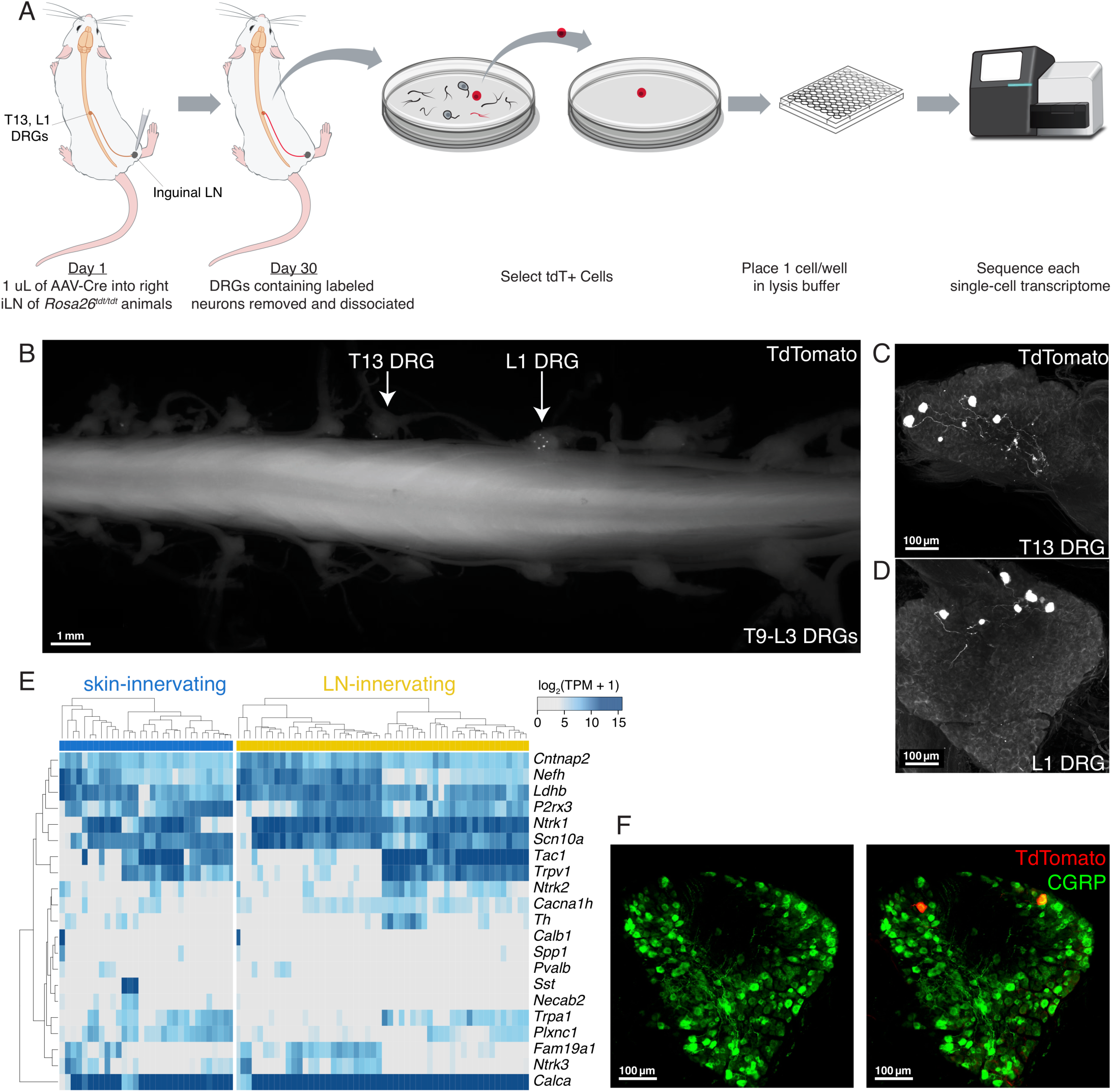
Retrograde labeling of LN-innervating sensory neurons for single-cell RNA-seq (scRNA-seq) **A.** Schematic of viral-based, long-term retrograde labeling from the LN, manual single-cell sorting and scRNA-seq pipeline. **B.** A representative epifluorescence image of tdTomato^+^ retrogradely-labeled iLN-innervating DRG neurons in a whole-mount spinal cord-DRG preparation without antibody amplification. **C** and **D.** Maximum projection view of confocal images of whole-mount ipsilateral T13 (**C**) and L1 (**D**) DRGs from **B** stained for tdTomato. **E**. Single-cell gene expression of neuronal subtype-specific markers in LN-innervating and skin-innervating sensory neurons. **F.** Representative confocal sections of whole-mount DRGs containing tdTomato^+^ retrogradely-labeled iLN-innervating neurons from *Rosa26^LSL-tdTomato/LSL-tdTomato^* animals following intranodal injection of AAV-Cre, stained for tdTomato (red) and CGRP (green). Percentage of tdTomato^+^ sensory neurons that express CGRP: 88.39% (mean) ± 8.672% (SEM) based on a total of 44 tdTomato^+^ neurons from 3 mice.

To independently evaluate the specificity of our retrograde viral labeling strategy, we also performed intranodal injections of another AAV2/1 vector carrying a Cre-dependent tdTomato cassette (AAV-Flex-tdTomato) in *Nav1.8^Cre^* animals, in which sensory neurons express Cre recombinase. The innervation pattern of retrogradely-labeled tdTomato^+^ neurons was strikingly similar to that of tdTomato^+^ LN-innervating sensory neurons in *Nav1.8^Cre/+^* x *Rosa26^tdTomato/+^* animals, thus confirming their identity (Figures 2D, **S3D** and **Movie 8**).

Having established that intranodal retrograde labeling with AAV2/1-Cre vector allows for selective fluorescent tagging of the cell bodies of true LN-innervating sensory neurons within DRGs, we harvested ipsilateral T13 and L1 DRGs from injected animals and manually isolated individual tdTomato^+^ neurons as previously described (Hempel et al., 2007). Each isolated cell was then subjected to scRNA-Seq using the SMART-seq2 protocol to yield a final dataset of 52 LN-innervating sensory neurons from 8 mice (average +/- SEM genes/cell: 9,843 +/- 229) (Picelli et al., 2014; Trombetta et al., 2014) (**Table S1**, see **Methods)**. As a control and reference population, we also generated scRNA-Seq libraries from 31 skin-innervating neurons from 4 *Rosa26^LSL-tdTomato/LSL-tdTomato^* mice that were retrogradely labeled by intradermal injection of AAV2/1-Cre (average +/- SEM genes/cell: 9,653 +/- 302) (**Table S1**, **Figure S3E)**. The specificity of retrograde labeling from the skin was demonstrated by revealing the axonal terminals of retrogradely-labeled Nav1.8^+^ neurons following intradermal injection of AAV2/1-Flex-eGFP into *Nav1.8^Cre^* animals, as well as lack of tdTomato labeling in draining LNs upon intradermal injection of AAV2/1-Cre into *Rosa26^LSL-tdTomato/LSL-tdTomato^* animals (data not shown) (**Figure S3F**).

To define the molecular identity of these LN-innervating sensory neurons, we examined each for expression of canonical markers of known sensory neuron molecular subtypes (Usoskin et al., 2015) (Figure 3E). As expected, the majority (96% with log_2_(1 + TPM) > 1) of LN-innervating sensory neurons expressed Nav1.8 (*Scn10a*), whereas few (23% with log_2_(1 + TPM) > 1) co-expressed TH (*Th*). Almost all LN neurons in which *Nav1.8* was detectable also expressed the high affinity receptor for nerve growth factor (NGF), TrkA (*Ntrk1)*, and the calcitonin gene-related peptide, CGRP (*Calca)*. By contrast, there was little expression of canonical markers for low-threshold mechanoreceptors, proprioceptors, and nonpeptidergic nociceptors, suggesting that the majority of LN-innervating sensory neurons are peptidergic nociceptors. Indeed, 88.4% ± 8.7% (mean ± SEM) of retrogradely-labeled LN-innervating DRG neurons expressed the CGRP neuropeptide by immunohistochemistry (Figure 3F). Notably, mutually exclusive expression of substance P (*Tac1*) and neurofilament heavy chain (NFH; gene name *Nefh*) within *Calca*^+^ LN-innervating sensory neurons allowed the identification of two LN-innervating peptidergic nociceptor subclasses akin to the previously defined PEP1 and PEP2 clusters which correspond to thermosensitive unmyelinated nociceptors and lightly myelinated Aδ nociceptors, respectively (Usoskin et al., 2015) (Figure 3E). Consistent with the heterogeneous expression of *Nefh*, a marker for medium-to-large diameter sensory neurons with myelinated axons (Rice and Albrecht, 2008), whole-mount DRG staining revealed that retrogradely-labeled LN-innervating sensory neurons were heterogeneous with respect to soma size. This diversity of cell dimensions matched the range of diameters observed in CGRP^+^ neurons, which are known to include neurons of different sizes (Lawson et al., 2002) (**Figure S3G**). Furthermore, NFH^+^ myelinated and NFH^-^ unmyelinated sensory fibers were both abundant in the perivascular and capsular/subcapsular space of LNs (**Figure S3H**).

To look beyond the expression of canonical DRG neuronal subtype markers alone, we next sought to contextualize our scRNA-Seq profiles of LN-innervating sensory neurons against a published scRNA-Seq Sensory Neuron Atlas (Usoskin et al., 2015). Using the published single-cell transcriptomic profiles of 622 DRG neurons, we calculated principal components (PC) over all neuronal cells and projected our LN- and skin-innervating sensory neurons into this PC space (Figure 4A, see **Methods**). Consistent with the strong peptidergic features (i.e., *Calca, Ntrk1* expression) described above, LN-innervating sensory neurons were distributed over an area in PC space in closest proximity to the peptidergic neurons (PEP) defined by Usoskin et al. (Figure 4B). To directly classify LN-innervating or skin-innervating sensory neurons relative to the 11 published DRG subtypes, we created pseudopopulation averages from single-cell transcriptomes of each subtype, and calculated the Spearman correlation between single LN-innervating or skin-innervating sensory neurons and the neuronal subtype pseudopopulations (Figure 4C). Using hierarchical clustering based upon the similarity of our single neurons to the neuronal subtypes defined by Usoskin et al., we recovered 4 major transcriptionally distinct neuronal classes within our dataset, termed Neuron Types 1 to 4. Each Neuron Type was represented, albeit in very different proportions, in both LN-innervating and skin-innervating sensory neurons, demonstrating both intrinsic heterogeneity within sensory neurons innervating the same target, as well as innervation target-dependent differences in subtype composition (p < 0.001 by Pearson’s Chi-square test, Figure 4D). Neuron Types 1 and 3, which share the strongest similarity with the PEP1 and PEP2 subtypes defined by Usoskin et al., were enriched in the LN-innervating population relative to the skin-innervating population (LN-innervating: 48% Neuron Type 1, 44% Neuron Type 3; skin-innervating: 29% Neuron Type 1, 16% Neuron Type 3). Conversely, Neuron Types 2 and 4, which correspond to nonpeptidergic nociceptors and myelinated non-nociceptors, respectively, were underrepresented in the LN-innervating population compared to the skin-innervating population (LN-innervating: 2% Neuron Type 2, 6% Neuron Type 4; skin-innervating: 45% Neuron Type 2, 10% Neuron Type 4).

**Figure 4:**
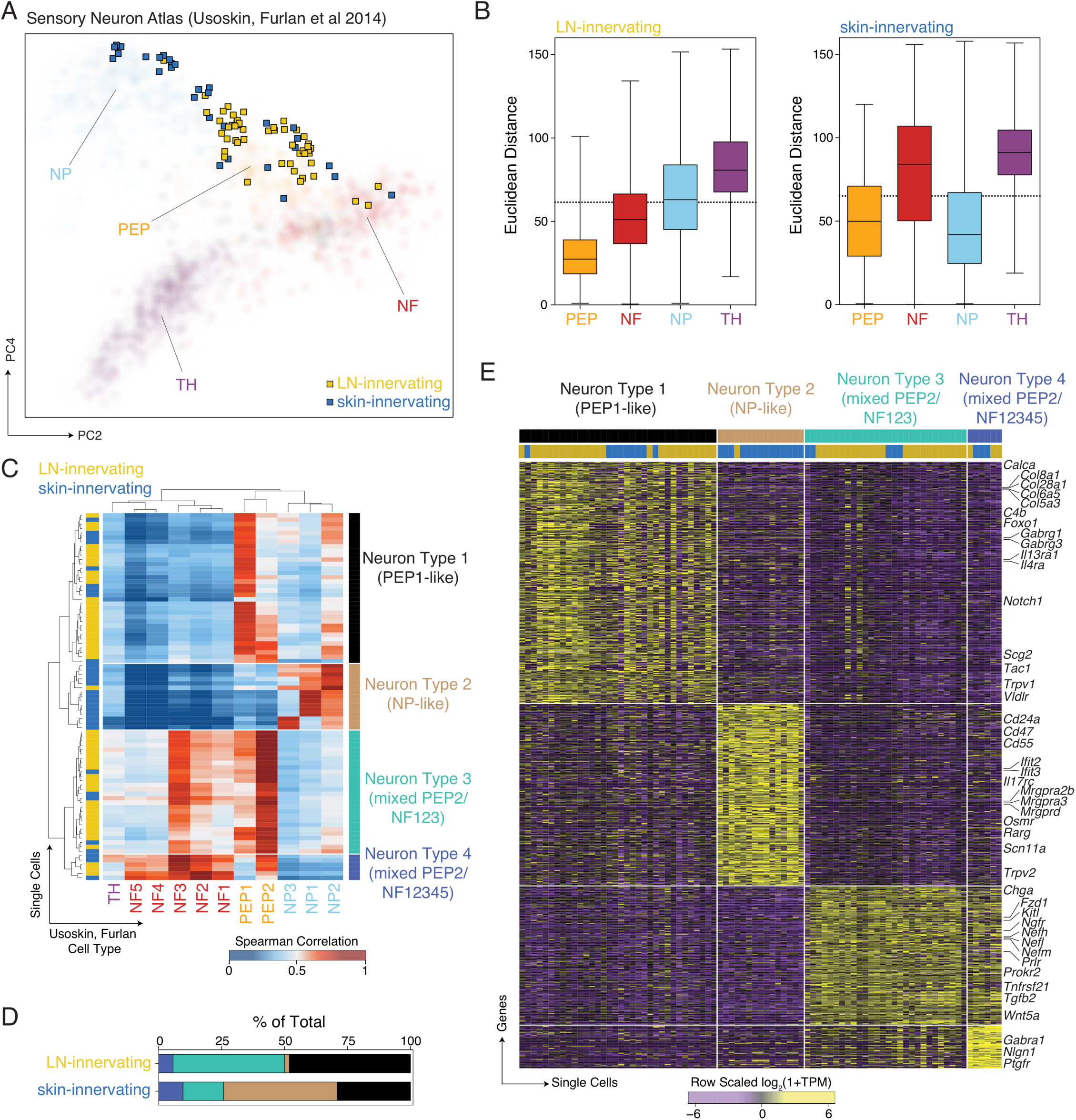
LN-innervating sensory neurons are primarily peptidergic nociceptors. **A.** Principal Components 2 vs. 4 of Usoskin, Furlan et al. Sensory Neuron Atlas (Usoskin et al., 2015), represented by transparent circles, colored by previously-defined cell types: non-peptidergic nociceptors (NP, light blue); peptidergic nociceptors (PEP, orange); neurofilament containing (NF, red); and tyrosine hydroxylase containing (TH, purple). LN-innervating (yellow squares) and skin-innervating (blue squares) neurons are projected onto the PC space. **B**. Euclidean distance between each LN-innervating neuron (left) or skin-innervating neuron (right) and neurons in the Usoskin, Furlan et al. Sensory Neuron Atlas (Usoskin et al., 2015), separated by cell type. Dashed lines represent the 99% confidence interval for distance between single cells categorized as the same cell type within the Sensory Neuron Atlas. Box represents 25-75 quantiles, error bars span min-max range. **C.** Spearman correlation between the scRNA-seq profiles of LN- or skin-innervating neurons and neuronal subsets from the Usoskin et al. Sensory Neuron Atlas. Hierarchical clustering divides LN- and skin-innervating neurons into 4 major subtypes: Neuron Type 1 (PEP1-like, black); Neuron Type 2 (NP-like, tan); Neuron Type 3 (mixed PEP2/NF123, turquoise); and Neuron Type 4 (mixed PEP2/NF12345, dark blue). **D.** Distribution of Neuron Types 1-4 by innervation target (LN-innervating, n=52; skin-innervating, n=31). **E.** Differentially expressed genes (SCDE, Holm adjusted p-value < 0.01) between each Neuron Type vs. all other Neuron Types. Top color bars

To further understand the distinct molecular phenotypes represented by Neuron Types, we performed differential expression analysis and discovered unique gene modules that cleanly define each Neuron Type (Figure 4E, **Table S2**). Importantly, these gene modules were consistent with the general phenotypic descriptions of the corresponding subtypes in Usoskin et al. (e.g., *Tac1^+^* and *Nefh^+^* for Neuron Types 1 and 3, respectively). Together, these data suggest that LN-innervating sensory neurons are heterogeneous at the transcriptomic level, yet are strongly biased toward peptidergic phenotypes.

### LN-innervating sensory neurons are molecularly distinct from their skin-innervating counterparts

In view of the observed innervation target-dependent differences in the representation of sensory neuron subtypes, we next assessed differences in gene expression between LN-innervating and skin-innervating sensory neurons to define gene programs that may support target tissue-specific development and function. A direct comparison of LN-innervating and skin-innervating sensory neurons identified 101 and 156 genes that were significantly upregulated in LN-innervating and skin-innervating neurons, respectively (Holm adjusted p-value < 0.05; Figures 5A, 5B, **Table S2**). While some of these differentially expressed (DE) genes could reflect differential subtype composition, robust gene expression differences between LN- and skin-innervating neurons were observed even when the two main neuron types innervating LNs (Neuron Types 1 and 3) were analyzed separately, indicating innervation target-dependent molecular distinction between otherwise highly similar neurons (**Table S2**, **Figures S4A, S4B**). Indeed, when DE genes were analyzed for enriched gene ontologies, LN- and skin-innervating sensory neurons differed with respect to many surface ion channels and synaptic proteins (LN-specific: *Trpc4, Trpm8, Kcnh5, Ache*; skin-specific: *Trpc3, Trpc6, Kctd16, Synpr, Gabra1, Kcnk12*), as well as secreted and cell surface molecules, which may reflect target-specific modes of communication between sensory neurons and their microenvironment (Figures 5C, 5D, **S4C-S4E**). The upregulation of *Ache* in LN neurons relative to skin neurons is particularly intriguing, considering that the protein product, acetylcholinesterase, is the main enzyme that mediates breakdown of acetylcholine, a neurotransmitter that transmits anti-inflammatory signals via the efferent arm of the inflammatory reflex (Tracey, 2007). This suggests the possibility that LN-innervating sensory neurons help shape the acetylcholine content in LNs, thereby fine-tuning cholinergic anti-inflammatory pathways. Moreover, LN-innervating sensory neurons overexpressed genes with inflammatory or immune-cell type interacting functions including *Tbxa2r, Il33, Ptgir,* and *Cd1d,* suggesting additional immunological functions (Figures 5C, 5D, **Table S2**). For example, *Tbxa2r* and *Ptgir* encode receptors for bioactive lipids, namely, the prostanoids thromboxane A_2_ and prostaglandin I2, respectively, both of which can either directly activate or sensitize sensory neurons (Andoh et al., 2007; Bley et al., 1998; Wacker et al., 2002). Thus, LN-innervating sensory neurons possess the molecular machinery to monitor the inflammatory state of LNs by sensing inflammation-induced prostanoids that may either reach LNs via the lymph or be produced directly within reactive LNs.

**Figure 5:**
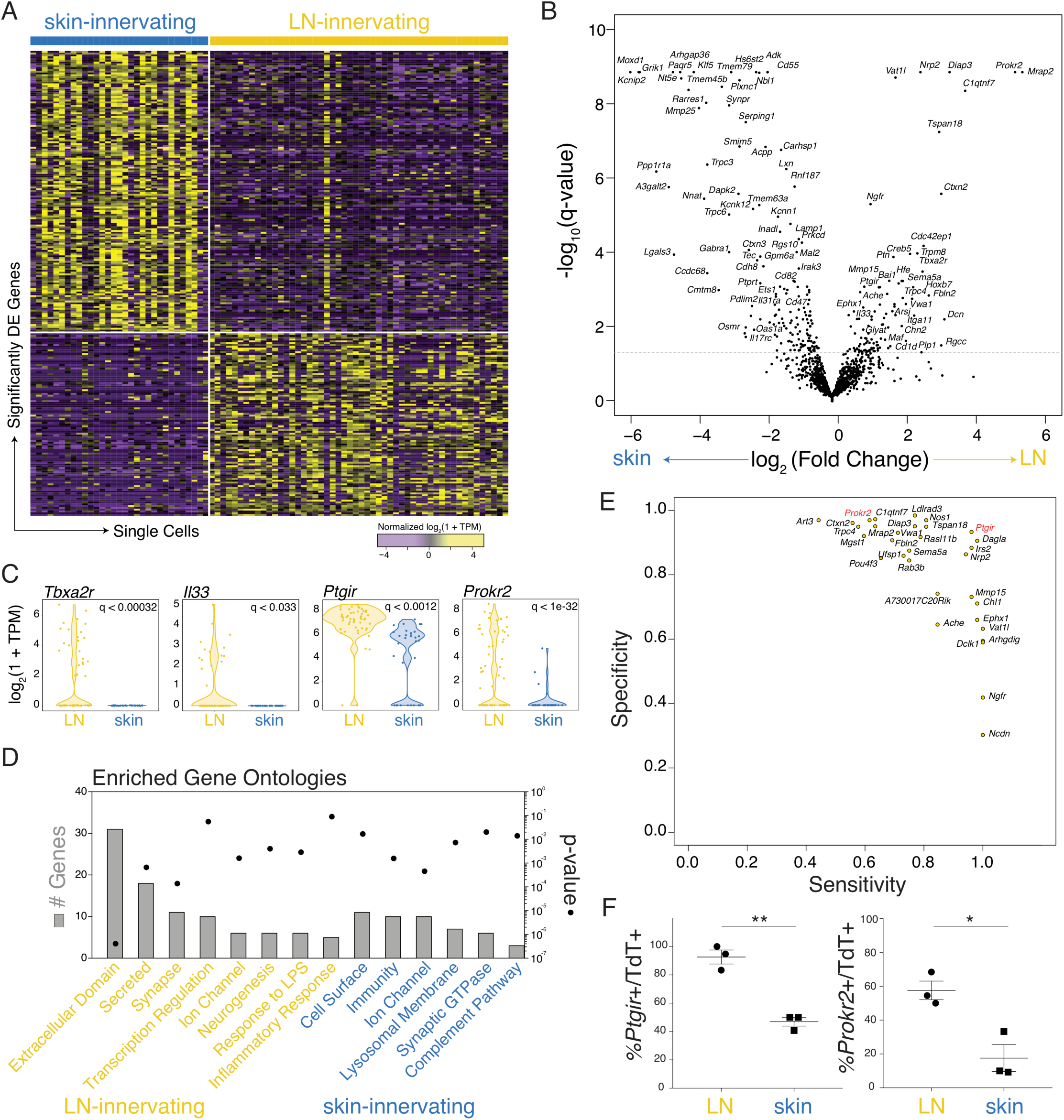
LN-innervating sensory neurons express unique defining markers and functional pathways. **A**. Heatmap of significantly differentially expressed (DE) genes (Holm corrected p-value <0.05, upregulated in LN-innervating: 101 genes; upregulated in skin-innervating: 156 genes). **B.** Volcano plot comparing fold-change differences and -log10(Holm corrected p-values) highlights significantly DE genes (using SCDE). Horizontal dashed line represents significance cutoff of corrected p-value (q-value) < 0.05. **C**. Violin plots for selected genes significantly upregulated in LN-innervating neurons. **D.** Enriched gene ontologies represented by genes upregulated in LN-innervating neurons (yellow) or skin-innervating neurons (blue). Left y axis: number of DE genes represented by gene ontology term; right y axis: p-value (Fisher’s Exact Test) for gene ontology enrichment. **E**. Identification of sensitive (true positive/(true positive + false negative)) and specific (true negative/(true negative + false positive)) markers for LN-innervating neurons compared to skin-innervating neurons and Usoskin, Furlan et al. Sensory Neuron Atlas. **F**. Quantification of *Ptgir* and *Prokr2* expression in *tdTomato*^+^ retrogradely-labeled LN- or skin-innervating neurons (“TdT+”) as percentage of *tdTomato*^+^ neurons that are *Ptgir*^+^ or *Prokr2*^+^ by RNAscope. Significance assessed by unpaired t-test, p= 0.0014 (**), p= 0.0142 (*), based on a total of 3 mice.

To identify candidate markers for LN-innervating neurons, we compared LN-innervating neurons to both skin-innervating neurons and the full diversity of sensory neurons captured in the previously published Sensory Neuron Atlas (Usoskin et al., 2015) (**Table S2**). To this end, we determined the “true positive” rate (sensitivity) and “true negative” rate (specificity) of LN-innervating neuron gene markers by assessing the fraction of LN-innervating vs. control populations expressing a given gene, and prioritized markers that appeared both specific and selective for LN-innervating neurons (Figure 5E). For example, *Ptgir* was identified as a generic marker for LN-innervating sensory neurons with relatively high specificity, while *Prokr2* appeared to be more specifically expressed by LN-innervating neurons, with lower sensitivity and enrichment within Neuron Types 1 and 3 (Figures 4E, **S4A, S4B**). The expression profiles of *Ptgir* and *Prokr2* in LN- and skin-innervating neurons were further validated by RNAscope-based multiplexed fluorescence *in situ* hybridization analysis of *Ptgir*, *Prokr2*, and *tdTomato* in DRGs containing tdTomato^+^ retrogradely labeled LN- or skin-innervating neurons (Figures 5F, **S4F-S4I**). Thus, in addition to subtype composition differences, sensory neurons innervating the LN are marked by a distinct transcriptional profile that includes high expression of *Ptgir* and *Prokr2*.

### scRNA-seq of LN cells nominates interacting partners of LN-innervating sensory neurons

Our molecular characterization of LN-innervating sensory neurons revealed expression of many genes and cellular programs poised to support interactions with other LN-resident cells. Thus, we set out to systematically map cellular interactions between the sensory nervous system and the various cell types that reside in LNs. To complete this effort, we first needed to realize a map of all LN cell subsets at the molecular level. We therefore generated a single-cell transcriptomic atlas of steady-state murine inguinal LNs (n=7) using the Seq-Well platform (Aicher et al., 2019; Gierahn et al., 2017) (**Table S3**, see **Methods**).

To minimize biases introduced during tissue dissociation, a gentle dissociation protocol optimized for reliable isolation of both stromal and hemopoietic LN cells was used to generate a suspension of single cells from both the non-immune and the immune compartments (Fletcher et al., 2011). To increase coverage of the many rare LN cell types – i.e., the non-T, non-B cells – which populate the preferentially-innervated LN periphery (Figure 2C), we profiled paired LN samples from before and after immuno-magnetic depletion of T and B cells (Figure 6A). Following quality filtering and preprocessing, we recovered libraries from 9,622 single cells and 25,929 unique genes (see **Methods**). For unbiased cell type identification, we reduced this high-dimensional data into a lower-dimensional manifold using principal component analysis (PCA) over variable genes, clustered cells using a mutual k nearest-neighbor graph, and visualized these clusters on t-distributed stochastic neighbor embedding (t-SNE) (Butler et al., 2018) (Figures 6B, **S5A-5O**). This approach uncovered 24 distinct cell types representing all major known lymphoid, myeloid and stromal populations in LNs. To name cell clusters, we identified gene signatures that defined each using a likelihood ratio test, and annotated based on well-defined markers of cell identity (Figure 6C, **S5P, Table S4**). By using this approach, most cell type clusters could be readily identified by their expression of canonical markers (e.g. co-expression of *Cd19, Cd22, Cd79a,* and *Cd79b* in B cells).

**Figure 6:**
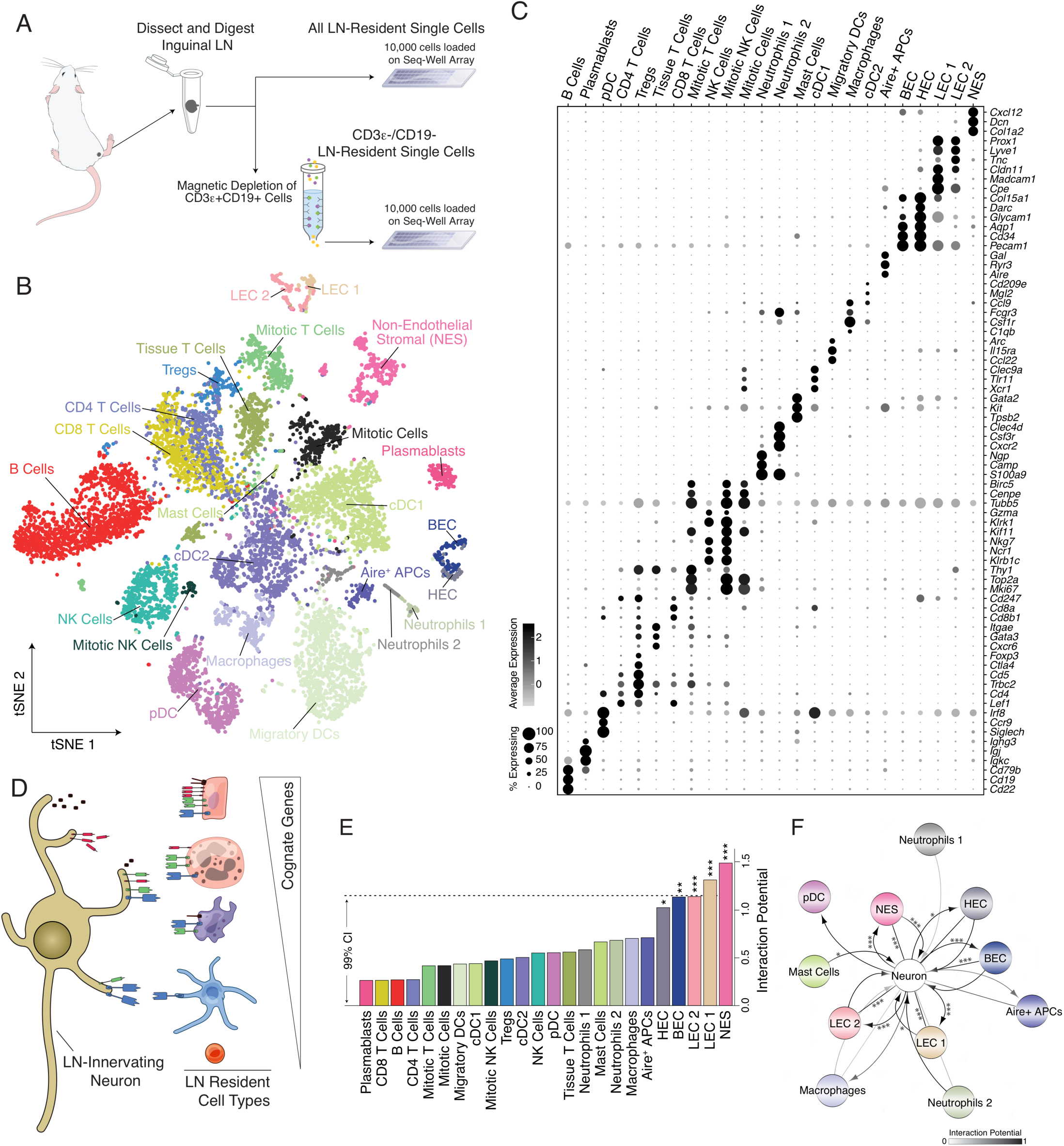
Single-cell transcriptomic profiling of iLN cells nominates likely interacting partners of iLN-innervating sensory neurons. **A.** Schematic for iLN isolation, dissociation, enrichment for rare iLN cell types. **B.** Visualization of cell types recovered by scRNA-seq of 9,662 cells using t-distributed stochastic neighbor embedding (tSNE). **C.** Dot plot representation of genes that distinguish major cell types within LNs (circle diameter reflects the percent of cells expressing a given marker within a cell type, circle color reflects relative expression abundance within a cell type; light grey: low, black: high) All highlighted genes are significantly upregulated in the corresponding cell type, with FDR-corrected p-value < by likelihood ratio test. **D.** Schematic of analysis of the expression of receptor-ligand pairs between LN-innervating neurons and potential interacting LN cell types. **E.** Barplot of Interaction Potential by cell type. Dashed lines represent 99% confidence interval over randomized permuted data. * p<0.05, ** p<0.01, *** p<0.001. **F**. Cell-cell interaction network displaying top interaction edges between LN-innervating neurons and non-neuronal LN cell types. Arrows emanating *from* the “Neuron” node indicate Interaction Potential only considering secreted molecules produced by LN-innervating neurons. Arrows emanating *to* the “Neuron” node indicate Interaction Potential only considering secreted molecules from LN resident cell types with cognate receptors expressed by LN-innervating neurons. Un-directed lines reflect Interaction Potential among ligand-receptor pairs with unknown directionality or bidirectional effects. Darker lines indicate higher Interaction Potential scores. * p<0.05, ** p<0.01, ***

Beyond this initial clustering of single cells, multiple cell clusters could be further subdivided into subclusters (**Figure S5A**). In these instances, we re-analyzed cell clusters using methods for unbiased cell type identification as described above, and partitioned them into appropriate subtypes. For example, we recovered two populations of blood endothelial cells, one of which likely corresponds to HEV endothelial cells (HEC), based on uniform and high expression of well-established HEC markers, e.g., *Ccl21a*, *Fut7*, *Chst4* (**Figures S5H, S5I**) (Homeister et al., 2001; Kawashima et al., 2005; Stein et al., 2000; Uchimura et al., 2005). The second blood endothelial cell cluster, termed BEC, includes a heterogeneous set of non-HEV endothelial cells, including arterial, capillary, and non-HEV venular cells (**Figures S5H, S5I**) (Vanlandewijck et al., 2018). Additionally, we identified two distinct populations of *Prox1*^+^ lymphatic endothelial cells (LECs), with LEC 1 likely representing a mixture of LECs lining the floor of SCS and medullary sinuses based on *Madcam1, Msr1, Bmp2, Vcam1,* and *CD274* (PD-L1) expression profiles (Cohen et al., 2014; Cordeiro et al., 2016; Takeda et al., 2019), and LEC 2 defined by unique expression of multiple extracellular matrix or structural proteins, including *Fbln2, Aqp1, Fbln5, Tnc and Reln* (**Figures S5N, S5O**). Interestingly, cells within the LEC 2 cluster shared molecular signatures with LECs positioned at the ceiling of SCS, within the cortical sinuses, and lining lymphatic vessels, including selective expression of *Emcn*, *Ackr4*, *Ackr2*, *Klf2*, *Fabp4* and *Cav1* (Iftakhar et al., 2016; Takeda et al., 2019; Ulvmar et al., 2014). We also identified a subtype of dendritic-cell-like cells (Aire^+^ APC), which likely represent the Aire-expressing ILC3-like cells that were described recently (Yamano et al., 2019). Similarly, we detected two subtypes of neutrophils, Neutrophils 1 and 2, which potentially reflect distinct maturation statuses similar to what was recently described for bone marrow neutrophils (Evrard et al., 2018) (**Figure S5L and S5M**): Neutrophils 1 expressed components of neutrophil granules and effector molecules at high levels, including *Elane, Prtn3, Ctsg, Ngp, Ltf, Camp,* and *Mpo*, whereas Neutrophils 2 expressed little or no effector molecules, but elevated levels of pro-inflammatory genes, including leukocyte traffic molecules, chemokines, cytokines, and cytokine receptors such as *Sell, Ccl4, Cxcr2, Cxcl2, Ccl6, Il1b,* and *Csf3r*. Other cell types were similarly sub-clustered and are discussed within the computational methods (**Figure S5A-S5O,** see **Methods**).

Next, we sought to determine the relative likelihood of each identified LN cell type to interact with LN-innervating sensory neurons by analyzing the expression of ligand-receptor pairs across our two single-cell datasets (Cohen et al., 2018; Smillie et al., 2019; Vento-Tormo et al., 2018). We reasoned that interacting cells might rely on inter-cellular ligand-receptor pairs for signaling crosstalk and/or physical association, either through interaction of membrane-anchored proteins on both cells or via secreted ligands binding to receptors. Therefore, LN cell types with higher expression of cognate receptors or ligands of neuron-expressed ligands or receptors, respectively, should be poised to interact with local sensory innervation. Utilizing public databases of ligand-receptor pairs involved in cellular interaction(Ramilowski et al., 2015), we filtered first for interaction pairs where at least one member was expressed by LN-innervating sensory neurons. Using the respective cognates of each of these molecules, we queried relative expression among all LN cell types (Figures 6D, **S6A**). In this approach, the number of co-expressed ligand-receptor cognates between a LN-innervating neuron and a LN cell type was used to estimate the “Interaction Potential” for that LN cell type (Figures 6E, **S6A**, see **Methods**).

Using this computational strategy, we determined that the non-immune compartment (Non-Endothelial Stroma (NES), BEC, HEC, LEC 1, LEC 2) exhibited the highest Interaction Potential compared to other LN cell types and randomized data (Figure 6E). This ranking was stable across multiple different calculation methods, ligand-receptor databases, and summary statistics, and was not influenced by technical confounders such as cell quality and cell type population size (**Figure S6B-S6E**). To take the directionality of the cognate pairs into consideration, we partitioned ligand-receptor pairs into three categories: (1) the gene product is secreted by LN-innervating neuron, (2) secreted by LN-resident cell types, or (3) non-directional/unknown directionality, often corresponding to interactions between two membrane-tethered proteins. Considering only molecules secreted by LN-innervating neurons (category 1), LEC 2, LEC 1, BEC, HEC, and NES expressed a significantly elevated abundance of cognate receptors (Figure 6F). Given a strong enrichment for peptidergic signatures among LN-innervating neurons, including high expression of CGRP (*Calca, Calcb*), substance P (*Tac1*), galanin (*Gal*), and pituitary adenylate cyclase-activating polypeptide (PACAP) (*Adcyap1)*, we assessed the expression of the corresponding neuropeptide receptors among LN cell types (**Figure S6F**). *Ramp1*, which together with *Calcrl*, a ubiquitously expressed gene among LN cell types, forms the CGRP receptor (McLatchie et al., 1998), was more highly expressed in innate immune cell types such as mast cells and dendritic cells (DCs), suggesting that LN-innervating sensory neurons may signal to select myeloid cell types via CGRP. The receptors for other neuropeptides, *Tac1, Adcyap1,* and *Gal* (*Tacr1*, *Adcyap1r1*, and *Galr2 & Galr1*, respectively) were uniquely expressed by NES, identifying substance P, PACAP, and galanin as potential signaling mediators between LN-innervating neurons and NES. Notably, the recently identified alternative substance P receptor *Mrgprb2* was almost exclusively expressed at high level in LN mast cells, suggesting a substance P/Mrgprb2 mediated sensory neuron-mast cell connection analogous to what was described in the skin (Green et al., 2019). By contrast, classic neuropeptides were not a primary mode of communication between LN-innervating sensory neurons and LN endothelial cells.

To more specifically explore the nature of these inferred axes of communication, we analyzed the cognate receptors and ligands responsible for high Interaction Potentials among the stromal compartments (**Figures S6G, S6H**). For example, predicted interaction with NES was strongly driven by molecules that can either signal both ways (or to and from neighboring cells), including extracellular matrix components (*Col3a1, Col5a2, Col5a1, Col6a1, Col6a2, Col6a3, Col1a2, Col1a2, Lama2, Thbs2, Fn1*) and growth factors/chemokines with diverse roles in neuronal development and function (*Vegfa, Ptn, Mdk, Cxcl12*) (Gonzalez-Castillo et al., 2014; Mackenzie and Ruhrberg, 2012; Mithal et al., 2012; Winkler and Yao, 2014), as well as receptors for growth factors known to regulate fibroblast proliferation and differentiation (*Pdgfra, Pdgfrb, Ntrk2*) (Andrae et al., 2008; Palazzo et al., 2012). Similarly, the putative interactions between blood endothelial cells (BEC and HEC) and LN-innervating sensory neurons were bidirectional, as evidenced by the expression of a distinct set of extracellular matrix and cell adhesion molecules (*Lama5, Itga5, Hspg2*), receptors of central signaling pathways for vascular development (*Flt1, Notch4, Fzd4*) (Mack and Iruela-Arispe, 2018; Shibuya, 2011; Ye et al., 2010), and classic axon guidance molecules with known roles in leukocyte-endothelial adhesion, angiogenesis, and arterial–venous differentiation (*Sema3f*, *Sema7a, Nrp1, Plxnd1, Efnb1, Epha4*) (Adams and Eichmann, 2010; Larrivee et al., 2009). Thus, our single-cell profiling of murine iLN identified stromal cells as the most likely interacting partners of LN-innervating sensory neurons, and revealed potential communication modalities that mediate cellular interactions.

### Optogenetics-assisted validation of local targets of LN-innervating sensory neurons

To directly test functional interactions between LN-innervating sensory neurons and LN cells, we systematically interrogated the effects of acute activation of LN-innervating sensory neurons on gene expression in all identifiable LN cell types by integrating optogenetic stimulation with Seq-Well scRNA-seq profiling. This enabled us to assess the potential neuron-to-immune signaling axis within LNs without *a priori* knowledge of the responding cells downstream. Optogenetics – the combined use of optics and genetics for temporally and spatially precise control of neuronal activity with light – commonly involves targeted expression of a light gated cation channel, e.g. channelrhodopsin-2 (ChR2), in specific neurons of interest, thereby rendering the targeted neurons activatable by blue light (Deisseroth, 2011). To specifically activate LN-innervating neurons *in vivo*, we used *Nav1.8^Cre/+^* x *Rosa26^ChR2-eYFP/+^* (ChR2+) mice in which Nav1.8 lineage neurons expressed ChR2. The iLN-innervating sensory neurons were selectively activated with blue light (473 nm) directed through an optical fiber (200 µm tip diameter) towards a region of the subiliac artery adjacent to the hilus, the predominant site where sensory nerve bundles enter the iLN (Figures 1A, 1B, 7A **and S7A**).

**Figure 7:**
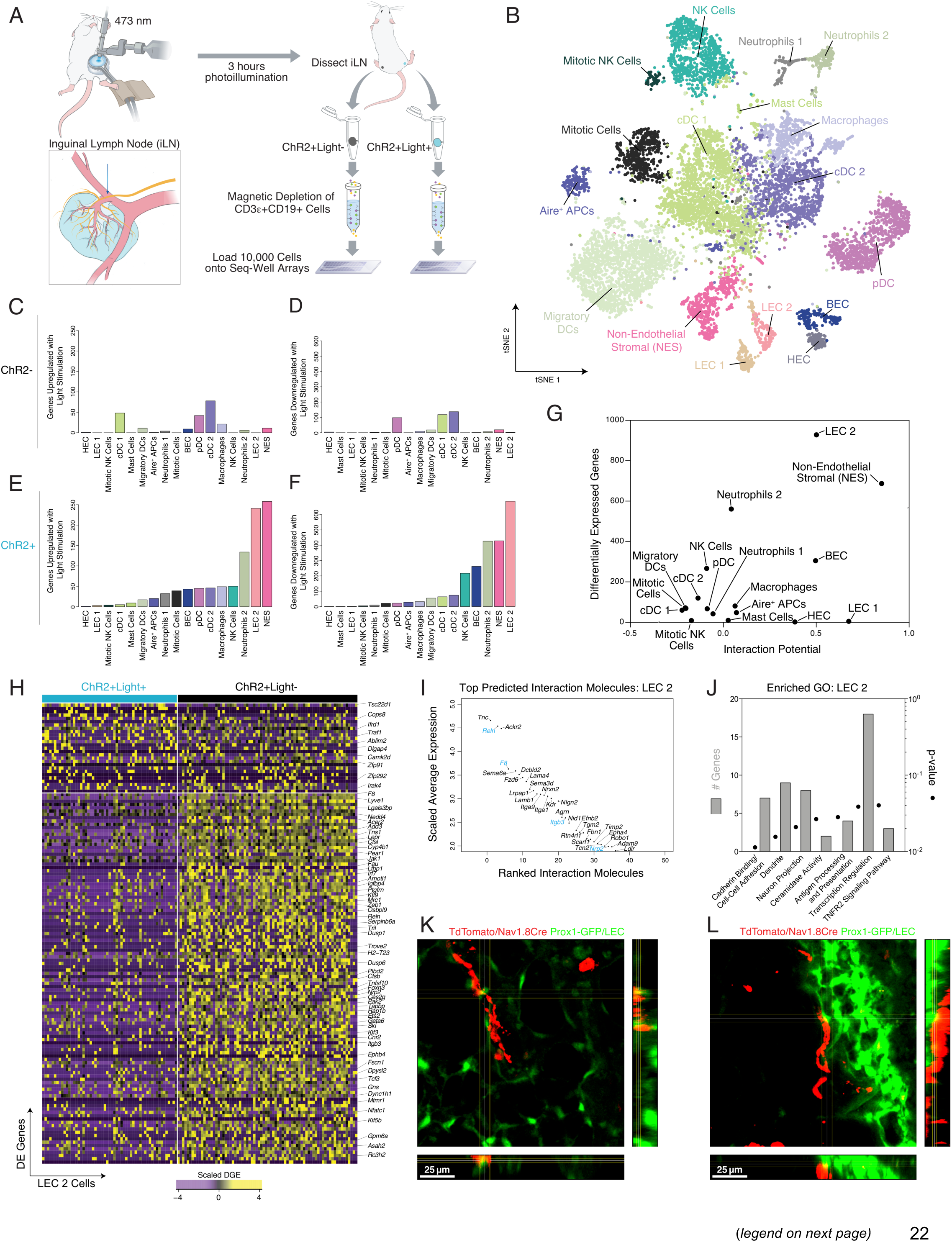
Optogenetics-assisted identification of potential postsynaptic cellular targets of LN-innervating sensory neurons in peripheral LNs. **A.** Schematic for ChR2-mediated activation of LN-innervating neurons and cell isolation protocol for scRNA-seq. **B**. tSNE visualization of cell types recovered by scRNA-seq of 10,364 cells from both light-stimulated and control LNs in ChR2+ and ChR2-animals. **C-F.** Abundance of DE genes with FDR-adjusted p-value < 0.05 and Cohen’s effect size > 0.2, separated by cell type: **C**. ChR2-(control) mice, upregulated by light stimulation; **D.** ChR2-(control) mice, downregulated by light stimulation; **E.** ChR2+ (experimental) mice, upregulated by light stimulation, omitting genes also induced in ChR2-(control) mice; and, **F.** ChR2+ (experimental) mice, downregulated by light stimulation, omitting genes also repressed in ChR2-(control) mice. **G.** Relationship between Interaction Potential and abundance of DE genes (Pearson’s r: 0.52, p = 0.03). **H.** Heatmap of DE genes between LEC 2 in light-stimulated vs. unstimulated LN in ChR2+ mice. **I**. Identity of top candidate neuron-interacting molecules in LEC 2 from steady-state LNs (Figure 6). Blue genes indicate genes that are also DE with neuronal stimulation. **J**. Enriched gene ontologies among DE genes in LEC 2 following neuronal stimulation. Left y axis: number of DE genes represented by gene ontology term; right y axis: p-value (Fisher’s Exact Test) for gene ontology enrichment. **K** and **L**, Section view of a representative two-photon micrograph of physical contact between tdTomato^+^ sensory fibers (red) and GFP^+^ LECs (green) in the medulla (**K**) and on the ceiling of SCS (**L**) of whole-mount

Following 3 hours of pulsed light exposure, iLNs from both the stimulated and unstimulated sides were processed in parallel and analyzed using Seq-Well as described above so that transcriptional changes could be tracked simultaneously in all identified cell types as a universal readout of their responses to neuronal stimulation (Figure 7A). Within a preliminary cohort, we observed negligible transcriptional changes among T and B cells, consistent with the low interaction potential between T/B cells and LN-innervating sensory neurons based on our previous anatomical and molecular characterization (**Figure S7B**). We therefore enriched for non-T and non-B cells by magnetically depleting CD3*ε*^+^ and CD19^+^ cells prior to Seq-Well analysis as described above for the steady-state iLN atlas and focused our analysis on this non-T/B cell compartment. To isolate the ChR2-dependent effect of optogenetic activation, iLNs in a separate cohort of *Nav1.8^Cre/+^* x *Rosa26^eYFP/+^* (ChR2-) animals, which expressed eYFP instead of ChR2 in Nav1.8 lineage neurons, were subjected to identical photostimulation, dissociation, cellular enrichment, and Seq-Well analysis. The changes in cellular composition and gene expression in ChR2+, but not ChR2-, animals were considered true effects of local stimulation of LN sensory fibers (**Figure S7C,** see **Methods**). Our final dataset included 4 ChR2+ mice and 3 ChR2-mice, two iLNs per mouse (one light-exposed, one control), and contained 26,887 unique genes over 10,364 cells after filtering for quality and removing residual T and B cells.

Using the same methods described above for generating the steady-state LN cell atlas (except that all T and B lineage lymphocytes were excluded), we identified a total of 17 cell types based on gene expression patterns that were in strong agreement with the diversity of LN cells described above (Figures 6B, 7B, **S7D**). Importantly, when compared to the contralateral control iLN, we did not observe significant light-induced changes in the abundance of any cell type in either ChR2+ or ChR2-animals, nor did we observe changes in LN cellularity upon light exposure (**Figures S7E and S7F**), indicating that neither the surgical/photostimulation procedures nor activation of LN-innervating sensory neurons dramatically altered the ecosystem of the exposed LNs, at least over the relatively short timescale of this analysis (3h).

To identify changes induced by ChR2-mediated neuronal excitation within each cell type, we compared gene expression between the same cell type in ChR2+ light-exposed LNs and ChR2+ control LNs. Among significantly DE genes with non-negligible effect sizes (FDR-corrected p-value < 0.05, Cohen’s d > 0.2), we filtered identified hits to remove genes with similar changes in ChR2-animals (**Table S5**). Remarkably, robust ChR2-dependent transcriptional changes were readily detected in a subset of cell types: LEC 2, BEC, NES, Neutrophils 2, and NK cells, indicating selective modulatory capabilities of LN-innervating sensory neurons (Figures 7C-7F). This ranking was independent of the effect size cutoff, and was robust to downsampling of single cells to match the abundance of cells in each cell type group (**Figure S7G**). Strikingly, we found that the magnitude of change in gene expression (as measured by number of DE genes) was significantly correlated with the overall Interaction Potential (Figure 6E) derived from the steady-state iLN atlas – i.e., the cell types predicted *in silico* to be most likely to interact with LN-innervating sensory neurons also experienced the largest magnitude transcriptional change upon experimental neuronal stimulation (Pearson’s r = 0.52, p<0.03, Figures 7G, **S7G**).

The top-impacted LN cell type was LEC 2, based on the abundance of DE genes with substantial effect sizes (**Figure S7G**). LEC 2 cells were also the cell type with the highest overall Interaction Potential (Figure 6E, 7G), and were characterized by high expression of potential molecular mediators of interaction with LN-innervating neurons in both directions, including extracellular matrix molecules (*Tnc, Fbn1, Nid1),* synaptic proteins (*Agrn, Nrxn2, Nlgn2*) (Sudhof, 2018; Wu et al., 2010), and axon guidance molecules known to regulate lymphangiogenesis (*Efnb2, Nrp2, Robo1*) (Vaahtomeri et al., 2017; Yang et al., 2010) (Figure 7I). Interestingly, we observed that some interaction-capable molecules, such as *Reln, F8, Itgb3,* and *Nrp2*, were downregulated following neuronal stimulation, suggesting a potential negative feedback loop that may act to maintain or restore homeostasis by limiting the effect of neuronal stimulation on LEC 2 once initiated (Figure 7I). Among neuronal stimulation-induced gene expression changes in LEC 2, which were dominated by downregulation, there was overrepresentation of genes in multiple pathways involved in neuronal synapses and dendrite projection, ceramidase activity, cathepsin expression, pathways involved in antigen processing and presentation, and multiple transcriptional regulators including *Gata6, Ets2, Irf7* and *Nfatc1* (Figure 7J). Interestingly, we observed a general trend toward downregulation of LEC-enriched genes including key regulators of lymphatic development and patterning, e.g., *Reln*, *Nrp2*, *Ephb4*, *Nfatc1*, *Lyve1*, as well as angiogenic molecules, e.g., *Dlg1*, *Glul*, suggesting anti-angiogenic/lymphangiogenic action of LN-innervating sensory neurons (Cho et al., 2019; Eelen et al., 2018; Kulkarni et al., 2009; Lutter et al., 2012; Vaahtomeri et al., 2017; Wu et al., 2014; Zhang et al., 2015). Moreover, downregulation of ceramidases, i.e., *Acer2* and *Asah2*, implicated in production of sphingosine-1-phosphate (S1P) in LECs, a major source of S1P in lymph, may impact lymphocyte egress from LNs, a process previously shown to be under the control of adrenergic nerves (Mao and Obeid, 2008; Nakai et al., 2014; Pappu et al., 2007; Pham et al., 2010). Consistent with the observation that LEC 2 exhibited the largest neuronal activation-evoked transcriptional changes, direct contact between sensory nerves and LECs was frequently observed in the medulla and on the ceiling of the SCS, providing further support for direct cell-cell contacts and communication between LECs and sensory innervation of LNs (Figures 7K, 7L). These data not only corroborate our analysis of Interaction Potential among candidate postsynaptic cell types, but also suggest that sensory neurons innervating LNs, when activated, may rapidly modulate the non-immune compartment to influence LN composition and/or function.

## DISCUSSION

Despite their critical roles in regulating immunological processes in barrier tissues and in the spleen (Chavan et al., 2017; Ordovas-Montanes et al., 2015), neuroimmune interactions, particularly those involving somatosensory neurons, have yet to be systematically studied in LNs. Here, we set out to directly address this gap in knowledge by analyzing the morphological, molecular, and functional attributes of the sensory neurons that innervate skin-draining LNs. We found that fibers of definitive sensory origin are preferentially located in the LN periphery, particularly the perivascular and capsular/subcapsular regions. Our data show that LN-innervating sensory neurons are a heterogeneous population with dominant peptidergic nociceptor signatures, and are molecularly distinct from their skin-innervating counterparts. We further determined that sensory neurons are most likely to interact with LN stromal cells, including BECs in microvessels other than HEV, a subset of LECs (here termed LEC 2), and NES. We first nominated these cell types *in silico* as likely interacting partners with sensory neurons based on elevated cognate receptor-ligand pair expression *in trans*. We then directly tested these predictions using selective optogenetic triggering of LN innervating neurons and observing cell-type-specific transcriptional changes within our top interacting partners. This study thus identifies a sensory neuron-stroma cell axis within skin-draining LNs as a previously unappreciated circuit of peripheral neuroimmune communication.

Previous light and electron microscopic studies had revealed the localization of peptidergic fibers within LNs (Felten et al., 1985; Fink and Weihe, 1988). However, the extent to which those fibers reflect sensory innervation has not been directly assessed. Here, we demonstrate the definitive presence of sensory innervation in LNs. Importantly, consistent with what was previously described for peptidergic fibers, we found that *bona fide* sensory fibers organized into two nerve plexuses, namely, a perivascular and a capsular/subcapsular plexus. The perivascular plexus is particularly concentrated in the medulla where preferential association with arterioles and medullary LECs is observed, whereas fibers in the capsular/subcapsular plexus are in close contact with other LN resident cells, such as SCS macrophages and LECs. Unlike traditional histologic sectioning-based approaches, our whole-mount immunolabeling protocol allows reliable identification and 3D visualization of the entire network of neuronal fibers, a prerequisite for a quantitative description of the neuronal architecture in LNs. The ability to identify LN sensory innervation through genetic labeling and retrograde tracing instead of canonical markers or ultrastructural features allowed us to perform an unbiased and complete morphological characterization of this heterogeneous neuronal population.

Our analysis revealed that sensory fibers are enriched in the LN periphery, a location prone to inflammation-induced mechanical, chemical, and cellular changes, suggesting a potential role for those neurons as local sentinels of lymph node activation. This idea is particularly attractive in light of clinical reports of painful LNs as a result of infection and cancer, suggesting functional activation of the sensory nervous system by pathological processes within LNs. Indeed, although the exact type(s) of stimulus that LN sensory neurons are tuned to and the nature of the neuronal response to diverse challenges have yet to be defined, there are multiple plausible sensing mechanisms through which LN-innervating sensory neurons could detect changes in the immune status of LNs. For example, immune challenges in the periphery often result in local production of inflammatory mediators, such as TNF*α* or IL-1β, that may become lymph-borne and directly stimulate or sensitize LN-innervating sensory neurons via their cognate cytokine receptors, i.e. TNFR1 (*Tnfrsf1a*) or IL-1R (*Il1r1*), respectively. This putative effect of pro-inflammatory cytokines on draining LNs likely represents one component of their well-documented pro-algesic role in models of inflammatory pain (Baral et al., 2019).

It is also likely that sensory fibers, particularly those embedded within the LN capsule, can sense changing mechanical forces as a result of inflammation-induced rapid LN expansion. Indeed, most DRG neurons display mechanosensitivity albeit to different degrees due to different mechanotransduction mechanisms (Drew et al., 2002; Hu and Lewin, 2006; Lewin and Stucky, 2000; McCarter et al., 1999). Notably, LN-innervating sensory neurons appear particularly sensitive to mechanical stimuli experienced in the LN periphery, since most of them express high levels of mRNA for the mechanically gated ion channel, *Piezo2*, the major mechanotransducer for touch, proprioception, baroreception and airway stretch (Nonomura et al., 2017; Ranade et al., 2014; Woo et al., 2015; Zeng et al., 2018).

There is also growing evidence for direct activation of sensory neurons by infectious pathogens. For example, nociceptors in barrier tissues have been shown to detect bacterial products, such as pore forming toxins (PFTs), N-formyl peptides and endotoxin (Blake et al., 2018; Chiu et al., 2013; Diogenes et al., 2011; Meseguer et al., 2014). Conceivably, intact microbes and/or microbial products carried by the lymph may be similarly detected by sensory fibers within LNs. Consistent with this idea, we found that LN-innervating sensory neurons expressed mRNA for LPS receptors, i.e., TLR4 and TrpA1, as well as ADAM10, the receptor for α-hemolysin, a PFT from *S. aureus* (Diogenes et al., 2011; Inoshima et al., 2011; Meseguer et al., 2014; Wilke and Bubeck Wardenburg, 2010).

The exposure of LN-innervating sensory fibers to the contents of afferent lymph also represents a potential vulnerability to incoming neurotropic pathogens. For example, a subcutaneous challenge of mice with vesicular stomatitis virus (VSV) results in rapid transport of infectious lymph-borne virions to the draining LN where VSV can invade peripheral neurons to cause a fatal ascending infection of the CNS (Iannacone et al., 2010). Viral neuroinvasion is usually prevented by specialized SCS macrophages that rapidly produce copious amounts of type I interferon upon VSV infection to prevent viral replication in adjacent neuronal fibers (Iannacone et al., 2010). Although the anatomic origin and identity of VSV susceptible nerve fibers within LNs remain to be formally established, their morphology and location in the subcapsular/capsular space are consistent with them being of sensory origin. In fact, both subunits of the type I interferon receptor (*Ifnar1* and *Ifnar2*) were abundantly expressed in LN-innervating sensory neurons. These findings imply that upon local challenge with a neurotropic virus, SCS macrophage-derived type I interferon may elicit an antiviral response within LN-innervating sensory neurons as an effective resistance mechanism.

The fact that LN-innervating sensory neurons are a heterogeneous population of DRG neurons with widely varying sizes, myelination statuses, and molecular classes suggests that individual sensory neurons will have distinct activation thresholds for the same or different stimuli. We speculate that this heterogeneity could be a means for differential triggering of local neuroimmmune circuits in different immunological contexts. This idea is further supported by our scRNA-Seq analysis, which revealed a transcriptional profile of LN-innervating sensory neurons distinct from skin-innervating sensory neurons. Although molecular differences between sensory neurons innervating different targets have been noted previously (Robinson and Gebhart, 2008; Yang et al., 2013), innervation target-dependent differences between sensory neurons have not been addressed using unbiased, transcriptome-wide analytical methods. Here, we found at the whole transcriptome level that, consistent with previously-described differential representation of peptidergic and nonpeptidergic fibers in the skin vs. visceral organs (Robinson and Gebhart, 2008), peptidergic nociceptors are overrepresented among LN-innervating sensory neurons, whereas nonpeptidergic nociceptors are enriched among the skin-innervating population from the same axial levels. In addition to preferential innervation of a specific target tissue by certain nociceptor subtypes, we also identified substantial and reproducible gene expression differences between sensory neurons of the same subtype innervating different tissues. Some of the DE genes that were highly enriched in LN neurons relative to the skin, e.g., *Ptgir* and *Prokr2*, are also known to be expressed by sensory neurons that innervate other deep tissues. Thus, it will be important to extend this comparative analysis to sensory neurons innervating other tissues in order to identify gene expression patterns that may be truly unique to LN-innervating neurons (Yang et al., 2013). Notwithstanding, we propose that target-restricted gene expression patterns may reflect tissue-specific functional or developmental requirements that partly account for the previously noted context-dependent nature of sensory neuronal regulation of inflammation and immunity (Foster et al., 2017). For example, in the skin, nociceptors can play pro-inflammatory roles in settings such as psoriasiform inflammation and candidal infection (Kashem et al., 2015; Riol-Blanco et al., 2014), or immunosuppressive roles in bacterial infections, such as with *S. pyogenes* and *S. aureus* (Chiu et al., 2013; Pinho-Ribeiro et al., 2018). Innervation target-dependent subtype composition and gene expression differences that we uncovered here suggest that sensory neuron-immune interactions may be organized in an organ-specific manner, thereby contributing to the context-dependent nature of immune regulation.

Analysis of ligand-receptor gene expression patterns in single-cell datasets has been instrumental in deconstructing the complex cellular communication network in the context of tissue function, development and cancer biology (Camp et al., 2017; Cohen et al., 2018; Kumar et al., 2018; Smillie et al., 2019; Vento-Tormo et al., 2018). The possibility of applying this powerful approach to predict neuron-non-neuronal cell interactions has only just begun to be explored (Drokhlyansky et al., 2019). Here, we demonstrate the utility of this strategy in uncovering novel neuroimmune crosstalk by systematically testing predictions from our *in silico* cellular interaction analysis using optogenetic experiments. Because this ligand-receptor interactome dataset does not take into consideration parallel interactions amongst different LN cells, it does not allow us to assign biological relevance to specific ligand/receptors pairs. Nevertheless, this strategy can reveal candidate molecular mediators of sensory neuron-non-neuronal interactions in LNs that can be experimentally tested in the future by more targeted approaches. For example, expression patterns of classic synaptogenic complexes, i.e., agrin-alpha3 Na+/K+-ATPase and neurexin-neuroligin (Hilgenberg et al., 2006; Sudhof, 2018), in LN-innervating sensory neurons and the LEC 2 subset might suggest a novel function for those classical signaling molecules in establishing synapse-like sensory neuron-LEC contacts within LNs. Additionally, we note that our intercellular interaction network was generated based on expression data from resting cells, and thus is most relevant to steady-state LNs. The sensory neuron-immune interactions in inflamed or pathologically altered LNs are potentially distinct and remain to be addressed.

Optogenetic circuit mapping has revolutionized the field of neuroscience by linking neurons to network activity and behavior (Deisseroth, 2015). The downstream output is typically measured by electrophysiology, calcium imaging and behavioral assays (Kim et al., 2017). Only recently has this approach been applied to interrogate neuronal circuits underlying neuromodulation of peripheral tissues (Chang et al., 2015; Cohen et al., 2019; Mickle et al., 2019; Murray et al., 2019; Rajendran et al., 2019; Williams et al., 2016; Zeng et al., 2015). In those few cases, optogenetics-induced effects were generally assessed based on specific hypotheses, such as select physiological or cellular functions. While this targeted approach has undoubtedly helped to reveal the complex interplay between the nervous system and peripheral tissues, its utility is more limited for addressing the cellular mechanism and extent of neuromodulation in a complex tissue, especially one with varied functional outputs, such as a LN. Here, we have begun to address this challenge by using scRNA-seq as an unbiased and high-throughput readout of neuronal influence on every major LN cell type, based on the assumption that modulatory effects of optogenetic stimulation are reflected, at least in part, at the transcriptional level in most or all possible postsynaptic target cells. Crucially, the most impacted postsynaptic targets, stromal cells, also appear to be the most likely interacting partners of LN-innervating sensory neurons based on morphological and molecular criteria. As inflammation is a probable endogenous stimulus for LN-innervating sensory neurons, inflammation-induced remodeling of LN stroma might be mediated, in part, by local sensory innervation. While the current optogenetic stimulation paradigm was specifically designed to capture immediate effects of local activation of LN sensory fibers, thus identifying potential direct non-neuronal responders, alternative modes of activation that are more amenable to temporal profiling of neuronal effects should be explored to map non-neuronal responses over a longer period of time.

Although sensory neurons by definition relay information from the periphery to the CNS, recent studies have shown that sensory neurons are not exclusively afferent, but also possess the ability to act in a motor neuron-like, or efferent, fashion via a process known as the “axon reflex” (Chiu et al., 2012; Cohen et al., 2019; Richardson and Vasko, 2002). That is, action potentials generated locally at peripheral terminals of sensory neurons can back-propagate to neighboring axonal collaterals once they reach axonal branch points. For example, in a recent study, optogenetic stimulation of cutaneous TRPV1^+^ sensory neurons was sufficient to stimulate local IL-17 mediated inflammation and enhanced anti-bacterial immune responses, which appeared to depend largely on the efferent function of the sensory neurons (Cohen et al., 2019). It remains to be determined (but seems likely) whether axon reflexes occur also in LN-innervating neurons, but it is conceivable that this mechanism could result in local release of neurotransmitters from peripheral terminals of activated LN-innervating neurons and impact the function of LN resident cells, without engaging the CNS. In addition, sensory neurons might also impact LNs through their conventional afferent functions by initiating monosynaptic or polysynaptic reflex-like neural circuits in the CNS, which, in visceral organs, culminates in motor output from the autonomic nervous system mediated by sympathetic and/or parasympathetic neurons. Both efferent and afferent functions of sensory neurons have been implicated in immunomodulation (Baral et al., 2019; Chavan et al., 2017; Cohen et al., 2019), but their relative contributions to any specific immunological process are still poorly understood. Since LNs are dually innervated by sensory and sympathetic neurons, the robust modulatory effects of selective optogenetic stimulation of LN-innervating sensory neurons observed here could in theory result from either their atypical efferent or their more conventional afferent functions, or both. The fact that we found a close correlation between the *in silico* predicted interaction potential of individual LN cell populations with LN-innervating sensory neurons and their actual response to selective optogenetic stimulation of sensory neurons supports the idea of direct efferent communication, but does not rule out additional mechanisms that may function in parallel. Further studies will be needed to dissect the circuit-level mechanisms underlying optogenetics-induced gene expression changes in this system.

The finding that stromal cell subsets followed by certain innate leukocyte populations are preferred targets of LN-innervating sensory neurons was unexpected in light of the known modes of sensory neuronal regulation of immunity in barrier tissues. The profound transcriptional changes in the stromal compartment could be due, at least in part, to prototypic vascular effects of CGRP and substance P that are well documented in barrier tissues (Baral et al., 2019; McCormack et al., 1989; Saria, 1984). However, we consider it unlikely that the newly identified connections between sensory neurons and two distinct endothelial subsets in LNs, non-HEV blood endothelial cells (BEC) and LEC 2, were directly driven by CGRP or substance P because neither EC population was found to express the cognate neuropeptide receptors. It is noteworthy in this context that the LEC 1 subset remained essentially unaffected by optogenetic stimulation of sensory neurons despite the putative juxtaneuronal positioning of LEC 1 in the subcapsular region and the fact that the predicted Interaction Potential of LEC 1 was as high as that of LEC 2, BEC and NES. Among many other possibilities, this surprising observation may suggest that LEC 1 interact with LN-innervating sensory neurons by primarily sending signal to the neurons.

The identification of the LEC 2 subset, as the top postsynaptic targets of LN-innervating sensory neurons is particularly intriguing, as local sensory afferents have been implicated in the regulation of antigen, lymph, and lymphocyte flow through LNs, processes that are controlled by LECs (Hanes et al., 2016; Moore et al., 1989). Whether and to what extent the communication between sensory neurons and LEC 2 contributes to those processes will require a better understanding of the distribution and function of LEC 2 within LNs. Using a recently-published resource describing the transcriptional diversity and identity of LECs in human LNs, we further contextualized our two murine LEC subsets (Takeda et al., 2019). Within the LEC 2 cluster, we found cells with elevated expression of *Emcn*, *Ackr2*, *Ackr4*, *Cav1*, and *Fabp4*, among other gene sets, which suggests high similarity with LECs found in the SCS ceiling, afferent and efferent lymphatic vessels, as well as the cortical sinuses. In contrast, LEC 1 cells were defined by *Madcam1* expression, as well as variable *Bmp2*, *Msr1*, *Vcam1*, *Ccl20*, and *Cxcl1* expression, supporting placement of these cells within the medullary sinuses and the SCS floor. Of note, two ceramidases, *Acer2* and *Asah2*, implicated in sphingosine-1-phosphate (S1P) biosynthesis, were downregulated by optogenetic stimulation in LEC 2, suggesting that sensory neurons may impact lymphocyte trafficking by regulating S1P-mediated lymphocyte egress from LNs (Mao and Obeid, 2008; Pappu et al., 2007; Pham et al., 2010). This effect could either reflect direct efferent communication between sensory neurons and LEC 2 or involve an efferent autonomic reflex triggered by afferent sensory signals. Consistent with the latter mechanism, sympathetic neurons were previously shown to suppress lymphocyte egress from LNs (Nakai et al., 2014). Thus, sensory and sympathetic fibers of LNs may act together or independently of each other to regulate the dwell time of recirculating lymphocytes within LNs, thus potentially fine-tuning antigen encounters and the ensuing adaptive immune responses.

Within the immune compartment, certain innate leukocytes, including neutrophils, macrophages and mast cells, were also targets of optogenetically stimulated sensory fibers, whereas the bulk of adaptive lymphocytes remained essentially unaffected. Of note, a nociceptor-neutrophil communication axis was recently identified as a major immunosuppressive mechanism in mouse models of *S. aureus* induced pneumonia and *S. pyogenes* induced necrotizing fasciitis (Baral et al., 2018; Pinho-Ribeiro et al., 2018). Whether nociceptor-neutrophil communication exerts similar immunomodulatory activities in LNs remains to be determined. Somewhat unexpectedly, interactions between sensory fibers and DCs within LNs were notably limited, at least within the experimental parameters of this study. This finding distinguishes LNs from peripheral barrier tissues where nociceptors are both necessary and sufficient to drive IL-23 production by dermal DCs resulting in type 17 inflammation (Kashem et al., 2015; Riol-Blanco et al., 2014). However, it is important to note that neither our *in silico* nor optogenetic analyses can rule out DCs or, for that matter, adaptive T or B cells as direct or indirect interacting partners of sensory neurons within LNs. In fact, DCs were among the LN cell types that express the CGRP receptor subunits at a high level. Further experimentation is warranted to uncover potential effects of LN-innervating sensory neurons on innate and/or adaptive responses to specific immune challenges.

In conclusion, we have established LNs as a point of convergence between the sensory nervous system and the immune system by identifying a molecularly distinct and heterogeneous population of sensory neurons with the capacity to impact LN function and homeostasis. We speculate that the heterogeneous, but unique composition of LN-innervating sensory neurons may be an essential feature allowing for continuous monitoring of the immune status of LNs that may vary widely in magnitude and quality based on the type of challenge. It is intriguing to hypothesize that specific immune triggers could also engage one or more subtype of LN sensory neurons with presumably distinct immunomodulatory capacities, producing customized modes of immunomodulation. Further investigation will be needed to test this hypothesis.

## ACKNOWLEDGEMENTS

We thank Guiying Cheng for technical assistance with LN injections; Dr. Constance Cepko and Dr. Ralph Adams for sharing mouse lines; Dr. Stephen Liberles’s lab for sharing the laser/shutter setup for optogenetic stimulation; Dr. David Ginty’s lab for sharing the Zeiss microscope for whole mount spinal cord and DRG imaging; the Microscopy Resources On the North Quad (MicRoN) facility for the use of confocal microcopy; the HMS Center for Immune Imaging for provision of imaging and image analysis resources; the members of the von Andrian and Shalek labs for comments and discussion on the project. S.H. was supported by the NIH (NIAMS 5R01AR068383-03), (5T32HL066987-17) and GSK postdoc fellowship. A.K.S. was supported by the Searle Scholars Program, the Beckman Young Investigator Program, the Pew-Stewart Scholars Program for Cancer Research, a Sloan Fellowship in Chemistry, the NIH (1DP2GM119419 and 2RM1HG006193), and the Ragon Institute of MGH, MIT and Harvard. J.O.M. was supported by HHMI Damon Runyon Cancer Research Foundation Fellowship (DRG-2274-16). C.G.K.Z. was supported by T32GM007753 from the National Institute of General Medical Sciences. U.v.A. was supported by the Ragon Institute of MGH, MIT and Harvard and by NIH grant AR068383.

## AUTHOR CONTRIBUTIONS

S.H., C.G.K.Z., A.K.S., and U.v.A. conceived the study. S.H. performed and analyzed *in vivo* experiments with help from J.A. and N.M. for image analysis. C.G.K.Z., with the help of J. O.-M. and M. V., generated scRNA-seq data. C.G.K.Z. analyzed scRNA-seq data. S.H., C.G.K.Z., A.K.S., and U.V.A. interpreted the results. S.H., C.G.K.Z., A.K.S., and U.v.A. wrote manuscript with input from J. O.-M.

## MATERIALS AND METHODS

### Experimental model and subject details

Mouse lines used in this study were all previously described and include *Nav1.8^Cre^* (RRID:IMSR_EM:04582) (Nassar et al., 2004), *Rosa26^LSL-tdTomato^* (RRID:IMSR_JAX:007914), *Bmx-CreER^T2^* (MGI:5513853) (Ehling et al., 2013), *Rosa26^LSL-DTA^* (RRID:IMSR_JAX:009669), *Prox1-EGFP* (MGI:4847348) (Choi et al., 2011), *Rosa26^LSL-ChR2-eYFP^* (RRID:IMSR_JAX:024109), *Rosa26^LSL-eYFP^* (RRID:IMSR_JAX:007903) and *ChAT^BAC^-eGFP* (RRID:IMSR_JAX: 007902). All of the animals were handled according to approved institutional animal care and use committee (IACUC) protocols of Harvard Medical School. Unless indicated otherwise, adult mice of both sexes between 6-12 weeks of age were used for various experiments.

### Whole mount immunohistochemistry

Whole mount immunohistochemistry of LNs was performed using an iDISCO protocol with methanol pretreatment optimized for LNs (Renier et al., 2014). Briefly, adult animals (6-12 weeks) were perfused with 25 mL of PBS (Hyclone) and 25 mL of 4% paraformaldehyde (PFA, Sigma) sequentially at room temperature (RT). Peripheral lymph nodes (PLNs), including popliteal and inguinal lymph nodes (popLNs and iLNs), were postfixed with 4% PFA for 1 hr at 4°C. For methanol pretreatment, fixed LNs were washed sequentially in 50% methanol (Fisher Scientific) (in PBS) for 1 hr, 100% methanol for 1 hr, 50% methanol for 1 hr, PBS for 1 hr twice, and PBS/0.2% Triton X-100 (VWR) for 1 hr twice at RT. LNs were then left in PBS/0.2% Triton X-100/20% DMSO (Sigma)/0.3 M glycine (BioRad) overnight at RT and blocked in PBS/0.2% Triton X-100/10% DMSO/6% donkey serum (Jackson Immunoresearch) or goat serum (Gibco)/anti-CD16/CD32 (Fc block) (Bio X cell) overnight at RT. LNs were subsequently washed in PBS/0.2% Tween-20 (Fisher Scientific)/10 μg/mL heparin (Sigma) (PTwH), for 1 hr twice at RT, before incubation with antibody mix in PTwH/5% DMSO/3% donkey or goat serum/Fc block 1:100 for 3 days at RT. LNs were extensively washed in PTwH for at least 6 times over the course of a day at RT. For unconjugated antibodies, LNs were further incubated with a secondary antibody mix including a panel of species-specific anti-IgG (H+L) Alexa Fluro 488, 546, 647 and 594-conjugated antibodies (Invitrogen or Jackson Immunoresearch) in PTwH/5% DMSO/3% donkey or goat serum/Fc block 1:100 for 3 more days at RT. LNs were washed in the same way as after primary antibody incubation for 1 day. Immunolabeled LNs following one round of antibody incubation for conjugated antibodies (or two for unconjugated antibodies) were then processed for clearing, which includes sequential incubation with 50% methanol for 1 hr, 100% methanol for 1 hr for three times and a mixture of 1-part benzyl alcohol (Sigma): 2-parts benzyl benzoate (Sigma) (BABB) overnight at RT. For tdTomato immunolabeling, goat anti-mCherry antibody (ACRIS) was preabsorbed against PLNs from *tdTomato^-^* animals overnight at RT prior to use.

Whole mount immunohistochemistry of DRGs and the skin was performed as described previously (Li et al., 2011). Briefly, DRGs inside vertebral column and the depilated hairy skin from PFA-perfused animals (6-12 weeks) were postfixed with 4% PFA for 1 hr or Zamboni fixative (Fisher Scientific) overnight, respectively at 4°C. Samples were washed every 30 min with PBS/0.3% Triton-100 (0.3% PBST) for 4-6 hr, then incubated with primary antibodies in antibody diluent (0.3% PBST/20% DMSO/5% donkey or goat serum) for 2-3 days at RT. Samples were then washed with 0.3% PBST every 30 min for 5–8 hr before incubation with secondary antibodies in antibody diluent for 2-3 days at RT. After extensive washes as described above, samples were dehydrated and cleared in 50% methanol for 1 hr, 100% methanol for 1 hr for three times and BABB overnight at RT.

Cleared whole mount tissues were imaged in BABB between two coverglasses using Olympus FV3000 confocal imaging system, except for those shown in Figures 7K and 7L, which were acquired on BioRad 2100MP system and those shown in Figures 3B, **S3B** and **S3E**, which were acquired on Zeiss Stereo Discovery V16. The antibodies used were: rabbit anti-CGRP (Immunostar, 24112, 1:500), chicken anti-GFP (Aves Labs, GFP-1020, 1:500), chicken anti-NF200 (Aves Labs, NFH, 1:500), rabbit anti-Tyrosine Hydroxylase (Millipore, AB152, 1:500), goat anti mCherry antibody (1:500, ACRIS AB0040-200), rabbit anti-βIII-Tubulin (Biolegend, 802001, 1:500), Alexa Fluor 647-conjugated rat anti-CD31 (Biolegend, 102416, 1:50), FITC–conjugated mouse anti-smooth muscle actin (aSMA) (Sigma, F3777-.2ML, 1:500), eFluor 660-conjugated mouse anti-smooth muscle actin (aSMA) (Thermo Fisher, 50-9760-82, 1:100), eFluor 660-conjugated rat anti-CD169 (Thermo Fisher, 50-5755-80, 1:50), Pacific Blue-conjugated rat anti-CD45 (Biolegend, 103126, 1:50), Alexa Fluor 488-conjugated rat anti-PNAd (Thermo Fisher, 53-6036-82, 1:50)

### Retrograde labeling of LN- and skin-innervating neurons

To retrogradely label LN-innervating neurons, adult animals (6-12 weeks) were anesthetized by intraperitoneal injection of ketamine (Patterson Vet) (50 mg kg^-1^) and xylazine (Patterson Vet) (10 mg kg^-1^). The skin overlying the targeted iLN was shaved and depilated so that the LN underneath was visible percutaneously. A 5 mm incision was made directly on top of the iLN. The iLN was microdissected without perturbing afferent lymphatic vessels and surrounding blood vessels. 1 μl of Adeno-Associated Virus (AAV) (AAV2/1.CMV.HI.eGFP-Cre.WPRE.SV40, titer >=8E+12 vg/mL, Addgene) mixed with 0.5 μl of fast green (Sigma) was injected into the iLN of *Rosa26^LSL-tdTomato/LSL-tdTomato^* animals using a pulled and trimmed glass pipette (FHC) which was connected to a 5 mL syringe through the aspiration assembly system (Sigma). The injection site was immediately rinsed with 2 mL of saline (Patterson Vet) to wash away any off-target virus before the incision was closed with sutures. Animals were sacrificed between 1 month and 6 months after injection for histology or scRNA-seq. To directly visualize the axonal projections of sensory neurons retrogradely labeled from the iLN, AAV carrying Cre-dependent tdTomato cassette (AAV2/1.CAG.Flex.tdTomato.WPRE.bGH, titer ≥10^13^ vg/mL, Addgene) was injected into the iLN *Nav1.8^Cre/+^* animals as described above. For WGA-based retrograde labeling, 1 μl of WGA-AF488 (2 mg/mL in PBS, Invitrogen) was injected into the iLN of *Nav1.8^Cre/+^*; *Rosa26^LSL-tdTomato/+^* animals as described before and the animals were processed for histology 4 days post injection. Retrograde labeling of skin-innervating neurons was described previously (Kuehn et al., 2019). Briefly, following ketamine-xylazine mediated anesthesia, a single injection of 0.2 μl of various AAV2/1 viruses as described above and 0.1 μl of fast green was delivered using the injection device described above intradermally into the patch of depilated skin overlying the iLN of adult mice (6-12 weeks). Animals were sacrificed between 1 month and 6 months after injection for immunohistochemistry, RNAscope, or scRNA-seq.

### Immunohistochemistry of tissue sections

Adult animals (6-12 weeks) were perfused with 25 mL of PBS and 25 mL of 4% PFA sequentially at RT. The intact vertebral column was postfixed overnight with 4% PFA at 4°C. DRGs were subsequently dissected and processed for cryosectioning. 14 μm serial cryosections were collected and processed for immunohistochemistry as described previously (Li et al., 2011). In brief, sections were postfixed with 4% PFA for 10 min at RT. Following three washes with PBS, they were incubated with blocking buffer (PBS with 5% normal goat serum and 0.3% Triton-100) for 1 hr at RT. The sections were then incubated with Rabbit anti-TH (Millipore) in the same blocking buffer overnight at 4°C. The following day, sections were washed three times with wash buffer (PBS with 0.3% Triton-100) before incubation with goat Alexa Fluor 647-conjugated anti-rabbit (Invitrogen) for 1 hr at RT. Sections were then washed for three times with wash buffer before mounting in Fluoromount Aqueous Mounting Medium (Sigma). WGA-488 and tdTomato were visualized directly based on endogenous fluorescence. All the sections with tdTomato^+^ cells were imaged at 20x using Olympus FV3000 confocal imaging system.

### Intravital two-photon microscopy

Adult *Nav1.8^Cre/+^*; *Rosa26^LSL-tdTomato/+^* animals (6-12 weeks) were given 1 μg of FITC-conjugated rat anti-CD169 antibody (BioRad) diluted in a total volume of 20 μl of PBS into the right footpad to label CD169^+^ subscapular macrophages inside the draining LN. Immediately after, the animals were prepared microsurgically for intravital two-photon microscopy as described before (Mempel et al., 2004). Briefly, anesthesia during surgical preparation and imaging was achieved through the ketamine-xylazine method as described above. The right popLN was exposed and positioned with the cortex facing outwards with minimal perturbation to afferent lymphatic vessels and surrounding blood vessels, while the animal was immobilized onto a custom-built stage by its hip bone and the vertebral column. The imaging chamber was created around the exposed LN with high vacuum grease (VWR) on the side and a coverslip on top. A thermocouple (Omega) was placed next to the LN to monitor the local temperature, which was maintained between 36.5 and 37°C by a custom-built water bath heating system. Two-photon imaging was performed on a Bio-Rad Radiance 2100MP Confocal/Multiphoton microscopy system with two MaiTai Ti:sapphire lasers (Spectra-Physics) tuned to 800 nm and 900 nm for two photon excitation and second harmonic generation. Z-stacks of sensory innervation of the capsular/subcapsular space on the cortical side were acquired in 1 μm steps with a 20×, 0.95 numerical aperture objective (Olympus).

### Manual cell sorting for scRNA-seq

Adult mice with retrogradely-labeled LN- or skin-innervating neurons were sacrificed by CO_2_ asphyxiation. T13 and L1 DRGs ipsilateral to the side of injection were quickly removed without nerves attached and checked for tdTomato labeling in cold HBSS (1X, no Ca^2+^ or Mg^2+^) (VWR) under Leica MZ10 F stereomicroscope with fluorescence. DRGs were immediately digested with 1 mL of papain solution (HBSS/10 mM HEPES (VWR)/500 μM EDTA (Westnet)/0.4 mg/mL L-Cysteine (Sigma)/1.5 mM CaCl_2_ (Sigma)/20 unit/mL Papain (Worthington)) in a 37°C water bath for 10 min, with agitation every 2 min. DRGs were further digested with 1 mL of collagenase type II/dispase solution (HBSS/10 mM HEPES/4 mg/mL collagenase type II (Worthington)/5 mg/mL dispase (Thermo Fisher)) in a 37°C water bath for 30 min, with agitation every 10 min. Following centrifugation at 400 g for 4 min, digested DRGs were mechanically disrupted in 0.2 mL of complete L15 medium (L15 (Invitrogen)/10 mM HEPES/10% FBS (Germini)) by passing them first through a 1000 µL pipette tip up to 10 times, and then through a 200 µL pipette tip up to 5 times until the tissues were fully dissociated. To remove myelin/axonal debris, the cell suspension diluted in 1 mL of complete L15 medium was carefully layered on top of 5 mL of Percoll gradient (L15/10 mM HEPES/20% Percoll (GE Healthcare) and centrifuged at 400 g for 9 min. After removing the supernatant, cells were washed in 2 mL of L15/10 mM HEPES and centrifuged at 750 g for 3 min. Finally, cells were resuspended in 1 mL of cold sorting buffer (L15/10 mM HEPES/1 mg/mL BSA (VWR)/25 μg/mL DNase I (Roche)), and subjected to fluorescence-assisted single-cell picking as described previously (Hempel et al., 2007). Briefly, the cell suspension diluted in 3 mL of sorting buffer was immediately transferred to a 35 mm petri dish (Scanning dish) with lane markings 6 mm apart and let sit on ice until most cells had settled to the bottom which normally takes 15-20 min. Rare fluorescent cells were readily identified under Leica MZ10 F stereomicroscope with fluorescence (transillumination off) by scanning the bottom of the dish lane by lane to maximize recovery and avoid rescanning. Zoom was set such that the field of view corresponded to the width of a single lane. To pick out fluorescent cells with minimal contamination from nonfluorescent cells, a pulled and trimmed micropipette (World Precision Instruments) was carefully lowered under transillumination into the sorting buffer until it was in the vicinity of the target cell. Simultaneous positive pressure was applied by mouth through the aspiration assembly system, as described above for retrograde labeling. Once the micropipette was in position, the target cell was gently aspirated into the micropipette through capillary action by transient release of positive pressure. The micropipette was quickly removed to prevent aspiration of unwanted cells or debris. The content of the micropipette, including the target cell, was expelled gently into a droplet of cold fresh sorting buffer on a different 35 mm petri dish (wash dish 1) under transillumination. Wash dish 1 was kept on ice while subsequent scans for fluorescent cells occurred. Once 16 or all the fluorescent cells, whichever comes first, were collected in wash dish 1, cells were washed two additional times by moving them one by one into a new droplet of sorting buffer on clean 35 mm petri dishes. Micropipettes were not reused for different cells to avoid cross contamination. After the final wash, each fluorescent cell was pipetted up and down the micropipette three times to remove unwanted contamination before being ejected into 10 μl of cold RLT (Qiagen) supplemented with 1% β-mercaptoethanol (Sigma) in a 96-well plate, and snap-frozen on dry ice and stored at −80°C. The entire manual sorting procedure was routinely completed in 1.5 hr.

### scRNA-seq of neurons using Smart-Seq2

Single-cell libraries were generated according to the SMART-seq2 protocol (Picelli et al., 2014; Trombetta et al., 2014). Briefly, RNA from single-cell lysates was purified using AMPure RNA Clean Spri beads (Beckman Coulter) at a 2.2x volume ratio, and mixed with oligo-dT primer (SMART-seq2 3’ Oligo-dT Primer), dNTPs (NEB), and RNase inhibitor (Fisher Scientific) at 72°C for 3 minutes on a thermal cycler to anneal the 3’ primer to polyadenylated mRNA. Reverse transcription was carried out in a master mix of Maxima RNaseH-minus RT enzyme and buffer (Fisher Scientific), MgCl_2_ (Sigma), Betaine (Sigma), RNase inhibitor, and a 5’ template switch oligonucleotide (SMART-seq2 5’ TSO) using the following protocol: 42°C for 90 minutes, followed by 10 cycles of 50°C for 2 minutes, 42°C for 2 minutes, and followed by inactivation at 70°C for 15 minutes. Whole transcriptome amplification was achieved by addition of KAPA HiFi HotStart ReadyMix (Kapa Biosystems) and IS PCR primer (ISPCR) to the reverse transcription product and amplification on a thermal cycler using the following protocol: 98°C for 3 minutes, followed by 21 cycles of 98°C for 15 seconds, 67°C for 20 seconds, 72°C for 6 minutes, followed by a final 5-minute extension at 72°C. Libraries were purified using AMPure XP SPRI beads at a volume ratio of 0.8x followed by 0.9x.

Library size was assessed using a High-Sensitivity DNA chip (Agilent Bioanalyzer), confirming the expected size distribution of ∼1000-2000 bp. Tagmentation reactions were carried out with the Nextera XT DNA Sample Preparation Kit (Illumina) using 250 pg of cDNA per single cell as input, with modified manufacturer’s instructions as described. Libraries were purified twice with AMPure XP SPRI beads at a volume ratio of 0.9x, size distribution assessed using a High Sensitivity DNA chip (Agilent Bioanalyzer) and Qubit High-Sensitivity DNA kit (Invitrogen). Libraries were pooled and sequenced using NextSeq500/550 High Output v2 kits (75 cycles, Illumina) using 30-30 paired end sequencing with 8-mer dual indexing.

### RNAscope

The RNAscope Fluorescent Multiplex Assay (ACD Biosystems) was performed according to RNAscope Multiplex Fluorescent Reagent Kit v2 user manual for fresh-frozen tissue samples. Briefly, 14 μm fresh frozen sections from T13 and L1 DRGs with each side containing retrogradely-labeled tdTomato^+^ LN- or skin-innervating neurons from the same animal were hybridized with RNAscope probes for *Ptgir* (487851), *tdTomato* (317041-C2), and *Prokr2* (498431-C3) simultaneously. The probes were amplified and detected with TSA plus fluorescein, cyanine 3 and cyanine 5 (Perkin Elmer). The ACD 3-plex negative control probe was run in parallel on separate sections in each experiment to assess the background level and set the acquisition parameter. All sections with *tdTomato*^+^ cells were imaged at 20x using an Olympus FV3000 confocal imaging system. The frequency of *Ptgir*^+^ or *Prokr2*^+^ DRG neurons among the *tdTomato*^+^ LN- or skin-innervating population was determined by considering all the *tdTomato*^+^ cells that were recovered and uniquely-defined from a single animal.

### Tamoxifen treatment

Tamoxifen (Sigma) was dissolved in corn oil (Sigma) at a concentration of 20 mg/mL by shaking overnight at 37°C, and stored at 4°C for the duration of the injections. For labeling arterial vessels with *Bmx-CreER^T2^*, 0.5 mg of tamoxifen was delivered intraperitoneally into *Bmx-CreER^T2^*; *Rosa26^eYFP/+^* animals between 4-6 weeks of age daily for three consecutive days. Animals were analyzed between 1-3 weeks later.

### 6-OHDA treatment

For sympathetic denervation, the stock solution of 6-hydroxydopamine (6-OHDA) (Sigma) was prepared in water at 42 mg/mL and stored at −20°C. *Nav1.8^Cre/+^*; *Rosa26^LSL-tdTomato/+^* animals from the same litter between the ages of 6-12 weeks were injected intraperitoneally with 6-OHDA (100 mg kg^-1^) or an equal volume of saline daily for 5 consecutive days. Animals were analyzed the following day.

### Optogenetic stimulation of iLN-innervating sensory neurons

Age-matched adult *Nav1.8^Cre/+^*; *Rosa26^LSL-ChR2-eYFP/+^* (ChR2+) or *Nav1.8^Cre/+^*; *Rosa26^LSL-eYFP/+^* (ChR2-) animals (6-12 weeks) were deeply anesthetized (isoflurane, 1.5%–2%, Patterson Vet) maintained at normal body temperature with a water bath heating system (Baxter) during surgical preparation and photostimulation. The animals were surgically prepared for intravital optogenetic stimulation using a method that was adapted from a previously-described protocol for intravital microcopy of iLNs (von Andrian, 1996). Briefly, the skin with the left iLN was flipped inside out following a small incision immediately left to the midline and glued onto a metal block to keep the medulla side of LN exposed. Care was taken not to overstretch the skin flap and damage lymphatic and blood vessels. The site of illumination, the branch point of the antero-posterior-running segment of the y-shaped superficial epigastric artery from where LN feeding arterioles emerged was located and exposed with microdissection without compromising the blood vessel integrity while the tissue was kept moist with normal saline. The stimulation chamber was then built around the iLN with vacuum grease on the side to keep solution from leaking, as well as a metal hairpin shaped tubing with hot water flowing inside on top of vacuum grease to maintain the tissue between 36.5 and 37°C. A thermocouple was placed next to the branch point to monitor the temperature at the tissue. An optic fiber (200 µm core, Thorlabs) coupled to a DPSS laser light source (473 nm, Shanghai Laser & Optics Century) was positioned for focal illumination directly on top of the branch point. The stimulation chamber was subsequently filled to the metal tubing with GenTeal Tears Lubricant Eye Gel (Alcon) to keep the tissue from drying out during stimulation. Pulsed light stimulation (5 m pulses, 125 mW/mm^2^ intensity, 20 Hz) was delivered to the targeted region for 3 hr under the control of a shutter system (Uniblitz). iLNs from both sides were immediately removed after light stimulation and kept in ice cold LN media (HBSS (Corning)/2% FBS/10 mM HEPES/2 mM CaCl_2_) until subsequent processing.

### LN Dissociation and Single Cell Isolation

LNs were kept on ice until processing, < 60 minutes between animal sacrifice and tissue digestion. LN media was aspirated, and each LN was placed in 1 mL of pre-warmed digestion media (0.8 mg/mL dispase, 0.2 mg/mL collagenase P (Roche), 50 µg/mL of DNase I in LN media). Using a pair of needle-nose forceps, the capsule of each LN was gently pierced, and the LN in digestion media were placed in a 37°C water bath for 20 minutes with no agitation. Next, LNs were gently agitated without touching the tissue, pelleted by gravity, and the 1 mL of digestion media supernatant was removed and placed in a collection tube on ice containing 10 mL of quenching buffer (PBS/5 mM EDTA/5% FBS). A fresh 1 mL of digestion buffer was added to each LN, and the LNs were placed back in the 37°C water bath for an additional 5 minutes. The LN was gently agitated and triturated using a 1000 µL pipette tip, solid capsular and stromal matter was allowed to settle to the bottom of the tube without centrifuging, and the supernatant digestion media was added to the same collection tube containing quenching buffer. 5-minute incubation periods in fresh digestion buffer and trituration with a 1000 µL pipette tip continued until LNs were completely digested, typically requiring 3-4 additional digestion steps. The cellular suspension in quenching buffer was filtered through a 100 µm filter, and washed with an additional 15 mL of quenching buffer. Single-cell suspensions were centrifuged at 300g for 3 minutes at 4°C, and counted using a hemocytometer and light microscope. We recovered an average of 4.00 +/- 0.53 million cells per LN, and observed no differences in cellularity by treatment group or animal genotype. We saved an aliquot of 60,000 cells from each sample in quenching media on ice as the unenriched sample, and centrifuged the remaining cells at 300 g for 3 minutes at 4°C. Next, using the Miltenyi CD3*ε* microbead kit and CD19 mouse microbead kit, all remaining LN cells were stained according to manufacturer instructions with the following modifications. First, single cells were stained with CD3*ε* biotin for 10 minutes on ice, washed once with MACS buffer (PBS/0.5% BSA (Sigma)/2 mM EDTA) and stained simultaneously with CD19 microbeads and biotin microbeads. Cells were isolated using LD columns (Miltenyi) according to manufacturer specifications and the flow-through was collected as the non-T and non-B enriched sample. Single cells from both enriched and unenriched samples were pelleted by centrifugation at 300g for 3 minutes at 4°C, and counted using a hemocytometer with trypan blue staining to estimate cell viability. Across 14 LNs, we recovered an average of 270,000 +/- 31,000 (mean +/- SEM) cells per lymph node following CD3*ε* and CD19 depletion with > 90% viability.

For LN cellularity analysis, single-cell suspensions of the two iLNs from the same ChR2+ or ChR2-mouse (6-12 weeks) were prepared as above. The cells were then filtered through steel mesh and resuspended at the appropriate cell density in FACS buffer before being acquired on a BD *Accuri*™ *C6 Plus* flow cytometer (BD Biosciences).

### LN scRNA-seq using Seq-Well

Single cells from each lymph node prior to and post CD3*ε* and CD19 depletion were kept separate and diluted to 15,000 cells in 200 µL complete media (RPMI 1640/10% FBS). Seq-Well was performed as described with changes noted below (Aicher et al., 2019; Gierahn et al., 2017). Briefly, a pre-functionalized PDMS array containing ∼86,000 nanowells was loaded with uniquely-barcoded mRNA capture beads (ChemGenes) (Macosko et al., 2015) and suspended in complete media for at least 20 minutes. 15,000 cells were deposited onto the top of each PDMS array and let settle by gravity into distinct wells. The array was gently washed with PBS, and sealed using a functionalized polycarbonate membrane with a pore size of 0.01 µm, which allows exchange of buffers without permitting mixing of cell materials between different wells. Seq-Well arrays were sealed in a dry 37°C oven for 40 minutes, and submerged in a lysis buffer containing 5 M guanidium thiocyanate (Sigma), 1 mM EDTA, 1% beta-mercaptoethanol and 0.05% sarkosyl (Sigma) for 20 minutes at room temperature. Arrays were transferred to hybridization buffer containing 2 M NaCl (Fisher Scientific) with 8% (v/v) polyethylene glycol (PEG, Sigma) and agitated for 40 minutes at room temperature, mRNA capture beads with mRNA hybridized were collected from each Seq-Well array, and beads were resuspended in a master mix for reverse transcription containing Maxima H Minus Reverse Transcriptase and buffer, dNTPs, RNase inhibitor, a 5’ template switch oligonucleotide (Seq-Well 5’ TSO), and PEG for 30 minutes at room temperature, and overnight at 52°C with end-over-end rotation. Exonuclease digestion was carried out as described previously: beads were washed with TE with 0.01% tween-20 (Fisher Scientific) and TE with 0.5% SDS (Sigma), denatured while rotating for 5 minutes in 0.2 mM NaOH, and resuspended in ExoI (NEB) for 1 hour at 37°C with end-over-end rotation (Hughes et al., 2019). Next, beads were washed with TE + 0.01% tween-20, and second strand synthesis was carried out by resuspending beads in a master mix containing Klenow Fragment (NEB), dNTPs, PEG, and the dN-SMRT oligonucleotide (Seq-Well Second Strand Primer) to enable random priming off of the beads. PCR was carried out as described using 2X KAPA HiFi Hotstart Readymix and ISPCR primer (Seq-Well ISPCR), and placed on a thermal cycler using the following protocol: 95°C for 3 minutes, followed by 4 cycles of 98°C for 20 seconds, 65°C for 45 seconds, 72°C for 3 minutes, followed by 12 cycles of 98°C for 20 seconds, 67°C for 20 seconds, 72°C for 3 minutes, followed by a final 5-minute extension at 72°C. Post-whole transcriptome amplification proceeded as described above for SMART-seq2 libraries, with the following exceptions: AMPure XP SPRI bead cleanup occurred first at a 0.6 x volume ratio, followed by 0.8x. Library size was analyzed using an Agilent Tapestation hsD5000 kit, confirming the expected peak at ∼1000 bp, and absence of smaller peaks corresponding to primer. Libraries were quantified using Qubit High-Sensitivity DNA kit and prepared for Illumina sequencing using Nextera XT DNA Sample Preparation kit using 900 pg of cDNA library as input to tagmentation reactions. Amplified final libraries were purified twice with AMPure XP SPRI beads as before, with a volume ratio of 0.6x followed by 0.8x. Libraries from 3 Seq-Well arrays were pooled and sequenced together using a NextSeq 500/550 High Output v2 kit (75 cycles) using a paired end read structure with custom read 1 primer (Seq-Well CR1P): read 1: 20 bases, read 2: 50 bases, read 1 index: 8 bases.

### Image analysis

All image analyses were performed in Imaris 9.2.1 or 7.4.2 as detailed below. To better visualize neuronal architecture in or/and around LNs, for all LN images except for Figures 7K, 7L, **S1B, S2D, S2E, S3A, S3D** and **S7A**, an isosurface for the LN was generated by manually drawing LN contours on 2D slices every fifth slice and was used to mask the original images so that only what was inside the LN mask was shown. Depending on the purpose of the experiment, LN isosurfaces were defined with varying degrees of stringency: based on the outermost layer of LECs in Figures 2A, 2C **and S2A**, on collagen type I staining in Figure 2F, on SMA staining in Figures 2D, 2E and **S2B**, or on GFP background staining in **Figure S1A** or on tdTomato background staining everywhere else. In Figures 2A, 2C, 2D, 2E, 2F, **S2A S2B** and **S3H**, additional masking of the channel(s) where nerves were stained was performed with isosurfaces generated for neuronal signal within LNs based on morphology, i.e. fiber-like structures that can be traced through multiple slices, to highlight neuronal structures. To better visualize fibers in the capsular/subcapsular space of LNs as shown in Figure 2F, intranodal sensory fibers and total sensory fibers within and below the capsule were isolated by masking the original channel with LN isosurfaces defined based on GFP (LECs) and collagen type I staining, respectively. The resulting channel after subtracting the former channel from the latter one corresponds to the capsular/subcapsular plexus. Original, processed images and rendered isosurfaces were viewed as 3D reconstructions in surpass view with orthogonal camera setting unless indicated otherwise.

For quantification of innervation density of LNs as in Figures 1C-1F, relevant channels were first masked with the LN isosurface as described above. Isosurfaces for sensory and sympathetic fibers within the masked channels, i.e., inside the LN, were then generated by automatic creation based on features that distinguish neuronal signal from everything else, e.g., intensity, sphericity, followed by manual editing. Sensory or sympathetic fiber density for a given LN was defined as the ratio of the volume of isosurfaces for sensory or sympathetic fibers within the LN to that for the LN.

For quantification of penetration depth of intranodal sensory fibers, the outermost layer of LECs, which demarcates the LN boundary, was used to precisely segment LNs into isosurfaces. Isosurfaces for intranodal sensory fibers, sensory fibers within the relevant channel after applying the LN isosurface as a mask, were generated as described above. Using the distance transformation function, the closest distance from any given voxel within the LN isosurface to the surface of the LN in μm was computed and converted into an intensity value for that given voxel in a separate channel. To determine penetration depth of intranodal sensory fibers, the distance transformation channel was masked against isosurfaces for intranodal sensory fibers to generate a new channel where the penetration depth at any given voxel within the intranodal sensory fibers was encoded as the intensity value for that specific voxel with the maximum intensity value representing the maximum penetration depth for a given LN. Such a channel, when displayed in surpass view as in Figure 2A, allowed direct visualization of the spatial relationship between intranodal sensory fibers and the nearest LN surface. Additionally, the penetration depth of intranodal sensory fibers was described in Figure 2B in the form of the percentage of total intranodal fibers found within LN spaces with increasing distance away from the LN surface. For that analysis, the original distance transformation channel, as described above, was used to create a series of isosurfaces of decreasing sizes which represent increasingly-deep LN spaces with its closest distance to the LN surface increasing from 0 to 100 μm with 10 μm intervals. For example, 10 was set as the intensity threshold cutoff during automatic creation so that all voxels with intensity value larger than and equal to 10 were selected in one single surface which corresponds to the LN space 10 μm and more below the LN surface. To calculate the percentage of total intranodal sensory fibers in any of those LN spaces, the isosurface for total sensory fibers and that for a said LN space, e.g., 10 μm and more below the surface, as described above, were each used to generate their corresponding binary channels, where all voxels outside of a surface were set as 0, while those inside were set at 100. Colocalization analysis was then performed on those two binary channels, and the percentage of non 0 voxels in the binary nerve channel that were colocalized corresponded to the percentage of total intranodal sensory fibers found 10 μm and more below the LN surface. This process was repeated for increasingly smaller LN spaces with 10 μm interval. The percentage of total intranodal fibers in LN spaces in the form of 10um bins from 0 to 100 μm, e.g., 10-20 μm, was further derived from serial subtraction of the percentage in the LN space that is 10um deeper, e.g., (20 μm and more below the surface), from that in the current LN space, e.g., (10 μm and more below the surface), as shown in Figure 2B.

### Neuron scRNA-seq data preprocessing

Single cells were sequenced to a depth of 1.6 +/- 0.1 million (mean +/- SEM) reads per cell. Pooled libraries were demultiplexed using bcl2fastq (v2.17.1.14) with default settings, and aligned using STAR (Dobin et al., 2013) to the mouse UCSC genome reference (version mm10), and a gene expression matrix was generated using RSEM (v1.2.3) in paired-end mode. Single-cell libraries with fewer than 3,000 unique genes and fewer than 17% of reads mapping to transcriptomic regions were excluded from subsequent analysis, resulting in a final dataset of 52 LN-innervating neurons collected from 8 mice, and 31 skin-innervating neurons collected from 4 mice. Among cells retained for analysis, the number of unique genes captured was 9,843 +/- 229 (mean +/- SEM) among LN-innervating neurons and 9,653 +/- 302 among skin-innervating neurons. Libraries from LN-innervating neurons contained 50.45 +/- 2.3% transcriptome-aligning fragments, libraries from skin-innervating neurons contained 58.33 +/- 2.9 %. Among all alignment and library quality metrics assessed, we found no significant differences between LN-innervating and skin-innervating neurons (see **Figures S4A-S4C**). All analysis of gene expression was completed using the normalized RSEM output as transcripts per million (TPM).

### Neuron scRNA-seq differential gene expression

All analysis of scRNA-seq data was carried out using the R language for Statistical Computing. Single-cell libraries were first assessed for expression of canonical neuronal markers and known lineage-defining genes from accompanying imaging data, such as Nav1.8 (*Scn10a*) and tyrosine hydroxylase (*Th*). The full list of markers is supplied in **Table S2**. To directly assess differences in gene expression between LN-innervating and skin-innervating neurons, we used the R package Single Cell Differential Expression (SCDE, version 1.99.1) with default input parameters (Fan et al., 2016; Kharchenko et al., 2014). A cutoff of Holm corrected Z score >1.96 or < −1.96 (corresponding to a corrected p-value < 0.05) was used to identify significantly DE genes for subsequent analysis. Heatmaps were created using the R package gplots (version 3.0.1). DAVID was used for analysis of over-represented gene ontologies over significantly DE genes (Huang da et al., 2009a, b).

### Analysis of neuron scRNA-seq with Usoskin, Furlan et al. Sensory Neuron Atlas

As our target-specific single cells do not represent the full diversity of neurons contained in the DRG, we utilized the scRNA-seq atlas published by Usoskin, Furlan et al. *Nature Neuroscience* 2015 (subsequently referred to as the “Sensory Neuron Atlas”) to classify our cells (Usoskin et al., 2015). Using the raw data and accompanying metadata hosted at http://linnarssonlab.org/drg/, we first identified the intersection of expressed genes from the Sensory Neuron Atlas and LN-innervating and skin-innervating single cells, and eliminated cells identified as non-neuronal (“NoN” and “NoN outlier”) from the Sensory Neuron Atlas, resulting in a dataset of 148 neurofilamentous (NF), 81 peptidergic (PEP), 251 tyrosine hydroxylase (TH), 169 non-peptidergic (NP), and 39 “Central, unsolved” cells. To mimic the dimensionality reduction methods the previous authors used to identify major neuronal cell types, we transformed the data as log_2_(1+TPM), and calculated the gene variance across all cells. We cut to genes with a variance log_2_(1+TPM) > 0.5, resulting in 11,778 genes. Next, we performed principal component analysis over the log_2_-transformed, mean-centered data, and found that PC_2_ and PC_4_ reflected major axes of variability between TH, PEP, NF, and NP cell types – identified by the authors of the previous study as “Level.1” cell type subsets (Figure 4A). To identify how LN-innervating and skin-innervating cells related to major DRG cell types in a reduced dimensional space, we projected our target-specific data into PC_2_ and PC_4_ of the Sensory Neuron Atlas. This was completed by first calculating the principal components of the Sensory Neuron Atlas:

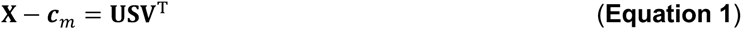

where **X** is the log_2_(1+TPM) data matrix of M genes by N cells from the Sensory Neuron Atlas. Equation 1 calculates the singular value decomposition of this matrix after subtracting the average of each row (gene) of **X**, denoted ***c_m_***, from **X**. **U** represents a matrix of M orthonormal vectors corresponding to M genes and **V** represents a matrix of N orthonormal vectors corresponding to N cells. To apply this same dimensionality reduction transformation to our new dataset of LN-innervating and skin-innervating single cells, **Y**, we use Equation 2:

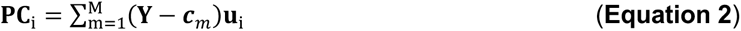

**Y** represents the log_2_(1+TPM) transformed matrix of our innervation-target-specific data, **c_m_** refers to the same vector of row (gene) averages calculated from **X**. The centered **Y** matrix is multiplied as a dot product with the i^th^ principal component gene eigenvector, or the i^th^ column vector of **U**, denoted **u**_i_. By taking the sum over all transformed rows for each column (cell), we project the LN-innervating and skin-innervating data (**Y**) into the principal component space calculated for the Sensory Neuron Atlas (**X**), denoted **PC**_i_. This data is visualized by plotting the PC_2_ and PC_4_ vectors from the Sensory Neuron Atlas (transparent circles, Figure 4A), with the PC_2_ and PC_4_ vectors from the transformed LN-innervating and skin-innervating cells (filled squares). The Euclidean distance between each innervation-target-specific single cell and all cells within the Sensory Neuron Atlas was calculated over PC2 and PC4 (Figure 4B). The range of cell-to-cell Euclidean distances between like-cells (e.g. PEP-to-PEP) within the Sensory Neuron Atlas is represented by a dashed line corresponding to the 99%ile.

To analyze the expression similarity between each single cell from our target-specific dataset and the Sensory Neuron Atlas subtypes in a more directed, supervised manner, we assessed how each single cell correlated with each subtype of Sensory Neuron Atlas. We elected to use the more detailed neuronal subtypes defined by Usoskin, Furlan, et al., termed “Level.3”, which breaks some of the major neuron subtypes, NP, PEP, and NF, into subtypes based on intra-population diversity. We calculated the average gene expression for each neuron subtype (e.g. NP1) over the log_2_(1+TPM) transformed single-cell data, generating pseudo-population averages for each Usoskin, Furlan-defined “Level.3” neuron subtype. Next, we only considered genes in our pseudo-population averages that were designated as “subtype-defining” by the Usoskin, Furlan et al. analysis, corresponding to the top 50 genes upregulated within each cell type when compared to all other cell types in their Sensory Neuron Atlas, yielding 379 unique genes. We similarly restricted our LN-innervating and skin-innervating single-cell libraries to only these 379 unique genes, and calculated the Spearman correlation between each target-specific single cell (following log_2_(1+TPM) transformation) and the Sensory Neuron Atlas pseudo-population averages (Figure 4C). We clustered LN-innervating and skin-innervating single cells by their correlation with each Sensory Neuron Atlas pseudo-population using complete linkage clustering, and using a cut height of 0.8 retained 4 distinct Neuron Types: Neuron type 1 “PEP1-like” (LN-innervating cells: 25, skin-innervating cells: 9), Neuron type 2 “NP-like” (LN-innervating cells: 1, skin-innervating cells: 14), Neuron type 3 “mixed PEP/NF123” (LN-innervating cells: 23, skin-innervating cells: 5), and Neuron type 4 “mixed PEP2/NF12345” (LN-innervating cells: 3, skin-innervating cells: 3) (Figure 4C).

To assess the gene expression phenotype of each Neuron Type, we used SCDE to identify DE genes between cells of each Neuron Type compared to all cells of the 3 remaining Neuron Types. SCDE was run as described above with default input parameters, genes with a Holm-corrected p-value < 0.01 were considered significant and presented in Figure 4E and **Table S2**.

### LN Seq-Well data preprocessing

Reads were aligned and processed according to the Drop-Seq Computational Protocol v2.0 (https://github.com/broadinstitute/Drop-seq). Briefly, reads were first demultiplexed according to index read 1 using bcl2fastq (v2.17.1.14) with default settings. Read 1 was split into the first 12 base pairs corresponding to the cell barcode (CB), and the 13-20^th^ base pairs, which encode the unique molecular identifier (UMI). CBs, UMIs, and read 2 sequences with low base quality were discarded, as were any that contained non-random sequences (e.g. primer sequences, poly-A tails). Following CB and UMI tagging, read 2 was aligned to the mouse genome (version mm10) using STAR v2.5.2b with default parameters including “--limitOutSJcollapsed 1000000 --twopassMode Basic”. STAR alignments were merged to recover cell and molecular barcodes, and any sequences within hamming edit distance 1 were merged, as these likely originated from the same original sequence. Additional methods to correct for bead synthesis errors in the CB or UMI are detailed in the Drop-Seq Computational Protocol v2.0 (“DetectBeadSynthesisErrors” function). Digital gene expression matrices for each array were retained following quality filtering and UMI-correction, and further processed using the R language for Statistical Computing. Cells with fewer than 300 unique genes were removed from analysis.

### Dimensionality reduction, clustering, visualization, and cell type identification of LN Seq-Well data

We restricted our primary analysis of LN-resident cell types to only arrays corresponding to steady state inguinal LN without surgical manipulation or optogenetic stimulation. A total of 9,662 cells were retained with 25,929 unique genes expressed across 7 mice with 1 LN per mouse (**Table S3**). For 2 mice, we sequenced arrays corresponding to all LN cells prior to CD3*ε*/CD19 depletion as well as CD3 *ε* /CD19 depleted cells on a separate array. The average cell recovery per array was 1,074 +/- 141 (mean +/- SEM) cells, with an average gene count of 1,581 +/- 11 genes and average UMI per cell of 4,251 +/- 48 UMI (mean +/- SEM). Data was normalized and scaled using the Seurat R package (https://github.com/satija.lab/seurat) (Butler et al., 2018): transforming the data to log_e_(UMI+1) and applying a scale factor of 10,000. We confirmed equivalent depth and cell quality across each of our arrays and the absence of major batch effects introduced by sequencing work-up day or other technical factors, and thus did not regress any batch-related covariates out of our data, including individual cell quality or mitochondrial percent. To identify major axes of variation within our data, we first subsetted our data to only highly-variable genes across all cells – all genes with dispersion (calculated as the variance to mean ratio) > 1.1 were kept, resulting in 2,348 variable genes. Principal component analysis was applied to the cells cut to variable genes for the top 100 principal components. Using the JackStraw function within Seurat, we identified the top significant PCs, and compared these significant PCs to the variance explained by each dimension, ultimately choosing 41 PCs for subsequent clustering and further dimensionality reduction (Shekhar et al., 2016). Critically, we completed all of the following analysis over a range of variable gene cutoffs and principal components to ensure that our cell identification results were robust to parameter choice.

For 2D visualization, we used the Barnes-Hut implementation of t-distributed stochastic neighbor embedding (t-SNE) with “perplexity” set to 40. This tSNE projection of the steady state LN atlas is represented in Figure 6B, **S5A**. To identify clusters of transcriptionally-similar cells, we employed unsupervised clustering with the Louvain algorithm with the Jaccard correction (Blondel et al., 2008). Briefly, this method involves constructing a k-nearest neighbor graph over the Euclidean distance between cells in the 41-PC reduced space, followed by a shared nearest neighbor (SNN)-based clustering and modularity optimization (Waltman and Van Eck, 2013). We implemented this using the FindClusters tool within the Seurat R package with default parameters and k.param set to 20 and resolution set to 0.4. Here, we intentionally underclustered our data to avoid erroneously splitting cells with shared cell type functions, as the variable genes calculated for this dimensionally-reduced space likely did not fully reflect more nuanced cell type differences (e.g. variable behavior between Neutrophil subtypes). The “Parent Cluster” results from first-pass cell type clustering are represented in the tSNE plot and clusters identified in **Figure S5A**. We used the Seurat function FindAllMarkers to identity differentially-expressed genes upregulated within each cluster compared to all other cells in the dataset, and tested differential expression using the likelihood-ratio test for single-cell gene expression (by setting test.use to “bimod”) (McDavid et al., 2013). The top 100 differentially-expressed genes for each cluster were analyzed, as ranked by the average fold change and restricted to only those with FDR-corrected p-values < 0.05. Next, to assess if any cell subtypes existed within each cluster, we restricted our data to only cells within a single “Parent Cluster”, and recalculated the variable genes over these cells. The above analysis, from calculation of variable genes to tSNE visualization and cluster identification, was repeated for each cluster listed in **Figure S5A**. Cell types for which we could identify sub-clusters with significant differentially-expressed genes are marked with asterisks next to their names in **Figure S5A**, and the sub-cluster tSNE projections and top differentially expressed genes are represented in **Figures S5B-S5O**. For the T cell parent cluster, we required two iterative sub-clustering steps to fully enumerate all constituent cell types: the first clustering step differentiated regulatory T cells (Tregs) from the remaining T cells (**Figure S5B, S5C**), and subsequent clustering on the non-Treg T cells uncovered CD4 T cells vs. CD8 T cells. All differentially-expressed genes within each sub-cluster can be found in **Table S4**.

After exhaustive assessment for cell subclusters within each cell type, we identified 24 unique cell types within our steady state dataset (Figure 6B). We calculated the differentially expressed genes between each cell type and all other cells using a likelihood ratio test (using the FindAllMarkers function with test.use set to “bimod”), the results of this analysis are presented in Figures 6C, **S5P**, and **Table S4**. By identifying canonical marker genes within these DE gene lists from the literature and using resources such as ImmGen (Heng et al., 2008), we attributed cell identities to each cell type within our dataset, as named in Figures 6B, 6C, and **S5P**.

### Analysis of cellular receptor-ligand pairs

We reasoned that cells or cell types within the LN that interact with innervating neurons would likely express proteins that enable such contact or communication. As we generated unbiased single-cell transcriptomic data from LN-innervating neurons and the potential targeted cell types, we incorporated databases of ligand and receptor pairs to understand if any of the LN-resident cell types expressed a high abundance of cognate molecules, and would thus be poised to interact with innervating neurons. A general schematic of this method is provided in **Figure S6A**. We used the database of receptor-ligand interactions curated by Ramilowski et al (Ramilowski et al., 2015), which consists of 2,422 total interactions over 708 unique genes (originally provided as human genes, and converted to mouse orthologs using the HUGO database). First, data from LN-innervating neurons was limited to only genes with non-negligible expression, using a cutoff of average log_2_(1+TPM) > 3, yielding 6,666 total genes for subsequent analysis. The intersection of genes within the Ramilowski interaction database and those expressed at non-negligible levels among LN-innervating neurons yielded 184 total genes. After limiting to only interactions with at least one participating gene expressed in the LN-innervating neurons, the interaction database was restricted to 750 total receptor-ligand pairs, and 471 unique potential cognates. We next assessed the expression of these 471 cognate genes within the LN-resident cell atlas. First, we summarized the expression of individual cells within the LN-resident atlas by taking the pseudo-population average of each cell type (over non-log single-cell data). We limited the LN-resident atlas data to only genes with non-negligible expression across all cell type pseudo-populations, cutting to genes with an average UMI expression > 1, yielding 256 total potential cognates (from the previous 471). Next, we developed a summary statistic to reflect the abundance of neuron cognates expressed within LN-resident cell types. First, we scaled our data by subtracting the mean and dividing by the standard deviation for each individual gene – this enabled us to assess the contribution of all genes equally such that signal was not dominated by genes with high total expression (Figure 6E). Finally, we calculated the “Interaction Potential”

(IP) as the mean of these scaled values for each cell type: cell types that expressed relatively higher abundances of all candidate neuron-cognates received a higher IP score. Our null model states that the interaction potentials we calculated are no more extreme than the IP we would have recovered by chance. To test our experimentally-derived IP, we generated a null distribution by shuffling the cell type labels over all single cells within the LN-resident cell atlas, and repeated the “cell type” averaging, scaling, and IP calculation for 1,000 permutations. By comparing our true IP scores to the null distribution, we were able to identify certain cell types with significantly higher IP than observed by chance, and could attribute a P-value to each cell type (Figure 6F, 99% confidence interval denoted by dashed line). IP scores were re-scaled such that the lower bound of the 99% confidence interval was equal to 0 for clarity. The results of this approach are presented in Figures 6D-6F, **S6A, S6B.**

Crucially, we were concerned that the method of calculation of the IP, the summary statistics applied, the choice of raw vs. scaled data, or confounding factors that differentiate cell types, including average genes/cell and number of cells per cell type, would influence our ranked list of top interacting cell types and bias our results. For example, we wondered whether differences in quality metrics or other technical factors between cell types might result in higher or lower IP rankings – for instance, a cell type with significantly higher RNA recovery per cell than another cell type would appear to have a higher interaction potential. We found no correlation between the IP (as reported in Figure 6F) and the median UMI per cell for each cell type (**Figure S6B,** p = 0.32). To address bias introduced by our choice in summary statistic or data normalization, we repeated the above pipeline without gene-wise scaling across cell types (**Figure S6C**), or by calculating the percent of cells with non-zero expression of a given gene, in the place of calculating of average expression per cell type (**Figure S6D**). In both of these cases, we observed that non-endothelial stroma, LEC 1, LEC 2, BEC, and HEC remained the top-scoring cell types for Interaction Potential (significance calculated by permutation test as described above). Finally, we reasoned that variations in the number of cells per cell type might limit our ability to compare between different cell types. We iteratively down-sampled our single-cell data to analyze interaction potentials (using the method in Figures 6D-6F) for only 25 total cells per cell type – the histograms of these calculations after 1,000 iterations are plotted in **Figure S6E**. Critically, non-endothelial stroma, LEC 1, LEC 2, BEC, and HEC cell types remained top-ranking in Interaction Potential after controlling for cell abundance per cell type.

Finally, we derived an alternative statistical testing strategy to assess the overrepresentation of neuron-interaction cognates among expressed genes between different cell types. Here, we binarized our data to classify genes as “expressed” or “not expressed” within a cell type, using an average gene expression cutoff of 1. We considered the list of 256 potential neuronal cognate genes, and used a Fisher’s Exact Test to assess whether the cognate gene list was overrepresented among expressed genes for a given cell type (mimicking the field-standard for gene ontology enrichment analysis) (Huang da et al., 2009a, b), and a Holm correction to adjust for multiple tests. In close agreement with the results from our interaction potential statistic above, we found significant overrepresentation of potential neuronal cognate genes in the following cell types (listed in decreasing statistical significance): non-endothelial stroma (p = 1.6 x 10^-28^), BEC (p = 2.5 x 10^-22^), LEC 1 (p = 4.5 x 10^-22^), HEC (p = 8.3 x 10^-21^), LEC 2 (p = 9.6 x 10^-20^), Macrophages (p = 8.7 x 10^-9^), Mast Cells (p = 6.5 x 10^-8^), Neutrophils 2 (p = 5.2 x 10^-6^), Neutrophils 1 (p = 1.8 x 10^-4^), pDC (p = 1.7 x 10^-3^), Aire^+^ APCs (p = 3.4 x 10^-3^), and cDC2 (8.9 x 10^-3^). All other cell types were non-significant by a Holm-adjusted p-value cutoff of 0.01. Critically, this ranking was not sensitive to the choice of binarization cutoff, tested over a range of 0.5 – 10 UMI, data not shown).

### Differential gene expression following optogenetic stimulation

Cells were partitioned into the cell types annotated in Figure 7B. Using the Seurat function DiffExpTest, which employs a likelihood ratio test to identify differentially expressed genes, we analyzed cells for each cell type from ChR2+Light+ LN vs. ChR2+Light-LN. Similarly, we identified differentially expressed genes by cell type between ChR2-Light+ LN vs. Chr2-Light-LN. We reasoned that the DE genes in ChR2+ mice represented both the effects of neuronal stimulation, as well as changes induced by surgery and/or phototoxicity, while the DE genes in the ChR2-mice only correspond to changes due to surgery and/or phototoxicity. For each cell type, we identified genes DE in ChR2+ animals by a Holm-adjusted p-value cutoff of 0.05, and eliminated genes from these lists that were also DE (using the same cutoff) in ChR2-LN. We calculated the effect size using Cohen’s d, and restricted our gene lists to only those genes with a non-negligible effect size, using a cutoff of 0.2 (analysis the effect of various effect-size cutoffs in **Figure S7F**). The results of these analyses for each cell type can be found in **Table S5**. In Figure 7H, we further restricted our DE gene lists for heatmap visualization, and in Figure 7J for gene ontology analysis (using DAVID, as described above) by only including genes that were also DE between LNs harvested from the same mouse in at least 2 of 4 ChR2+ mice.

### Statistical testing

Using prism software, we performed unpaired two-tailed Student’s t-tests for Figures 1E, 1H and Figure 5F, Welch’s t-tests for Figures S1G and S3C, 2-way ANOVA with Sidak’s multiple comparisons test with for **Figure S7E**. All other statistical tests corresponding to differential gene expression or assessment of interaction potential are described above and completed using R language for Statistical Computing. Tests of correlation and correlation significance are annotated by the correlation model used (Pearson vs. Spearman) were completed using R language for Statistical Computing. Parameters such as sample size, number of replicates, number of independent experiments, measures of center, dispersion, and precision (mean +/- SEM) and statistical significances are reported in Figures and Figure Legends. A p-value less than 0.05 was considered significant unless otherwise reported; a more stringent cutoff of 0.01 was used in some instances, and annotated as such. Where appropriate, a Holm correction was used to account for multiple tests, as noted in the figure legends or methods.

## Data and software availability

The raw data and gene expression matrices for all scRNA-seq data have been deposited in the Gene Expression Omnibus (https://www.ncbi.nlm.nih.gov/geo/) under the accession number GSE139600 for SMART-seq2 of LN-innervating or skin-innervating neurons, and accession number GSE139658 for Seq-Well of LN-resident cells. All raw data from LN-innervating and skin-innervating neuron scRNA-seq can be found in **Table S1**. Gene lists corresponding to differential expression tests in Figures 4E, 5A, 5B, **S4C-S4E** can be found in **Table S2**. Raw data and cell type-identifying gene lists corresponding to Figures 6B, 6C, **S5A-S5P** can be found in **Tables S3** and **S4**. All differentially expressed genes from optogenetic stimulation experiments presented in Figure 7C-7H can be found in **Table S5**. All R code for analysis available upon request.

## REFERENCES

1. Abrahamsen, B., Zhao, J., Asante, C.O., Cendan, C.M., Marsh, S., Martinez-Barbera, J.P., Nassar, M.A., Dickenson, A.H., and Wood, J.N. (2008). The cell and molecular basis of mechanical, cold, and inflammatory pain. Science 321, 702–705.

2. Adams, R.H., and Eichmann, A. (2010). Axon guidance molecules in vascular patterning. Cold Spring Harb Perspect Biol 2, a001875.

3. Aicher, T.P., Carroll, S., Raddi, G., Gierahn, T., Wadsworth, M.H., 2nd, Hughes, T.K., Love, C., and Shalek, A.K. (2019). Seq-Well: A Sample-Efficient, Portable Picowell Platform for Massively Parallel Single-Cell RNA Sequencing. Methods Mol Biol 1979, 111–132.

4. Andoh, T., Nishikawa, Y., Yamaguchi-Miyamoto, T., Nojima, H., Narumiya, S., and Kuraishi, Y. (2007). Thromboxane A2 induces itch-associated responses through TP receptors in the skin in mice. J Invest Dermatol 127, 2042–2047.

5. Andrae, J., Gallini, R., and Betsholtz, C. (2008). Role of platelet-derived growth factors in physiology and medicine. Genes Dev 22, 1276–1312.

6. Assas, B.M., Pennock, J.I., and Miyan, J.A. (2014). Calcitonin gene-related peptide is a key neurotransmitter in the neuro-immune axis. Front Neurosci 8, 23.

7. Baral, P., Udit, S., and Chiu, I.M. (2019). Pain and immunity: implications for host defence. Nat Rev Immunol.

8. Baral, P., Umans, B.D., Li, L., Wallrapp, A., Bist, M., Kirschbaum, T., Wei, Y., Zhou, Y., Kuchroo, V.K., Burkett, P.R., et al. (2018). Nociceptor sensory neurons suppress neutrophil and gammadelta T cell responses in bacterial lung infections and lethal pneumonia. Nat Med 24, 417–426.

9. Bellinger, D.L., Lorton, D., Felten, S.Y., and Felten, D.L. (1992). Innervation of lymphoid organs and implications in development, aging, and autoimmunity. Int J Immunopharmacol 14, 329–344.

10. Belvisi, M.G. (2002). Overview of the innervation of the lung. Curr Opin Pharmacol 2, 211–215.

11. Blake, K.J., Baral, P., Voisin, T., Lubkin, A., Pinho-Ribeiro, F.A., Adams, K.L., Roberson, D.P., Ma, Y.C., Otto, M., Woolf, C.J., et al. (2018). Staphylococcus aureus produces pain through pore-forming toxins and neuronal TRPV1 that is silenced by QX-314. Nat Commun 9, 37.

12. Bley, K.R., Hunter, J.C., Eglen, R.M., and Smith, J.A. (1998). The role of IP prostanoid receptors in inflammatory pain. Trends Pharmacol Sci 19, 141–147.

13. Blondel, V.D., Guillaume, J., Lambiotte, R., and Lefebvre, E. (2008). Fast unfolding of communities in large networks. J Stat Mech P10008.

14. Brierley, S.M., Jones, R.C., 3rd, Gebhart, G.F., and Blackshaw, L.A. (2004). Splanchnic and pelvic mechanosensory afferents signal different qualities of colonic stimuli in mice. Gastroenterology 127, 166–178.

15. Buettner, M., and Bode, U. (2012). Lymph node dissection--understanding the immunological function of lymph nodes. Clin Exp Immunol 169, 205–212.

16. Butler, A., Hoffman, P., Smibert, P., Papalexi, E., and Satija, R. (2018). Integrating single-cell transcriptomic data across different conditions, technologies, and species. Nat Biotechnol 36, 411–420.

17. Camp, J.G., Sekine, K., Gerber, T., Loeffler-Wirth, H., Binder, H., Gac, M., Kanton, S., Kageyama, J., Damm, G., Seehofer, D., et al. (2017). Multilineage communication regulates human liver bud development from pluripotency. Nature 546, 533–538.

18. Caterina, M.J. (2003). Vanilloid receptors take a TRP beyond the sensory afferent. Pain 105, 5–9.

19. Chang, R.B., Strochlic, D.E., Williams, E.K., Umans, B.D., and Liberles, S.D. (2015). Vagal Sensory Neuron Subtypes that Differentially Control Breathing. Cell 161, 622–633.

20. Chavan, S.S., Pavlov, V.A., and Tracey, K.J. (2017). Mechanisms and Therapeutic Relevance of Neuro-immune Communication. Immunity 46, 927–942.

21. Chiu, I.M., Heesters, B.A., Ghasemlou, N., Von Hehn, C.A., Zhao, F., Tran, J., Wainger, B., Strominger, A., Muralidharan, S., Horswill, A.R., et al. (2013). Bacteria activate sensory neurons that modulate pain and inflammation. Nature 501, 52–57.

22. Chiu, I.M., Hehn, C., and Woolf, C.J. (2012). Neurogenic inflammation and the peripheral nervous system in host defense and immunopathology. Nat Neurosci 15.

23. Cho, C., Wang, Y., Smallwood, P.M., Williams, J., and Nathans, J. (2019). Dlg1 activates beta-catenin signaling to regulate retinal angiogenesis and the blood-retina and blood-brain barriers. Elife 8.

24. Choi, I., Chung, H.K., Ramu, S., Lee, H.N., Kim, K.E., Lee, S., Yoo, J., Choi, D., Lee, Y.S., Aguilar, B., et al. (2011). Visualization of lymphatic vessels by Prox1-promoter directed GFP reporter in a bacterial artificial chromosome-based transgenic mouse. Blood 117, 362–365.

25. Cohen, J.A., Edwards, T.N., Liu, A.W., Hirai, T., Jones, M.R., Wu, J., Li, Y., Zhang, S., Ho, J., Davis, B.M., et al. (2019). Cutaneous TRPV1(+) Neurons Trigger Protective Innate Type 17 Anticipatory Immunity. Cell 178, 919–932 e914.

26. Cohen, J.N., Tewalt, E.F., Rouhani, S.J., Buonomo, E.L., Bruce, A.N., Xu, X., Bekiranov, S., Fu, Y.X., and Engelhard, V.H. (2014). Tolerogenic properties of lymphatic endothelial cells are controlled by the lymph node microenvironment. PLoS One 9, e87740.

27. Cohen, M., Giladi, A., Gorki, A.D., Solodkin, D.G., Zada, M., Hladik, A., Miklosi, A., Salame, T.M., Halpern, K.B., David, E., et al. (2018). Lung Single-Cell Signaling Interaction Map Reveals Basophil Role in Macrophage Imprinting. Cell 175, 1031–1044 e1018.

28. Cordeiro, O.G., Chypre, M., Brouard, N., Rauber, S., Alloush, F., Romera-Hernandez, M., Benezech, C., Li, Z., Eckly, A., Coles, M.C., et al. (2016). Integrin-Alpha IIb Identifies Murine Lymph Node Lymphatic Endothelial Cells Responsive to RANKL. PLoS One 11, e0151848.

29. Deisseroth, K. (2011). Optogenetics. Nat Methods 8, 26–29.

30. Deisseroth, K. (2015). Optogenetics: 10 years of microbial opsins in neuroscience. Nat Neurosci 18, 1213–1225.

31. Diogenes, A., Ferraz, C.C., Akopian, A.N., Henry, M.A., and Hargreaves, K.M. (2011). LPS sensitizes TRPV1 via activation of TLR4 in trigeminal sensory neurons. J Dent Res 90, 759–764.

32. Dobin, A., Davis, C.A., Schlesinger, F., Drenkow, J., Zaleski, C., Jha, S., Batut, P., Chaisson, M., and Gingeras, T.R. (2013). STAR: ultrafast universal RNA-seq aligner. Bioinformatics 29, 15–21.

33. Drew, L.J., Wood, J.N., and Cesare, P. (2002). Distinct mechanosensitive properties of capsaicin-sensitive and - insensitive sensory neurons. J Neurosci 22, RC228.

34. Drokhlyansky, E., Smillie, C.S., Wittenberghe, N.V., Ericsson, M., Griffin, G.K., Dionne, D., Cuoco, M.S., Goder-Reiser, M.N., Sharova, T., Aguirre, A.J., et al. (2019). The enteric nervous system of the human and mouse colon at a single-cell resolution BioRxiv.

35. Eelen, G., Dubois, C., Cantelmo, A.R., Goveia, J., Bruning, U., DeRan, M., Jarugumilli, G., van Rijssel, J., Saladino, G., Comitani, F., et al. (2018). Role of glutamine synthetase in angiogenesis beyond glutamine synthesis. Nature 561, 63–69.

36. Ehling, M., Adams, S., Benedito, R., and Adams, R.H. (2013). Notch controls retinal blood vessel maturation and quiescence. Development 140, 3051–3061.

37. Evrard, M., Kwok, I.W.H., Chong, S.Z., Teng, K.W.W., Becht, E., Chen, J., Sieow, J.L., Penny, H.L., Ching, G.C., Devi, S., et al. (2018). Developmental Analysis of Bone Marrow Neutrophils Reveals Populations Specialized in Expansion, Trafficking, and Effector Functions. Immunity 48, 364–379 e368.

38. Fan, J., Salathia, N., Liu, R., Kaeser, G.E., Yung, Y.C., Herman, J.L., Kaper, F., Fan, J.B., Zhang, K., Chun, J., et al. (2016). Characterizing transcriptional heterogeneity through pathway and gene set overdispersion analysis. Nat Methods 13, 241–244.

39. Felten, D.L., Felten, S.Y., Bellinger, D.L., and Lorton, D. (1992). Noradrenergic and peptidergic innervation of secondary lymphoid organs: role in experimental rheumatoid arthritis. Eur J Clin Invest 22 *Suppl 1*, 37–41.

40. Felten, D.L., Felten, S.Y., Carlson, S.L., Olschowka, J.A., and Livnat, S. (1985). Noradrenergic and peptidergic innervation of lymphoid tissue. J Immunol 135, 755s–765s.

41. Fink, T., and Weihe, E. (1988). Multiple neuropeptides in nerves supplying mammalian lymph nodes: messenger candidates for sensory and autonomic neuroimmunomodulation? Neurosci Lett 90, 39–44.

42. Fletcher, A.L., Malhotra, D., Acton, S.E., Lukacs-Kornek, V., Bellemare-Pelletier, A., Curry, M., Armant, M., and Turley, S.J. (2011). Reproducible isolation of lymph node stromal cells reveals site-dependent differences in fibroblastic reticular cells. Front Immunol 2, 35.

43. Foster, S.L., Seehus, C.R., Woolf, C.J., and Talbot, S. (2017). Sense and Immunity: Context-Dependent Neuro-Immune Interplay. Front Immunol 8, 1463.

44. Gautron, L., Sakata, I., Udit, S., Zigman, J.M., Wood, J.N., and Elmquist, J.K. (2011). Genetic tracing of Nav1.8-expressing vagal afferents in the mouse. J Comp Neurol 519, 3085–3101.

45. Gierahn, T.M., Wadsworth, M.H., 2nd, Hughes, T.K., Bryson, B.D., Butler, A., Satija, R., Fortune, S., Love, J.C., and Shalek, A.K. (2017). Seq-Well: portable, low-cost RNA sequencing of single cells at high throughput. Nat Methods 14, 395–398.

46. Godinho-Silva, C., Cardoso, F., and Veiga-Fernandes, H. (2019). Neuro-Immune Cell Units: A New Paradigm in Physiology. Annu Rev Immunol 37, 19–46.

47. Gonzalez-Castillo, C., Ortuno-Sahagun, D., Guzman-Brambila, C., Pallas, M., and Rojas-Mayorquin, A.E. (2014). Pleiotrophin as a central nervous system neuromodulator, evidences from the hippocampus. Front Cell Neurosci 8, 443.

48. Green, D.P., Limjunyawong, N., Gour, N., Pundir, P., and Dong, X. (2019). A Mast-Cell-Specific Receptor Mediates Neurogenic Inflammation and Pain. Neuron 101, 412–420 e413.

49. Hanes, W.M., Olofsson, P.S., Talbot, S., Tsaave, T., Ochani, M., Imperato, G.H., Levine, Y.A., Roth, J., Pascal, M.A., Foster, S.L., et al. (2016). Neuronal circuits modulate antigen flow through lymph nodes. Bioelectron Med 3, 18–28.

50. Hempel, C.M., Sugino, K., and Nelson, S.B. (2007). A manual method for the purification of fluorescently labeled neurons from the mammalian brain. Nat Protoc 2, 2924–2929.

51. Heng, T.S., Painter, M.W., and Immunological Genome Project, C. (2008). The Immunological Genome Project: networks of gene expression in immune cells. Nat Immunol 9, 1091–1094.

52. Hilgenberg, L.G., Su, H., Gu, H., O’Dowd, D.K., and Smith, M.A. (2006). Alpha3Na+/K+-ATPase is a neuronal receptor for agrin. Cell 125, 359–369.

53. Homeister, J.W., Thall, A.D., Petryniak, B., Maly, P., Rogers, C.E., Smith, P.L., Kelly, R.J., Gersten, K.M., Askari, S.W., Cheng, G., et al. (2001). The alpha(1,3)fucosyltransferases FucT-IV and FucT-VII exert collaborative control over selectin-dependent leukocyte recruitment and lymphocyte homing. Immunity 15, 115–126.

54. Hu, J., and Lewin, G.R. (2006). Mechanosensitive currents in the neurites of cultured mouse sensory neurones. J Physiol 577, 815–828.

55. Huang da, W., Sherman, B.T., and Lempicki, R.A. (2009a). Bioinformatics enrichment tools: paths toward the comprehensive functional analysis of large gene lists. Nucleic Acids Res 37, 1–13.

56. Huang da, W., Sherman, B.T., and Lempicki, R.A. (2009b). Systematic and integrative analysis of large gene lists using DAVID bioinformatics resources. Nat Protoc 4, 44–57.

57. Hughes, T.K., Wadsworth, M.H., Gierahn, T.M., Do, T., Weiss, D., Andrade, P.R., Ma, F., de Andrade Silva, B.J., Shao, S., Tsoi, L.C., et al. (2019). Highly Efficient, Massively-Parallel Single-Cell RNA-Seq Reveals Cellular States and Molecular Features of Human Skin Pathology. bioRxiv, 689273.

58. Iannacone, M., Moseman, E.A., Tonti, E., Bosurgi, L., Junt, T., Henrickson, S.E., Whelan, S.P., Guidotti, L.G., and von Andrian, U.H. (2010). Subcapsular sinus macrophages prevent CNS invasion on peripheral infection with a neurotropic virus. Nature 465, 1079–1083.

59. Iftakhar, E.K.I., Fair-Makela, R., Kukkonen-Macchi, A., Elima, K., Karikoski, M., Rantakari, P., Miyasaka, M., Salmi, M., and Jalkanen, S. (2016). Gene-expression profiling of different arms of lymphatic vasculature identifies candidates for manipulation of cell traffic. Proc Natl Acad Sci U S A 113, 10643–10648.

60. Inoshima, I., Inoshima, N., Wilke, G.A., Powers, M.E., Frank, K.M., Wang, Y., and Bubeck Wardenburg, J. (2011). A Staphylococcus aureus pore-forming toxin subverts the activity of ADAM10 to cause lethal infection in mice. Nat Med 17, 1310–1314.

61. Karrer, U., Althage, A., Odermatt, B., Roberts, C.W., Korsmeyer, S.J., Miyawaki, S., Hengartner, H., and Zinkernagel, R.M. (1997). On the key role of secondary lymphoid organs in antiviral immune responses studied in alymphoplastic (aly/aly) and spleenless (Hox11(-)/-) mutant mice. J Exp Med 185, 2157–2170.

62. Kashem, S.W., Riedl, M.S., Yao, C., Honda, C.N., Vulchanova, L., and Kaplan, D.H. (2015). Nociceptive Sensory Fibers Drive Interleukin-23 Production from CD301b+ Dermal Dendritic Cells and Drive Protective Cutaneous Immunity. Immunity 43, 515–526.

63. Kawashima, H., Petryniak, B., Hiraoka, N., Mitoma, J., Huckaby, V., Nakayama, J., Uchimura, K., Kadomatsu, K., Muramatsu, T., Lowe, J.B., et al. (2005). N-acetylglucosamine-6-O-sulfotransferases 1 and 2 cooperatively control lymphocyte homing through L-selectin ligand biosynthesis in high endothelial venules. Nat Immunol 6, 1096–1104.

64. Kharchenko, P.V., Silberstein, L., and Scadden, D.T. (2014). Bayesian approach to single-cell differential expression analysis. Nat Methods 11, 740–742.

65. Kim, C.K., Adhikari, A., and Deisseroth, K. (2017). Integration of optogenetics with complementary methodologies in systems neuroscience. Nat Rev Neurosci 18, 222–235.

66. Klein, R.S., Garber, C., and Howard, N. (2017). Infectious immunity in the central nervous system and brain function. Nat Immunol 18, 132–141.

67. Kuehn, E.D., Meltzer, S., Abraira, V.E., Ho, C.Y., and Ginty, D.D. (2019). Tiling and somatotopic alignment of mammalian low-threshold mechanoreceptors. Proc Natl Acad Sci U S A 116, 9168–9177.

68. Kulkarni, R.M., Greenberg, J.M., and Akeson, A.L. (2009). NFATc1 regulates lymphatic endothelial development. Mech Dev 126, 350–365.

69. Kumar, M.P., Du, J., Lagoudas, G., Jiao, Y., Sawyer, A., Drummond, D.C., Lauffenburger, D.A., and Raue, A. (2018). Analysis of Single-Cell RNA-Seq Identifies Cell-Cell Communication Associated with Tumor Characteristics. Cell Rep 25, 1458–1468 e1454.

70. Kupari, J., Haring, M., Agirre, E., Castelo-Branco, G., and Ernfors, P. (2019). An Atlas of Vagal Sensory Neurons and Their Molecular Specialization. Cell Rep 27, 2508–2523 e2504.

71. Lakkis, F.G., Arakelov, A., Konieczny, B.T., and Inoue, Y. (2000). Immunologic ‘ignorance’ of vascularized organ transplants in the absence of secondary lymphoid tissue. Nat Med 6, 686–688.

72. Larrivee, B., Freitas, C., Suchting, S., Brunet, I., and Eichmann, A. (2009). Guidance of vascular development: lessons from the nervous system. Circ Res 104, 428–441.

73. Lawson, S.N., Crepps, B., and Perl, E.R. (2002). Calcitonin gene-related peptide immunoreactivity and afferent receptive properties of dorsal root ganglion neurones in guinea-pigs. J Physiol 540, 989–1002.

74. Lewin, G.R., and Stucky, C.L. (2000). Sensory neuron mechanotransduction:regulation and underlying molecular mechanisms. In Molecular basisof pain transduction (New York: Wiley), pp. pp 129–148.

75. Li, L., Rutlin, M., Abraira, V.E., Cassidy, C., Kus, L., Gong, S., Jankowski, M.P., Luo, W., Heintz, N., Koerber, H.R., et al. (2011). The functional organization of cutaneous low-threshold mechanosensory neurons. Cell 147, 1615–1627.

76. Lorton, D., Lubahn, C., Engan, C., Schaller, J., Felten, D.L., and Bellinger, D.L. (2000). Local application of capsaicin into the draining lymph nodes attenuates expression of adjuvant-induced arthritis. Neuroimmunomodulation 7, 115–125.

77. Lutter, S., Xie, S., Tatin, F., and Makinen, T. (2012). Smooth muscle-endothelial cell communication activates Reelin signaling and regulates lymphatic vessel formation. J Cell Biol 197, 837–849.

78. Mack, J.J., and Iruela-Arispe, M.L. (2018). NOTCH regulation of the endothelial cell phenotype. Curr Opin Hematol 25, 212–218.

79. Mackenzie, F., and Ruhrberg, C. (2012). Diverse roles for VEGF-A in the nervous system. Development 139, 1371–1380.

80. Macosko, E.Z., Basu, A., Satija, R., Nemesh, J., Shekhar, K., Goldman, M., Tirosh, I., Bialas, A.R., Kamitaki, N., Martersteck, E.M., et al. (2015). Highly Parallel Genome-wide Expression Profiling of Individual Cells Using Nanoliter Droplets. Cell 161, 1202–1214.

81. Malin, S., Molliver, D., Christianson, J.A., Schwartz, E.S., Cornuet, P., Albers, K.M., and Davis, B.M. (2011). TRPV1 and TRPA1 function and modulation are target tissue dependent. J Neurosci 31, 10516–10528.

82. Mao, C., and Obeid, L.M. (2008). Ceramidases: regulators of cellular responses mediated by ceramide, sphingosine, and sphingosine-1-phosphate. Biochim Biophys Acta 1781, 424–434.

83. Mason, M.R., Ehlert, E.M., Eggers, R., Pool, C.W., Hermening, S., Huseinovic, A., Timmermans, E., Blits, B., and Verhaagen, J. (2010). Comparison of AAV serotypes for gene delivery to dorsal root ganglion neurons. Mol Ther 18, 715–724.

84. McCarter, G.C., Reichling, D.B., and Levine, J.D. (1999). Mechanical transduction by rat dorsal root ganglion neurons in vitro. Neurosci Lett 273, 179–182.

85. McCormack, D.G., Mak, J.C., Coupe, M.O., and Barnes, P.J. (1989). Calcitonin gene-related peptide vasodilation of human pulmonary vessels. J Appl Physiol (1985) 67, 1265–1270.

86. McDavid, A., Finak, G., Chattopadyay, P.K., Dominguez, M., Lamoreaux, L., Ma, S.S., Roederer, M., and Gottardo, R. (2013). Data exploration, quality control and testing in single-cell qPCR-based gene expression experiments. Bioinformatics 29, 461–467.

87. McLatchie, L.M., Fraser, N.J., Main, M.J., Wise, A., Brown, J., Thompson, N., Solari, R., Lee, M.G., and Foord, S.M. (1998). RAMPs regulate the transport and ligand specificity of the calcitonin-receptor-like receptor. Nature 393, 333–339.

88. McMahon, S.B., La Russa, F., and Bennett, D.L. (2015). Crosstalk between the nociceptive and immune systems in host defence and disease. Nat Rev Neurosci 16, 389–402.

89. Mempel, T.R., Henrickson, S.E., and Von Andrian, U.H. (2004). T-cell priming by dendritic cells in lymph nodes occurs in three distinct phases. Nature 427, 154–159.

90. Meseguer, V., Alpizar, Y.A., Luis, E., Tajada, S., Denlinger, B., Fajardo, O., Manenschijn, J.A., Fernandez-Pena, C., Talavera, A., Kichko, T., et al. (2014). TRPA1 channels mediate acute neurogenic inflammation and pain produced by bacterial endotoxins. Nat Commun 5, 3125.

91. Mickle, A.D., Won, S.M., Noh, K.N., Yoon, J., Meacham, K.W., Xue, Y., McIlvried, L.A., Copits, B.A., Samineni, V.K., Crawford, K.E., et al. (2019). A wireless closed-loop system for optogenetic peripheral neuromodulation. Nature 565, 361–365.

92. Mithal, D.S., Banisadr, G., and Miller, R.J. (2012). CXCL12 signaling in the development of the nervous system. J Neuroimmune Pharmacol 7, 820–834.

93. Moore, T.C., Lami, J.L., and Spruck, C.H. (1989). Substance P increases lymphocyte traffic and lymph flow through peripheral lymph nodes of sheep. Immunology 67, 109–114.

94. Mooster, J.L., Le Bras, S., Massaad, M.J., Jabara, H., Yoon, J., Galand, C., Heesters, B.A., Burton, O.T., Mattoo, H., Manis, J., et al. (2015). Defective lymphoid organogenesis underlies the immune deficiency caused by a heterozygous S32I mutation in IkappaBalpha. J Exp Med 212, 185–202.

95. Murray, K., Barboza, M., Rude, K.M., Brust-Mascher, I., and Reardon, C. (2019). Functional circuitry of neuro-immune communication in the mesenteric lymph node and spleen. Brain Behav Immun.

96. Nakai, A., Hayano, Y., Furuta, F., Noda, M., and Suzuki, K. (2014). Control of lymphocyte egress from lymph nodes through beta2-adrenergic receptors. J Exp Med 211, 2583–2598.

97. Nassar, M.A., Stirling, L.C., Forlani, G., Baker, M.D., Matthews, E.A., Dickenson, A.H., and Wood, J.N. (2004). Nociceptor-specific gene deletion reveals a major role for Nav1.7 (PN1) in acute and inflammatory pain. Proc Natl Acad Sci U S A 101, 12706–12711.

98. Nonomura, K., Woo, S.H., Chang, R.B., Gillich, A., Qiu, Z., Francisco, A.G., Ranade, S.S., Liberles, S.D., and Patapoutian, A. (2017). Piezo2 senses airway stretch and mediates lung inflation-induced apnoea. Nature 541, 176–181.

99. Norris, G.T., and Kipnis, J. (2019). Immune cells and CNS physiology: Microglia and beyond. J Exp Med 216, 60–70.

100. Oaklander, A.L., and Siegel, S.M. (2005). Cutaneous innervation: form and function. J Am Acad Dermatol 53, 1027–1037.

101. Ordovas-Montanes, J., Rakoff-Nahoum, S., Huang, S., Riol-Blanco, L., Barreiro, O., and von Andrian, U.H. (2015). The Regulation of Immunological Processes by Peripheral Neurons in Homeostasis and Disease. Trends Immunol 36, 578–604.

102. Palazzo, E., Marconi, A., Truzzi, F., Dallaglio, K., Petrachi, T., Humbert, P., Schnebert, S., Perrier, E., Dumas, M., and Pincelli, C. (2012). Role of neurotrophins on dermal fibroblast survival and differentiation. J Cell Physiol 227, 1017–1025.

103. Pappu, R., Schwab, S.R., Cornelissen, I., Pereira, J.P., Regard, J.B., Xu, Y., Camerer, E., Zheng, Y.W., Huang, Y., Cyster, J.G., et al. (2007). Promotion of lymphocyte egress into blood and lymph by distinct sources of sphingosine-1-phosphate. Science 316, 295–298.

104. Pham, T.H., Baluk, P., Xu, Y., Grigorova, I., Bankovich, A.J., Pappu, R., Coughlin, S.R., McDonald, D.M., Schwab, S.R., and Cyster, J.G. (2010). Lymphatic endothelial cell sphingosine kinase activity is required for lymphocyte egress and lymphatic patterning. J Exp Med 207, 17–27.

105. Picelli, S., Faridani, O.R., Bjorklund, A.K., Winberg, G., Sagasser, S., and Sandberg, R. (2014). Full-length RNA-seq from single cells using Smart-seq2. Nat Protoc 9, 171–181.

106. Pinho-Ribeiro, F.A., Baddal, B., Haarsma, R., O’Seaghdha, M., Yang, N.J., Blake, K.J., Portley, M., Verri, W.A., Dale, J.B., Wessels, M.R., et al. (2018). Blocking Neuronal Signaling to Immune Cells Treats Streptococcal Invasive Infection. Cell 173, 1083–1097 e1022.

107. Prinz, M., and Priller, J. (2017). The role of peripheral immune cells in the CNS in steady state and disease. Nat Neurosci 20, 136–144.

108. Rajendran, P.S., Challis, R.C., Fowlkes, C.C., Hanna, P., Tompkins, J.D., Jordan, M.C., Hiyari, S., Gabris-Weber, B.A., Greenbaum, A., Chan, K.Y., et al. (2019). Identification of peripheral neural circuits that regulate heart rate using optogenetic and viral vector strategies. Nat Commun 10, 1944.

109. Ramilowski, J.A., Goldberg, T., Harshbarger, J., Kloppmann, E., Lizio, M., Satagopam, V.P., Itoh, M., Kawaji, H., Carninci, P., Rost, B., et al. (2015). A draft network of ligand-receptor-mediated multicellular signalling in human. Nat Commun 6, 7866.

110. Ranade, S.S., Woo, S.H., Dubin, A.E., Moshourab, R.A., Wetzel, C., Petrus, M., Mathur, J., Begay, V., Coste, B., Mainquist, J., et al. (2014). Piezo2 is the major transducer of mechanical forces for touch sensation in mice. Nature 516, 121–125.

111. Reardon, C., Duncan, G.S., Brustle, A., Brenner, D., Tusche, M.W., Olofsson, P.S., Rosas-Ballina, M., Tracey, K.J., and Mak, T.W. (2013). Lymphocyte-derived ACh regulates local innate but not adaptive immunity. Proc Natl Acad Sci U S A 110, 1410–1415.

112. Renier, N., Wu, Z., Simon, D.J., Yang, J., Ariel, P., and Tessier-Lavigne, M. (2014). iDISCO: a simple, rapid method to immunolabel large tissue samples for volume imaging. Cell 159, 896–910.

113. Rice, F.L., and Albrecht, P.J. (2008). Cutaneous mechanisms of tactile perception: morphological and chemical organization of the innervation to the skin. (San Diego: Academic Press).

114. Richardson, J.D., and Vasko, M.R. (2002). Cellular mechanisms of neurogenic inflammation. J Pharmacol Exp Ther 302, 839–845.

115. Riol-Blanco, L., Ordovas-Montanes, J., and Perro, M. (2014). Nociceptive sensory neurons drive interleukin-23-mediated psoriasiform skin inflammation. Nature 510.

116. Robertson, B. (1990). Wheat germ agglutinin binding in rat primary sensory neurons: a histochemical study. Histochemistry 94, 81–85.

117. Robinson, D.R., and Gebhart, G.F. (2008). Inside information: the unique features of visceral sensation. Mol Interv 8, 242–253.

118. Rosas-Ballina, M., Olofsson, P.S., Ochani, M., Valdes-Ferrer, S.I., Levine, Y.A., Reardon, C., Tusche, M.W., Pavlov, V.A., Andersson, U., Chavan, S., et al. (2011). Acetylcholine-synthesizing T cells relay neural signals in a vagus nerve circuit. Science 334, 98–101.

119. Saria, A. (1984). Substance P in sensory nerve fibres contributes to the development of oedema in the rat hind paw after thermal injury. Br J Pharmacol 82, 217–222.

120. Shekhar, K., Lapan, S.W., Whitney, I.E., Tran, N.M., Macosko, E.Z., Kowalczyk, M., Adiconis, X., Levin, J.Z., Nemesh, J., Goldman, M., et al. (2016). Comprehensive Classification of Retinal Bipolar Neurons by Single-Cell Transcriptomics. Cell 166, 1308–1323 e1330.

121. Shepherd, A.J., Beresford, L.J., Bell, E.B., and Miyan, J.A. (2005a). Mobilisation of specific T cells from lymph nodes in contact sensitivity requires substance P. J Neuroimmunol 164, 115–123.

122. Shepherd, A.J., Downing, J.E., and Miyan, J.A. (2005b). Without nerves, immunology remains incomplete -in vivo veritas. Immunology 116, 145–163.

123. Shibuya, M. (2011). Vascular Endothelial Growth Factor (VEGF) and Its Receptor (VEGFR) Signaling in Angiogenesis: A Crucial Target for Anti- and Pro-Angiogenic Therapies. Genes Cancer 2, 1097–1105.

124. Smillie, C.S., Biton, M., Ordovas-Montanes, J., Sullivan, K.M., Burgin, G., Graham, D.B., Herbst, R.H., Rogel, N., Slyper, M., Waldman, J., et al. (2019). Intra- and Inter-cellular Rewiring of the Human Colon during Ulcerative Colitis. Cell 178, 714–730 e722.

125. Stein, J.V., Rot, A., Luo, Y., Narasimhaswamy, M., Nakano, H., Gunn, M.D., Matsuzawa, A., Quackenbush, E.J., Dorf, M.E., and von Andrian, U.H. (2000). The CC chemokine thymus-derived chemotactic agent 4 (TCA-4, secondary lymphoid tissue chemokine, 6Ckine, exodus-2) triggers lymphocyte function-associated antigen 1-mediated arrest of rolling T lymphocytes in peripheral lymph node high endothelial venules. J Exp Med 191, 61–76.

126. Sudhof, T.C. (2018). Towards an Understanding of Synapse Formation. Neuron 100, 276–293.

127. Suvas, S. (2017). Role of Substance P Neuropeptide in Inflammation, Wound Healing, and Tissue Homeostasis. J Immunol 199, 1543–1552.

128. Takahashi, Y., and Nakajima, Y. (1996). Dermatomes in the rat limbs as determined by antidromic stimulation of sensory C-fibers in spinal nerves. Pain 67, 197–202.

129. Takeda, A., Hollmen, M., Dermadi, D., Pan, J., Brulois, K.F., Kaukonen, R., Lonnberg, T., Bostrom, P., Koskivuo, I., Irjala, H., et al. (2019). Single-Cell Survey of Human Lymphatics Unveils Marked Endothelial Cell Heterogeneity and Mechanisms of Homing for Neutrophils. Immunity 51, 561–572 e565.

130. Tallini, Y.N., Shui, B., Greene, K.S., Deng, K.Y., Doran, R., Fisher, P.J., Zipfel, W., and Kotlikoff, M.I. (2006). BAC transgenic mice express enhanced green fluorescent protein in central and peripheral cholinergic neurons. Physiol Genomics 27, 391–397.

131. Tracey, K.J. (2007). Physiology and immunology of the cholinergic antiinflammatory pathway. J Clin Invest 117, 289–296.

132. Trombetta, J.J., Gennert, D., Lu, D., Satija, R., Shalek, A.K., and Regev, A. (2014). Preparation of Single-Cell RNA-Seq Libraries for Next Generation Sequencing. Curr Protoc Mol Biol 107, 4 22 21-17.

133. Uchimura, K., Gauguet, J.M., Singer, M.S., Tsay, D., Kannagi, R., Muramatsu, T., von Andrian, U.H., and Rosen, S.D. (2005). A major class of L-selectin ligands is eliminated in mice deficient in two sulfotransferases expressed in high endothelial venules. Nat Immunol 6, 1105–1113.

134. Ulvmar, M.H., Werth, K., Braun, A., Kelay, P., Hub, E., Eller, K., Chan, L., Lucas, B., Novitzky-Basso, I., Nakamura, K., et al. (2014). The atypical chemokine receptor CCRL1 shapes functional CCL21 gradients in lymph nodes. Nat Immunol 15, 623–630.

135. Usoskin, D., Furlan, A., Islam, S., Abdo, H., Lonnerberg, P., Lou, D., Hjerling-Leffler, J., Haeggstrom, J., Kharchenko, O., Kharchenko, P.V., et al. (2015). Unbiased classification of sensory neuron types by large-scale single-cell RNA sequencing. Nat Neurosci 18, 145–153.

136. Vaahtomeri, K., Karaman, S., Makinen, T., and Alitalo, K. (2017). Lymphangiogenesis guidance by paracrine and pericellular factors. Genes Dev 31, 1615–1634.

137. Vanlandewijck, M., He, L., Mae, M.A., Andrae, J., Ando, K., Del Gaudio, F., Nahar, K., Lebouvier, T., Lavina, B., Gouveia, L., et al. (2018). A molecular atlas of cell types and zonation in the brain vasculature. Nature 554, 475–480.

138. Veiga-Fernandes, H., and Mucida, D. (2016). Neuro-Immune Interactions at Barrier Surfaces. Cell 165, 801–811.

139. Vento-Tormo, R., Efremova, M., Botting, R.A., Turco, M.Y., Vento-Tormo, M., Meyer, K.B., Park, J.E., Stephenson, E., Polanski, K., Goncalves, A., et al. (2018). Single-cell reconstruction of the early maternal-fetal interface in humans. Nature 563, 347–353.

140. von Andrian, U.H. (1996). Intravital microscopy of the peripheral lymph node microcirculation in mice. Microcirculation 3, 287–300.

141. von Andrian, U.H., and Mempel, T.R. (2003). Homing and cellular traffic in lymph nodes. Nat Rev Immunol 3, 867–878.

142. Wacker, M.J., Tehrani, R.N., Smoot, R.L., and Orr, J.A. (2002). Thromboxane A(2) mimetic evokes a bradycardia mediated by stimulation of cardiac vagal afferent nerves. Am J Physiol Heart Circ Physiol 282, H482–490.

143. Waltman, L., and Van Eck, N.J. (2013). A smart local moving algorithm for large-scale modularity-based community detection. Eur Phys J B 86.

144. Wilke, G.A., and Bubeck Wardenburg, J. (2010). Role of a disintegrin and metalloprotease 10 in Staphylococcus aureus alpha-hemolysin-mediated cellular injury. Proc Natl Acad Sci U S A 107, 13473–13478.

145. Williams, E.K., Chang, R.B., Strochlic, D.E., Umans, B.D., Lowell, B.B., and Liberles, S.D. (2016). Sensory Neurons that Detect Stretch and Nutrients in the Digestive System. Cell 166, 209–221.

146. Winkler, C., and Yao, S. (2014). The midkine family of growth factors: diverse roles in nervous system formation and maintenance. Br J Pharmacol 171, 905–912.

147. Woo, S.H., Lukacs, V., de Nooij, J.C., Zaytseva, D., Criddle, C.R., Francisco, A., Jessell, T.M., Wilkinson, K.A., and Patapoutian, A. (2015). Piezo2 is the principal mechanotransduction channel for proprioception. Nat Neurosci 18, 1756–1762.

148. Wood, J.N., Emery, E.C., and Ernfors, P. (2018). Dorsal Root Ganglion Neuron Types and Their Functional Specialization (Oxford University Press).

149. Wu, H., Xiong, W.C., and Mei, L. (2010). To build a synapse: signaling pathways in neuromuscular junction assembly. Development 137, 1017–1033.

150. Wu, M., Du, Y., Liu, Y., He, Y., Yang, C., Wang, W., and Gao, F. (2014). Low molecular weight hyaluronan induces lymphangiogenesis through LYVE-1-mediated signaling pathways. PLoS One 9, e92857.

151. Yamano, T., Dobes, J., Voboril, M., Steinert, M., Brabec, T., Zietara, N., Dobesova, M., Ohnmacht, C., Laan, M., Peterson, P., et al. (2019). Aire-expressing ILC3-like cells in the lymph node display potent APC features. J Exp Med 216, 1027–1037.

152. Yang, F.C., Tan, T., Huang, T., Christianson, J., Samad, O.A., Liu, Y., Roberson, D., Davis, B.M., and Ma, Q. (2013). Genetic control of the segregation of pain-related sensory neurons innervating the cutaneous versus deep tissues. Cell Rep 5, 1353–1364.

153. Yang, X.M., Han, H.X., Sui, F., Dai, Y.M., Chen, M., and Geng, J.G. (2010). Slit-Robo signaling mediates lymphangiogenesis and promotes tumor lymphatic metastasis. Biochem Biophys Res Commun 396, 571–577.

154. Ye, X., Wang, Y., and Nathans, J. (2010). The Norrin/Frizzled4 signaling pathway in retinal vascular development and disease. Trends Mol Med 16, 417–425.

155. Zeng, W., Pirzgalska, R.M., Pereira, M.M., Kubasova, N., Barateiro, A., Seixas, E., Lu, Y.H., Kozlova, A., Voss, H., Martins, G.G., et al. (2015). Sympathetic neuro-adipose connections mediate leptin-driven lipolysis. Cell 163, 84–94.

156. Zeng, W.Z., Marshall, K.L., Min, S., Daou, I., Chapleau, M.W., Abboud, F.M., Liberles, S.D., and Patapoutian, A. (2018). PIEZOs mediate neuronal sensing of blood pressure and the baroreceptor reflex. Science 362, 464–467.

157. Zhang, G., Brady, J., Liang, W.C., Wu, Y., Henkemeyer, M., and Yan, M. (2015). EphB4 forward signalling regulates lymphatic valve development. Nat Commun 6, 6625.

158. Zhou, P., Hwang, K.W., Palucki, D., Kim, O., Newell, K.A., Fu, Y.X., and Alegre, M.L. (2003). Secondary lymphoid organs are important but not absolutely required for allograft responses. Am J Transplant 3, 259–266.

