## Supplemental Figures for "Sensory Neurons Innervate Peripheral Lymph Nodes and Locally Regulate Gene Expression in Postsynaptic Endothelium, Stromal Cells, and Innate Leukocytes"

#### SUPPLEMENTARY FIGURE LEGENDS

**Figure S1. Confirmation of specific labeling of sensory and sympathetic innervation of LN with Nav1.8$^{Cre}$ and TH**

*Related to Figure 1*

**A.** 3D reconstruction of a representative confocal image of whole-mount popLNs from *ChAT$^{BAC}$-eGFP* animals, where cholinergic cells are genetically labeled with GFP, stained for β3-Tubulin (red), GFP (green). **B.** 3D reconstruction of a representative confocal image of whole-mount iLNs from tamoxifen-treated *Bmx-CreER$^{T2}$ Rosa26$^{LSL\text{-}eYFP/+}$* animals, where arterial endothelial cells are genetically labeled with YFP, stained for β3-Tubulin (red), YFP (green), and smooth muscle actin (SMA, blue), demonstrating preferential association between nerves and arterial vessels supplying LNs. **C.** The schematic for fluorescent wheat-germ agglutinin (WGA) -based retrograde tracing of iLN-innervating sensory neurons. **D** and **E.** Representative confocal images of sections from DRGs and SGs containing WGA-AF488$^{+}$ retrogradely-labeled iLN-innervating neurons of *Nav1.8$^{Cre/+}$; Rosa26$^{LSL\text{-}tdTomato/+}$* animals with endogenous fluorescence from tdTomato and WGA-AF488 shown in red and green, respectively. **F** and **G.** Representative confocal images of sections from DRGs and SGs containing WGA-AF488$^{+}$ retrogradely-labeled iLN-innervating neurons of *Nav1.8$^{Cre/+}$; Rosa26$^{LSL\text{-}tdTomato/+}$* animals, with TH stained in red and endogenous fluorescence from WGA-AF488 shown in green. **H.** Quantification of the percentage of WGA-AF488$^{+}$ retrogradely-labeled LN-innervating DRG or SG neurons that are genetically labeled with Nav1.8$^{Cre}$ or are TH$^{+}$**.** Significance assessed by Welch's t-test, p=0.0003 (***), p<0.001 (***), based on a total of 60 WGA-AF88$^{+}$ DRG neurons and 167 WGA-AF88$^{+}$ SG neurons from 3 mice.

**Figure S2. Anatomical characterization of sensory innervation of different LN subdomains**

*Related to Figure 2*

**A.** A representative confocal section of whole-mount iLNs from *Nav1.8$^{Cre/+}$*; *Rosa26$^{tdTomato/+}$; Prox1-EGFP* animals, stained for tdTomato (red), GFP (green), CD45 (blue) and CD169 (white), allowing identification of LN subdomains, including T cell zone labeled as (T), B follicles labeled as (B) and the medulla labeled as (M). Note that the outermost layer of GFP$^{+}$ LECs (green) delineates LN boundary. **B**. A representative confocal section of whole-mount iLNs from *Nav1.8$^{Cre/+}$; Rosa26$^{LSL\text{-}tdTomato/LSL\text{-}tdTomato}$* animals, stained for tdTomato (red), PNAd (green) and SMA (blue), demonstrating a general lack of contact between tdTomato$^{+}$ sensory fibers (arrowhead) and PNAd$^{+}$ high endothelial venules (HEVs). **C.** 3D view of a representative confocal substack of whole-mount iLNs from *Nav1.8$^{Cre/+}$; Rosa26$^{LSL\text{-}tdTomato/LSL\text{-}tdTomato}$* animals, stained for tdTomato (red) and CD31 (green), illustrating the perivascular origin of capsular/subcapsular fibers (arrowhead). **D** and **E.** Section view of an *intravital* two-photon micrograph of representative capsular/subcapsular tdTomato$^{+}$ sensory fibers (red) in relation to *in vivo* antibody labeled CD169$^{+}$ SCS macrophages (green) and collagen fibers (blue) detected by second-harmonic generation microscopy in popliteal LNs from *Nav1.8*$^{Cre/+}$*; Rosa26*$^{LSL\text{-}tdTomato/+}$ animals, demonstrating the presence of capsular (**D**) and subcapsular (**E**) fibers.

**Figure S3. Evaluation of the specificity and efficiency of the viral-based retrograde labeling**

*Related to Figure 3*

**A.** 3D reconstruction of a representative confocal image of whole-mount iLNs targeted for intranodal injection with AAV-Cre from *Rosa26*$^{LSL\text{-}tdTomato/LSL\text{-}tdTomato}$ animals, stained for tdTomato (red) to show the extent of labeling at the site of injection. **B.** Representative epifluorescence image of tdTomato$^{+}$ DRG neurons retrogradely labeled from perinodal injection in a whole-mount spinal cord-DRG preparation without antibody amplification. **C.** Quantification of the number of retrograde-labeled neurons from intranodal and perinodal injection. Intranodal: 16.25 (mean) ± 2.394 (SEM); perinodal: 1.500 (mean) ± 0.6455 (SEM), p=0.0064 (**) by Welch's t-test, n=4 per group. **D.** 3D reconstruction of a representative confocal image of whole-mount iLNs targeted for intranodal injection with AAV-Flex-tdTomato from *Nav1.8*$^{Cre/+}$*; Rosa26*$^{LSL\text{-}eYFP/+}$ animals, stained for tdTomato (red) and CD31 (green) to visualize the axonal projections of retrogradely-labeled neurons. **E.** Representative epifluorescence image of tdTomato$^{+}$ DRG neurons retrogradely labeled from the skin overlying the iLN in a whole-mount spinal cord-DRG preparation without antibody amplification. **F.** 3D reconstruction of a representative confocal image of whole-mount skin targeted for intradermal injection with AAV-Flex-eGFP from *Nav1.8*$^{Cre/+}$ animals, stained for GFP (green) and DAPI (blue) to demonstrate the specificity of labeling from the skin. **G.** Soma size distribution of retrogradely-labeled iLN-innervating and randomly sampled CGRP$^{+}$ neurons from the same DRGs following intranodal injection of AAV-Cre into *Rosa26*$^{LSL\text{-}tdTomato/LSL\text{-}tdTomato}$ animals based on 49 tdTomato$^{+}$ iLN-innervating and 114 CGRP$^{+}$ DRG neurons from 3 and 2 mice, respectively. **H.** 3D reconstruction of a representative confocal image of whole-mount iLNs from *Nav1.8*$^{Cre/+}$*; Rosa26*$^{LSL\text{-}tdTomato/LSL\text{-}tdTomato}$ animals, stained for tdTomato (red), NFH (green) and TH (blue) to visualize co-innervation of LNs by myelinated and unmyelinated sensory fibers.

**Figure S4. Expression of neuronal functional pathways by LN- and skin-innervating sensory neurons and evaluation of specificity and sensitivity of select genes as markers of LN-innervating sensory neurons**

*Related to Figure 5*

**A.** and **B.** Expression of Prokr2 (**A**) and Ptgir (**B**) by innervation target and Neuron Type. Significance assessed by Mann-Whitney-Wilcoxon test * p<0.05, *** p<0.001. **C**-**E** Heatmaps of gene lists curated in the Usoskin et al. Sensory Neuron Atlas (Usoskin et al., 2015). Blue bar represents skin-innervating neurons, yellow bar represents LN-innervating neurons. **F-I**. Representative confocal images of RNAscope analysis for indicated target (green) on DRG sections containing tdTomato+ retrogradely-labeled neurons (red) (**F, H**: LN-innervating, **G, I**: skin-innervating).

**Figure S5. A single-cell transcriptomic atlas of inguinal LNs (iLN) at steady state**

*Related to Figure 6*

**A.** tSNE of 9,662 single cells from an initial clustering analysis. * indicates clusters that required additional subclustering to resolve all constituent cell types. **B.** tSNE of T cell parent cluster. **C**. Dot plot of the top DE genes between Tregs and remaining non-Treg T cells. **D**. tSNE of additional subclustering of non-Treg T cells from **B**. **E.** Dot plot of the top DE genes between CD4 T cells and CD8 T cells. **F.** tSNE of parent macrophage and dendritic cell cluster. **G**. Dot plot of the top DE genes between Macrophages and cDC 2 cells. **H**. tSNE of parent blood endothelial cell cluster. **I**. Dot plot of the top DE genes between BEC and HEC. **J**. tSNE of parent NK cells. **K**. Dot plot of the top DE genes between mitotic NK cells and non-mitotic NK cells ("NK cells"). **L**. tSNE of parent Neutrophil cluster. **M**. Dot plot of the top DE genes between Neutrophils 1 and Neutrophils 2. **N**. tSNE of parent Lymphatic Endothelial Cell (LEC) cluster. **O.** Dot plot of the top DE genes between LEC 1 and LEC 2. For all dot plots, circle diameters reflect the percent of cells expressing a given marker within that cell type, circle colors reflect relative expression abundance within that cell type; light grey: low, black: high. **P.** Heatmap of gene-normalized, scaled DGE of the top 10 cell type-specific genes for each identified cell type (yellow: high relative expression; purple: low relative expression). Cell types reflect final clustering after subclustering, cell type-specific genes identified by comparing one cell type to all other cells in the dataset. Cell types are subsampled to 25 cells per cell type for heatmap (see **Table S3**).

**Figure S6. Evaluation of methods to determine neuron-LN cell interaction potentials**

*Related to Figure 6*

**A.** Schematic of method for identifying cognate neuron-LN-cell pairs. **B**. Pearson correlation of Interaction Potential (calculated as in **Figure 6G**) with cell quality (measured as median UMI/cell for each cell type). **C** and **D.** Comparison of cell type summary statistics to calculate Interaction Potential (**C**: average expression, **D**: percent expression) * $p<0.05$, ** $p<0.01$, *** $p<0.001$, dashed lines represent 99% confidence interval from permutation test (see **STAR Methods**). **E.** Comparison of Interaction Potential after downsampling to equivalent cell numbers (using the scaled average Interaction Potential, as in **Figure 6G**). **F.** Left: Heatmap of top expressed neuropeptides in LN-innervating sensory neurons; Right: Dotplot of respective cognate receptor expression (denoted by arrows) in LN cells at steady state (circle diameter reflects the percent of cells expressing a given marker within that cell type, circle color reflects relative expression abundance within that cell type; light grey: low, black: high). **G.** and **H.** Top candidate interaction molecules driving high Interaction Potential in **G.** Non-Endothelial Stromal, **H.** BEC**.**

**Figure S7. scRNA-seq of iLNs following optogenetic neuronal stimulation**

*Related to Figure 7*

**A.** 3D reconstruction of representative confocal sections of whole-mount iLNs following intranodal injection of AAV-Flex-tdTomato into *Nav1.8*$^{Cre/+}$*; Rosa26*$^{LSL\text{-}eYFP/+}$ animals, stained for tdTomato (red) and CD31 (green) to reveal the axonal trajectory of iLN-innervating neurons in relation to the site of illumination (arrow). **B**. Abundance of light-induced differentially expressed genes (left: downregulated with light stimulation; right: upregulated with light stimulation) among 2 ChR2+ animals, representing preliminary analysis of optogenetic-based neuronal triggering. **C.** Schematic of mouse cohort and study design. **D.** Correlation between the pseudopopulation average gene expression profile of each cell type from the steady state iLN atlas (**Figure 6**, rows) and the light-stimulated iLN atlas (**Figure 7**, columns). **E.** Quantification of alterations in cell composition by photostimulation, represented as fold change in fractional abundance of each cell type between matched light-exposed and light-unexposed sides within each mouse. Blue circles: ChR2+ mice, n=4; black squares: ChR2- mice, n=3. **F.** Cellularity of light-exposed and light-unexposed iLNs from ChR2- and ChR2+ animals at the end of photostimulation. ChR2-: p= 0.2470 (ns); ChR2: p= 0.6755 (ns) by 2-way ANOVA with Sidak's multiple comparisons test; 5 mice per genotype. **G.** Correlation between abundance of DE genes and interaction potential over multiple of DE gene cutoff parameters.

#### SUPPLEMENTARY TABLES

**Table S1**. Metadata-annotated neuron scRNA-Seq data

*Related to Figures 3 and 4*

**Table S2.** Differentially expressed genes by Neuron Type and between LN-innervating and skin-innervating neurons, gene ontology lists.

*Related to Figures 4 and 5*

**Table S3.** Metadata-annotated steady state LN cell atlas UMI vs. cell matrix

*Related to Figure 6*

**Table S4**. Steady state LN cell atlas: Cell type cluster defining genes and subcluster defining genes.

*Related to Figure 6*

**Table S5**. Differentially expressed genes following neuronal stimulation by cell type and genotype.

*Related to Figure 7*

#### SUPPLEMENTARY MOVIES

**Movie 1.** 3D animation of Figure 1A.

**Movie 2.** 3D animation of Figure 1B.

**Movie 3.** 3D animation of Figure 2A.

**Movie 4**. Slice view of Figure 2C.

**Movie 5.** 3D animation of Figure 2E.

**Movie 6.** 3D animation of surface rendering of Figure S2B.

**Movie 7.** 3D animation of Figure 2F.

**Movie 8.** 3D animation of Figure S3D.

Figure S1

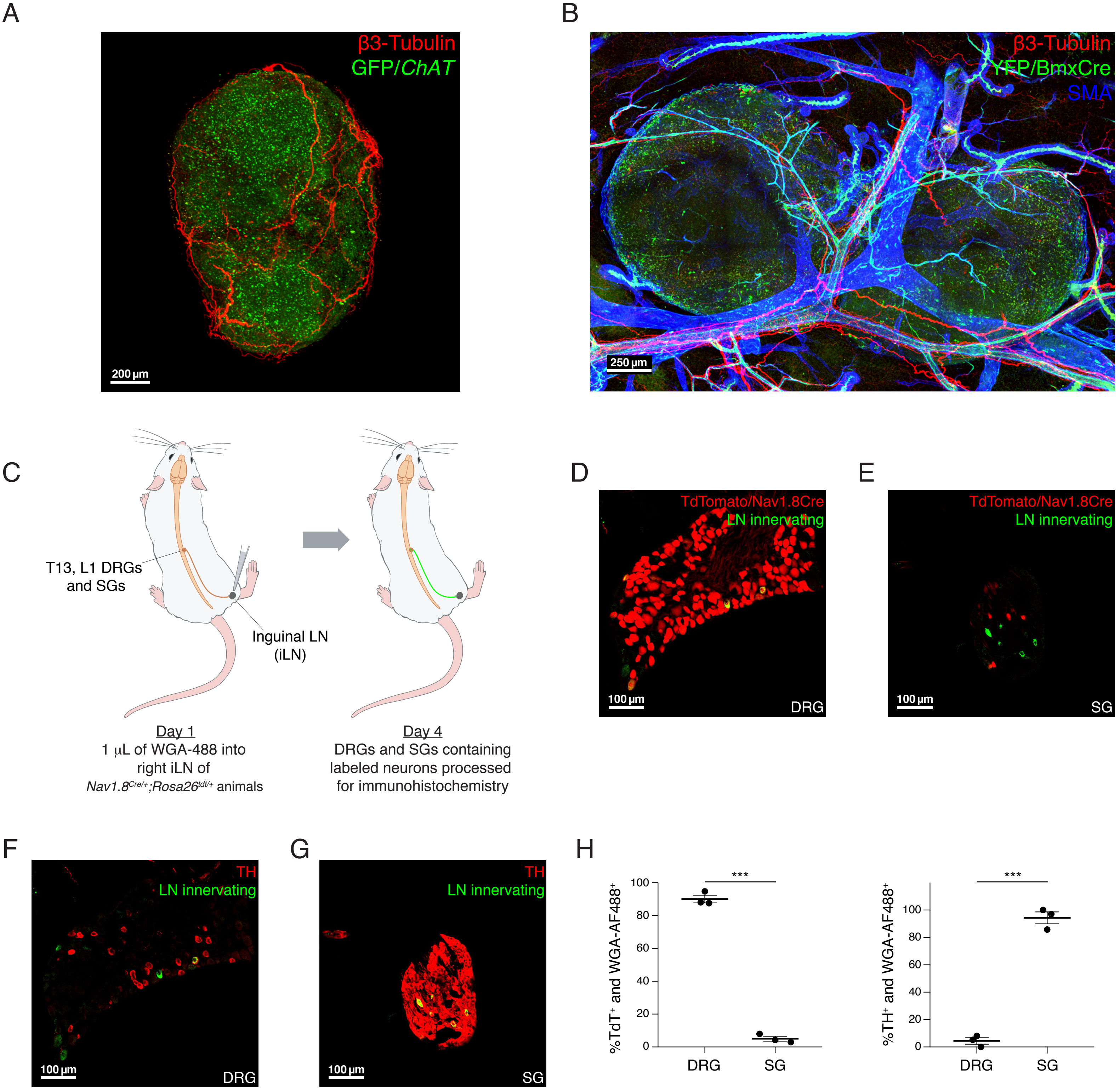
A
β3-Tubulin
GFP/ChAT
200 µm
B
β3-Tubulin
YFP/BmxCre
SMA
250 µm
C
T13, L1 DRGs
and SGs
Inguinal LN
(iLN)
Day 1
1 µL of WGA-488 into
right iLN of
Nav1.8Cre/+;Rosa26tdt/+ animals
Day 4
DRGs and SGs containing
labeled neurons processed
for immunohistochemistry
D
TdTomato/Nav1.8Cre
LN innervating
100 µm
DRG
E
TdTomato/Nav1.8Cre
LN innervating
100 µm
SG
F
TH
LN innervating
100 µm
DRG
G
TH
LN innervating
100 µm
SG
H
%TdT+ and WGA-AF488+
***
0
20
40
60
80
100
DRG
SG
%TH+ and WGA-AF488+
***
0
20
40
60
80
100
DRG
SG

### Figure S2

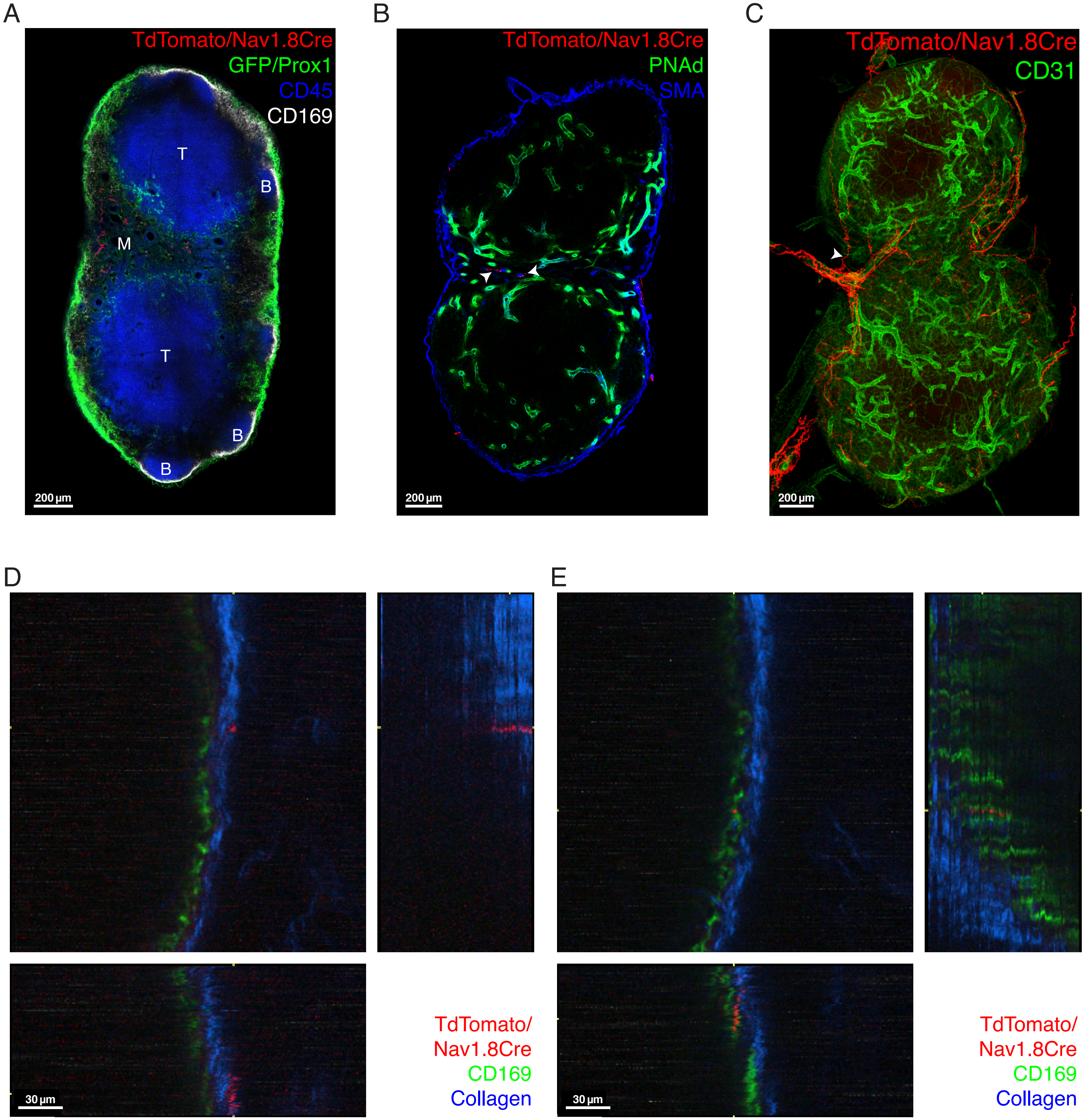
A
TdTomato/Nav1.8Cre
GFP/Prox1
CD45
CD169
T
B
M
T
B
B
200 µm
B
TdTomato/Nav1.8Cre
PNAd
SMA
200 µm
C
TdTomato/Nav1.8Cre
CD31
200 µm
D
30 µm
TdTomato/
Nav1.8Cre
CD169
Collagen
E
30 µm
TdTomato/
Nav1.8Cre
CD169
Collagen

### Figure S3

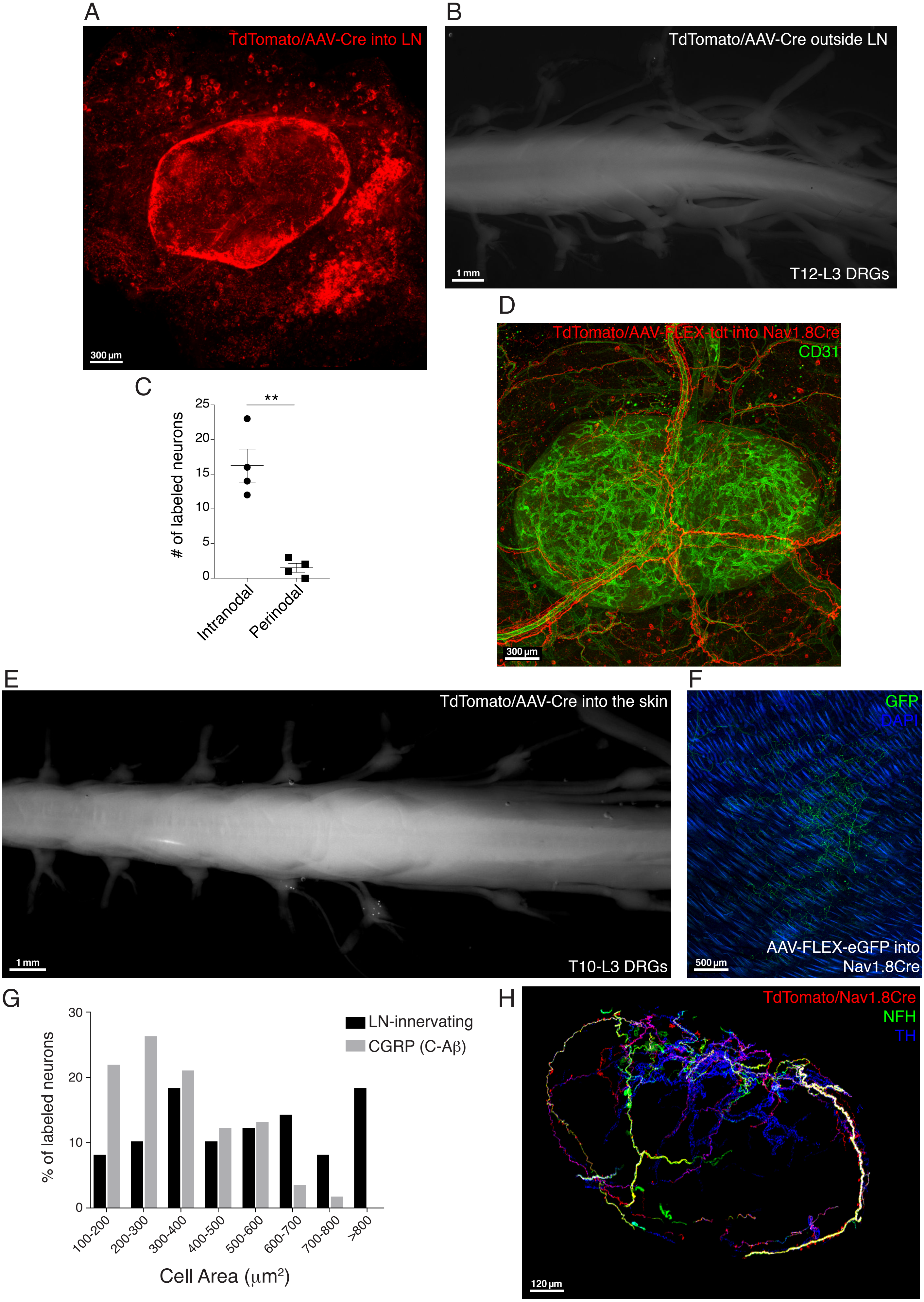
A
TdTomato/AAV-Cre into LN
300 µm
B
TdTomato/AAV-Cre outside LN
1 mm
T12-L3 DRGs
C
### of labeled neurons
25
20
15
10
5
0
**
Intranodal
Perinodal
D
TdTomato/AAV-FLEX-tdt into Nav1.8Cre
CD31
300 µm
E
TdTomato/AAV-Cre into the skin
1 mm
T10-L3 DRGs
F
GFP
DAPI
AAV-FLEX-eGFP into
Nav1.8Cre
500 µm
G
% of labeled neurons
30
20
10
0
LN-innervating
CGRP (C-Aβ)
100-200
200-300
300-400
400-500
500-600
600-700
700-800
>800
Cell Area (µm2)
H
TdTomato/Nav1.8Cre
NFH
TH
120 µm

### Figure S4

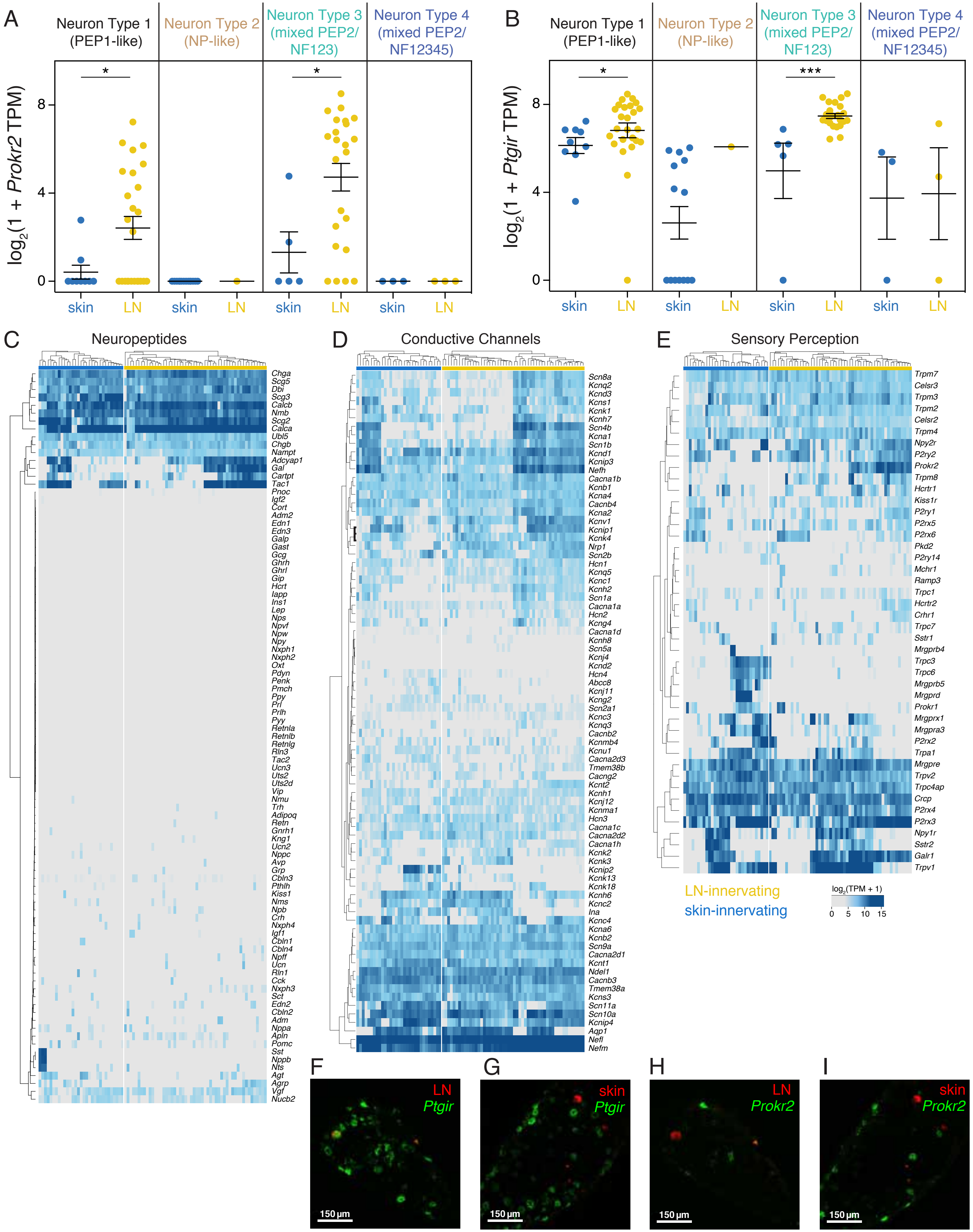
A
Neuron Type 1 (PEP1-like)
Neuron Type 2 (NP-like)
Neuron Type 3 (mixed PEP2/ NF123)
Neuron Type 4 (mixed PEP2/ NF12345)
log2(1 + Prokr2 TPM)
skin
LN
B
log2(1 + Ptgir TPM)
C
Neuropeptides
D
Conductive Channels
E
Sensory Perception
LN-innervating
skin-innervating
log2(TPM + 1)
0 5 10 15
F
LN
Ptgir
G
skin
Ptgir
H
LN
Prokr2
I
skin
Prokr2
150 µm

### Figure S5

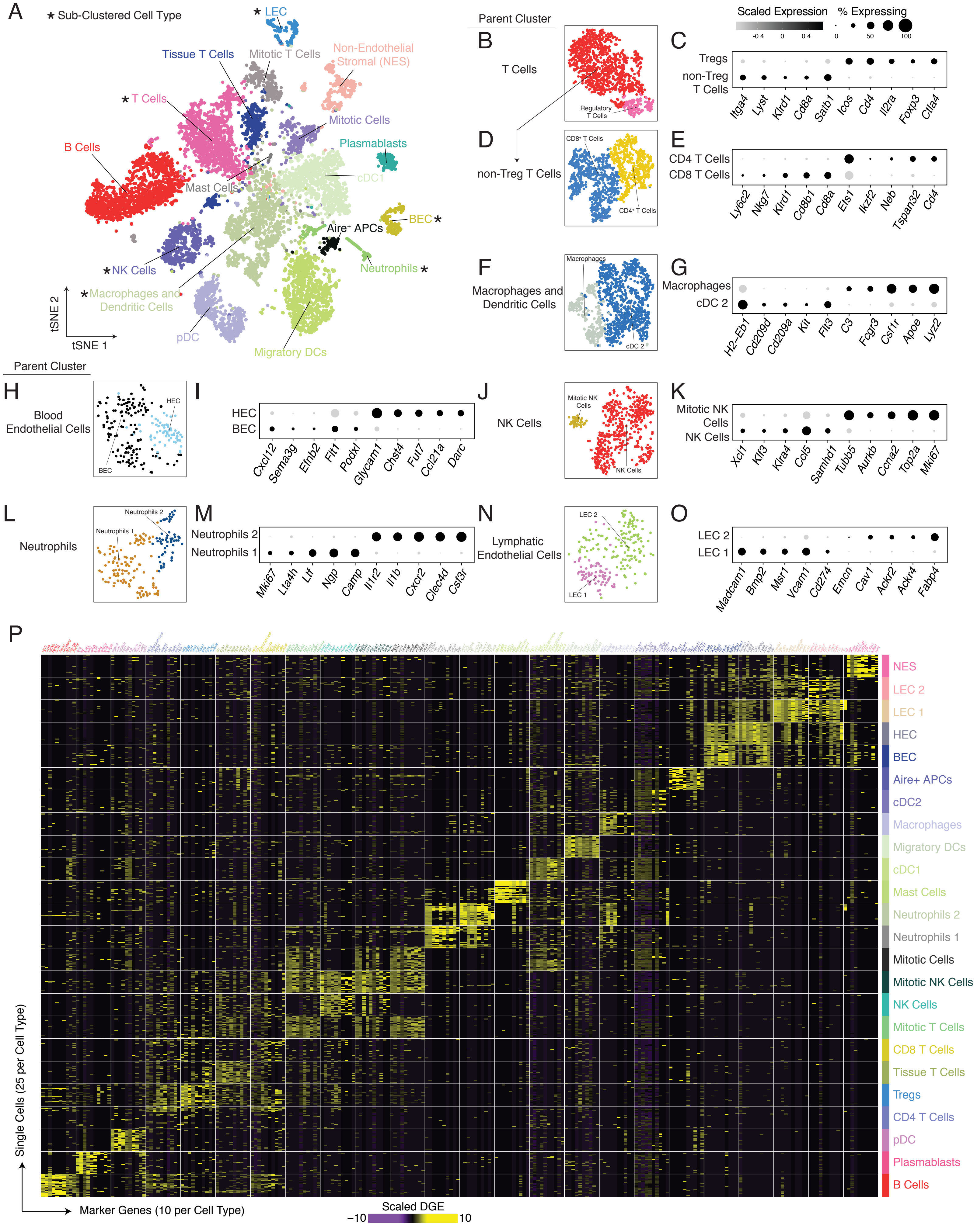
A
* Sub-Clustered Cell Type
* LEC
Tissue T Cells
Mitotic T Cells
Non-Endothelial Stromal (NES)
* T Cells
Mitotic Cells
B Cells
Plasmablasts
Mast Cells
cDC1
BEC *
Aire+ APCs
Neutrophils *
* NK Cells
* Macrophages and Dendritic Cells
pDC
Migratory DCs
tSNE 2
tSNE 1
Parent Cluster
B
T Cells
Regulatory T Cells
C
Scaled Expression
% Expressing
-0.4
0
0.4
0
50
100
Tregs
non-Treg T Cells
Itga4
Lyst
Klrd1
Cd8a
Satb1
Icos
Cd4
Il2ra
Foxp3
Ctla4
D
non-Treg T Cells
CD8+ T Cells
CD4+ T Cells
E
CD4 T Cells
CD8 T Cells
Ly6c2
Nkg7
Klrd1
Cd8b1
Cd8a
Ets1
Ikzf2
Neb
Tspan32
Cd4
F
Macrophages and Dendritic Cells
Macrophages
cDC 2
G
Macrophages
cDC 2
H2-Eb1
Cd209d
Cd209a
Kit
Flt3
C3
Fcgr3
Csf1r
Apoe
Lyz2
Parent Cluster
H
Blood Endothelial Cells
HEC
BEC
I
HEC
BEC
Cxcl12
Sema3g
Efnb2
Flt1
Podxl
Glycam1
Chst4
Fut7
Ccl21a
Darc
J
NK Cells
Mitotic NK Cells
NK Cells
K
Mitotic NK Cells
NK Cells
Xcl1
Klf3
Klra4
Ccl5
Samhd1
Tubb5
Aurkb
Ccna2
Top2a
Mki67
L
Neutrophils
Neutrophils 2
Neutrophils 1
M
Neutrophils 2
Neutrophils 1
Mki67
Lta4h
Ltf
Ngp
Camp
Il1r2
Il1b
Cxcr2
Clec4d
Csf3r
N
Lymphatic Endothelial Cells
LEC 2
LEC 1
O
LEC 2
LEC 1
Madcam1
Bmp2
Msr1
Vcam1
Cd274
Emcn
Cav1
Ackr2
Ackr4
Fabp4
P
NES
LEC 2
LEC 1
HEC
BEC
Aire+ APCs
cDC2
Macrophages
Migratory DCs
cDC1
Mast Cells
Neutrophils 2
Neutrophils 1
Mitotic Cells
Mitotic NK Cells
NK Cells
Mitotic T Cells
CD8 T Cells
Tissue T Cells
Tregs
CD4 T Cells
pDC
Plasmablasts
B Cells
Single Cells (25 per Cell Type)
Marker Genes (10 per Cell Type)
Scaled DGE
−10
10

### Figure S6

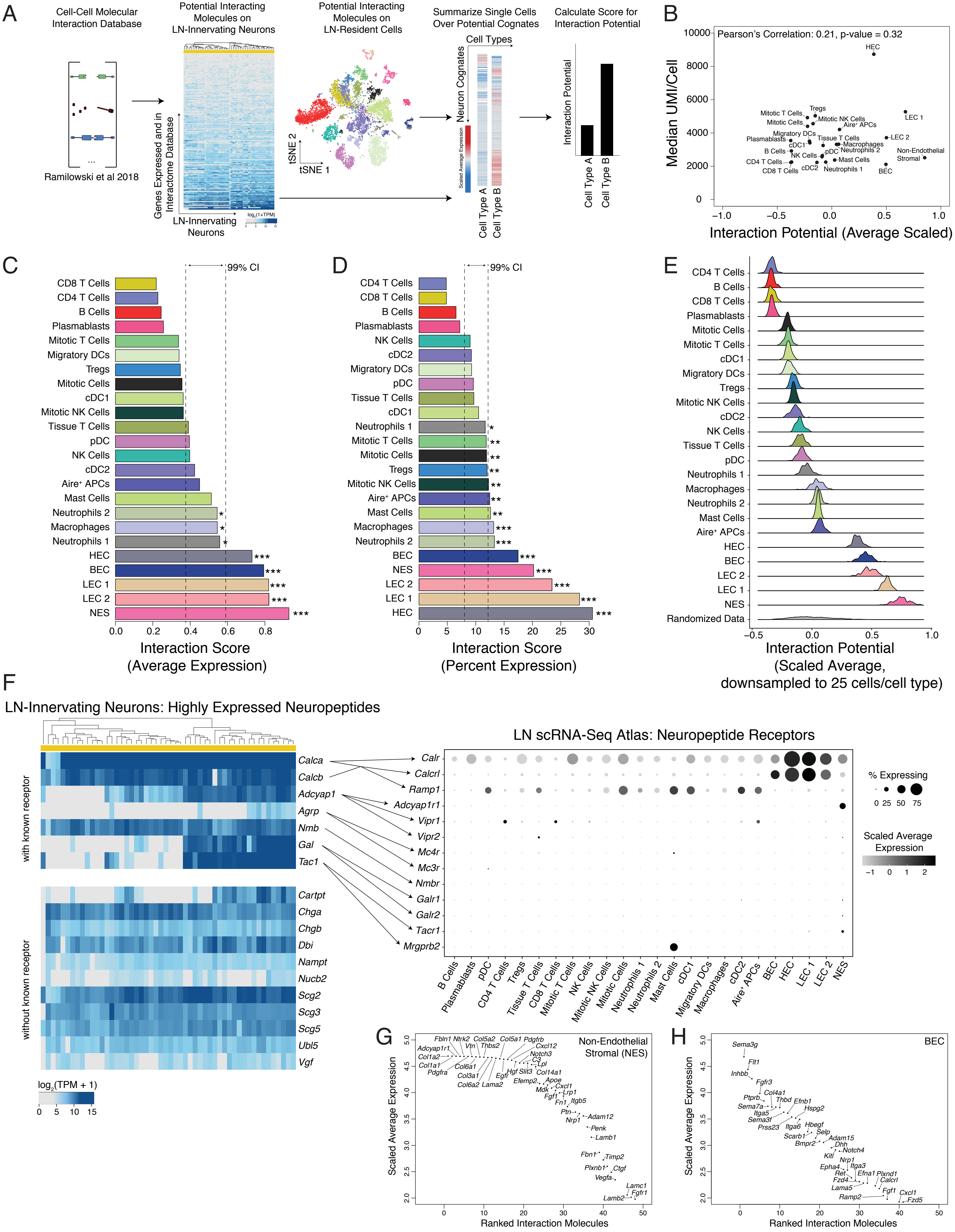

A
Cell-Cell Molecular Interaction Database
Ramilowski et al 2018
Potential Interacting Molecules on LN-Innervating Neurons
Genes Expressed and in Interactome Database
LN-Innervating Neurons
Potential Interacting Molecules on LN-Resident Cells
tSNE 2
tSNE 1
Summarize Single Cells Over Potential Cognates
Cell Types
Neuron Cognates
Cell Type A
Cell Type B
Calculate Score for Interaction Potential
Interaction Potential
B
Pearson's Correlation: 0.21, p-value = 0.32
Median UMI/Cell
Interaction Potential (Average Scaled)
C
99% CI
Interaction Score (Average Expression)
D
99% CI
Interaction Score (Percent Expression)
E
Interaction Potential (Scaled Average, downsampled to 25 cells/cell type)
Randomized Data
F
LN-Innervating Neurons: Highly Expressed Neuropeptides
with known receptor
without known receptor
log2(TPM + 1)
LN scRNA-Seq Atlas: Neuropeptide Receptors
% Expressing
Scaled Average Expression
G
Non-Endothelial Stromal (NES)
Scaled Average Expression
Ranked Interaction Molecules
H
BEC
Scaled Average Expression
Ranked Interaction Molecules

### Figure S7

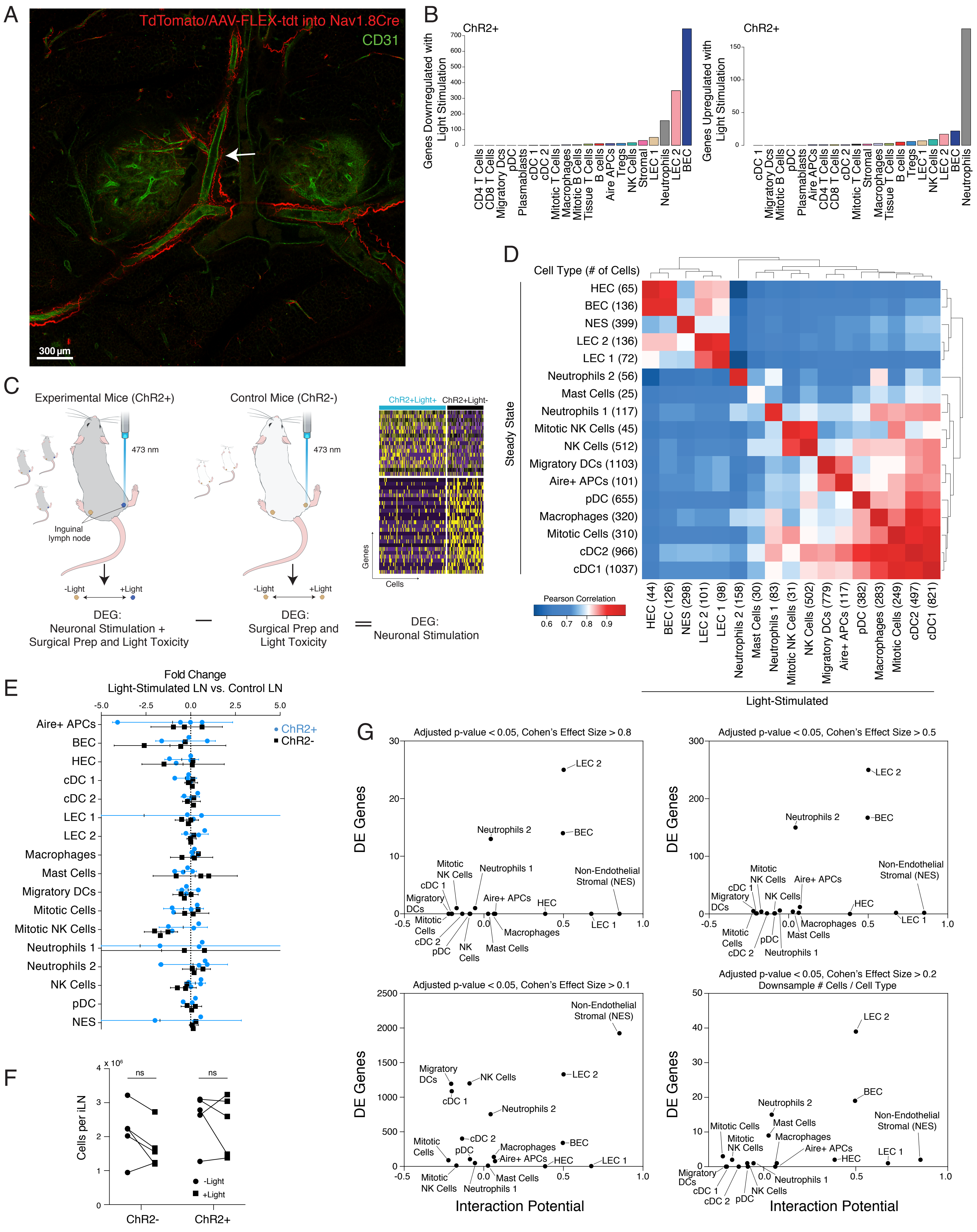

A
TdTomato/AAV-FLEX-tdt into Nav1.8Cre
CD31
300 μm
B
ChR2+
Genes Downregulated with Light Stimulation
CD4 T Cells
CD8 T Cells
Migratory DCs
pDC
Plasmablasts
cDC 1
cDC 2
Mitotic T Cells
Macrophages
Mitotic B Cells
Tissue T Cells
B cells
Aire APCs
Tregs
NK Cells
Stromal
LEC 1
Neutrophils
LEC 2
BEC
ChR2+
Genes Upregulated with Light Stimulation
cDC 1
Migratory Dcs
Mitotic B Cells
pDC
Plasmablasts
Aire APCs
CD4 T Cells
CD8 T Cells
cDC 2
Mitotic T Cells
Stromal
Macrophages
Tissue T Cells
B cells
Tregs
LEC 1
NK Cells
LEC 2
BEC
Neutrophils
C
Experimental Mice (ChR2+)
Control Mice (ChR2-)
473 nm
Inguinal lymph node
-Light
+Light
DEG: Neuronal Stimulation + Surgical Prep and Light Toxicity
DEG: Surgical Prep and Light Toxicity
DEG: Neuronal Stimulation
ChR2+Light+
ChR2+Light-
Genes
Cells
D
Cell Type (# of Cells)
Steady State
HEC (65)
BEC (136)
NES (399)
LEC 2 (136)
LEC 1 (72)
Neutrophils 2 (56)
Mast Cells (25)
Neutrophils 1 (117)
Mitotic NK Cells (45)
NK Cells (512)
Migratory DCs (1103)
Aire+ APCs (101)
pDC (655)
Macrophages (320)
Mitotic Cells (310)
cDC2 (966)
cDC1 (1037)
Pearson Correlation
HEC (44)
BEC (126)
NES (298)
LEC 2 (101)
LEC 1 (98)
Neutrophils 2 (158)
Mast Cells (30)
Neutrophils 1 (83)
Mitotic NK Cells (31)
NK Cells (502)
Migratory DCs (779)
Aire+ APCs (117)
pDC (382)
Macrophages (283)
Mitotic Cells (249)
cDC2 (497)
cDC1 (821)
Light-Stimulated
E
Fold Change
Light-Stimulated LN vs. Control LN
ChR2+
ChR2-
Aire+ APCs
BEC
HEC
cDC 1
cDC 2
LEC 1
LEC 2
Macrophages
Mast Cells
Migratory DCs
Mitotic Cells
Mitotic NK Cells
Neutrophils 1
Neutrophils 2
NK Cells
pDC
NES
F
Cells per iLN
ns
-Light
+Light
ChR2-
ChR2+
G
Adjusted p-value < 0.05, Cohen's Effect Size > 0.8
Adjusted p-value < 0.05, Cohen's Effect Size > 0.5
Adjusted p-value < 0.05, Cohen's Effect Size > 0.1
Adjusted p-value < 0.05, Cohen's Effect Size > 0.2
Downsample # Cells / Cell Type
DE Genes
Interaction Potential
Non-Endothelial Stromal (NES)
